# +A CAR T cell-amplified suppressive B cell circuit in AML

**DOI:** 10.64898/2026.10.12.758644

**Authors:** Adrian Gottschlich, Vladyslav Kavaka, Thomas Alexander Janert, Veronika Igl, Emanuele Carlini, Lisa Rohrbacher, Wendy den Hartog, Jakob Harwood, Marcel P. Trefny, Selin Sogukcelik, Philipp Penovici, Bernhard Ransmayr, Christian Rausch, Franziska Brauneck, Stephanie Frenz-Wiessner, Isabel Goek, Gerulf Hänel, Cristina Zucchinetti, Arman Oner, Paula Sophie Zwicker, Ruth Grünmeier, Simon Delahais, Philipp Maximilian Schnell, Sayantan Nandi, Moritz Thomas, Ruben Brabenec, Daria Briukhovetska, Stefanos Michaelides, Rasmus David Müller, Lilli Ye Ballheimer, Christina Angeliki Tsiverioti, Gordon Victor Hoffmann, Sophia Stock, Ignazio Piseddu, Philipp Jie Müller, Moritz Kruppa, Walter Fiedler, Michael Rade, Markus Kreuz, Niklas Kehl, Anand S Bhagwat, Johannes Tobias Thiel, Dominik Steiner, Henrik Lauer, Adrien Daigeler, Christoph Klein, Carsten Marr, Stefan Endres, Kristin Reiche, Mirco Friedrich, Christine Bollwein, Stephan Ledderose, Martina Rudelius, Tobias Herold, Karsten Spiekermann, Maximilian Merz, Thorsten Mempel, Saar Gill, Marion Subklewe, Michael von Bergwelt-Baildon, Sebastian Kobold

## Abstract

Immunotherapy in acute myeloid leukemia (AML) is constrained by a hostile bone marrow niche. Here we identify atypical memory B cells (B_atyp_) as a suppressive population within this microenvironment. Across single cell and flow cytometry cohorts, B_atyp_ were expanded in AML, and their molecular and phenotypic profiles resembled those of atypical B cells associated with chronic inflammatory conditions. B_atyp_ were polarized within the AML microenvironment, and this was amplified by CAR T cell–AML interactions. Mechanistically, polarization was dependent on Toll-like receptor, interferon and CD70–CD27 signaling. B_atyp_ suppressed T cell and CAR T cell responses via PD-1-and TGF-β-R signaling, and were enriched in non-responding patients with AML and myeloma. CD180-directed CAR T cells, targeting an antigen shared by B_atyp_ and AML blasts, resisted suppression *in vitro* and cleared disease in B cell–humanized xenografts while sparing hematopoietic stem cells, with B cell engagement further enhancing their expansion. CAR T cells thus reinforce their own suppressive niche through B_atyp_ polarization, a constraint that can be relieved by dual blast- and B cell-targeting strategies.

## Introduction

B cell-directed cellular therapies achieved remarkable success in B cell malignancies and established chimeric antigen receptor (CAR) T cells as a curative option for relapsed or refractory disease^1,2^. However, acute myeloid leukemia (AML) remains difficult to treat even with CAR T cell therapy^3^. Despite the identification of numerous candidate antigens^4–9^ the clinical performance of CAR T cells in AML has been inferior to other types of hematological cancers. These underwhelming results include poor efficacy^10^, early relapse^11^, and significant toxicity^12^, largely due to the immunosuppressive tumor microenvironment and on-target–off-tumor recognition of healthy hematopoietic cells^3^. The anti-tumor efficacy of T cell-based immunotherapies in AML is limited by T cell exhaustion, expansion of regulatory myeloid populations, and a suppressive cytokine milieu^10,13^. Despite the general recognition of B cells as important modulators of anti-tumor immunity, the mechanistic contribution of B cells to immune regulation in AML has received little attention^14^.

Atypical memory B cells (B_atyp_) form a B cell population that expands under chronic immune stimulation^15–19^. Their study is complicated by inconsistent nomenclature, with closely related or overlapping populations variously termed age-associated, CD11c , T-bet , double-negative (DN2), or tissue-like/atypical memory B cells depending on context and markers^20^. They exhibit shared transcriptional profiles across malaria, HIV infection, and autoimmunity in cross-disease analyses, suggesting common drivers of their expansion and function^21^. The functional role of these B_atyp_ remains insufficiently understood. They have been most consistently linked to autoantibody production in aging and autoimmunity^22^, especially in diseases such as systemic lupus erythematosus (SLE).

Phenotypically, B_atyp_ are often defined by the combined loss of surface IgD and CD27 (a double-negative phenotype), low or absent expression of CD21 and the germinal-center–homing receptor CXCR5, and high expression of the integrin CD11c (*ITGAX*) and Fc-receptor-like molecules (FCRL3–5)^20^. The double-negative phenotype alone is not sufficient, however, as it also captures DN1 (CXCR5 CD11c ), DN3 (CXCR5 CD11c ) and DN4 (CXCR5 CD11c ) cells that lack the T-bet program^23^.

In humans, several Breg subsets have been described, e.g. CD19 CD24^high^CD38^high^CD1d^high 24,25^ and CD19 CD24^high^CD27 ^26^, whose operational criterion is the suppression of effector T cell responses primarily through IL-10 and to a lesser extent TGF-β^14^. Their developmental relationships remain unresolved. Unlike FOXP3 in regulatory T cells, no single master regulator defines Bregs, whose identity is regarded as a fluid, functionally acquired state^14^. As this state is acquired rather than lineage-fixed, atypical B cells may themselves adopt Breg-like functional features under appropriate conditions, a possibility only beginning to be explored in cancer. Recent work in hepatocellular carcinoma, for example, indicates that atypical memory B cells can acquire regulatory features and dampen T cell co-stimulation^27^.Whether atypical B cells arising in cancer correspond to those described in infection and autoimmunity, and how their functions differ across tumor entities, remains only partially understood.

In AML, single-cell atlases of the bone marrow microenvironment^28,29^, have annotated atypical, inflammation-associated B cell states. There, authors provided evidence for their presence and correlation with inflammation, but deeper phenotypic or functional characterization of these B cell states is missing. Consequently, their developmental drivers, detailed phenotype, functional properties, and -most importantly -their impact on T cell activity and on the efficacy of T cell–based immunotherapies in AML have remained largely undefined. Overall, whether and how B_atyp_ actively contribute to the bone marrow microenvironment, especially in the setting of immunotherapies has not been systematically addressed.

The differentiation and activation of B cells, including atypical subsets, are closely tied to Toll-like receptor signaling. A B cell surface molecule of particular interest in this regard is CD180 (also known as RP105 or LY64), a member of the Toll-like receptor (TLR) family that, unlike canonical TLRs, lacks the intracellular Toll/interleukin-1 receptor (TIR) signaling domain and is expressed at the cell surface in complex with its partner molecule MD-1^30,31^. It is found predominantly on mature B cells, as well as on monocytes, macrophages, and dendritic cells^30^. Physiologically, the RP105/MD-1 complex modulates TLR4-dependent signaling and, on B cells, delivers activating and proliferative signals, thereby regulating B cell activation and antibody responses^32,33^.

In the present study, we characterize B_atyp_ in AML and define their role in shaping anti-leukemic T cell immunity. Combining integrative single-cell and flow cytometric analyses across independent cohorts with functional studies of patient-derived cells, we establish their phenotype and suppressive capacity, and resolve how B_atyp_ are sustained: leukemic blasts polarize B_atyp_ through TLR, interferon and CD70–CD27 signals, CAR T cell activity amplifies these cues, and the resulting B_atyp_ restrain both conventional and CAR T cell responses. Because B_atyp_ and blasts share high-level surface expression of CD180, this circuit can be interrupted at a single target, which we explore using CD180-directed CAR T cells.

## Results

### B_atyp_ accumulate in the AML bone marrow and mirror chronic-inflammatory atypical B cells

Given the emerging relevance of distinct B cell subsets in AML, we analyzed a recently published single-cell RNA sequencing (scRNA-seq) dataset^29^ to comprehensively characterize the composition and transcriptional states of B cells in the AML bone marrow microenvironment.

The dataset comprised 24 bone marrow samples from newly diagnosed AML patients (1^st^ AML), 8 relapsed or refractory AML patients (r/r AML), and 6 healthy donors (HD; **Fig. 1A**^29^). Clustering and dimensional reduction using uniform manifold approximation projection (UMAP) identified seven B cell clusters spanning the differentiation continuum from precursor (Pro/Pre) and naïve B cells to plasmablasts and plasma cells. Activated and memory B cells were detected, including a distinct subset of activated B cells with atypical transcriptional profiles herein described as atypical memory B cells (B_atyp_; **Fig. 1B**).

**Figure 1:**
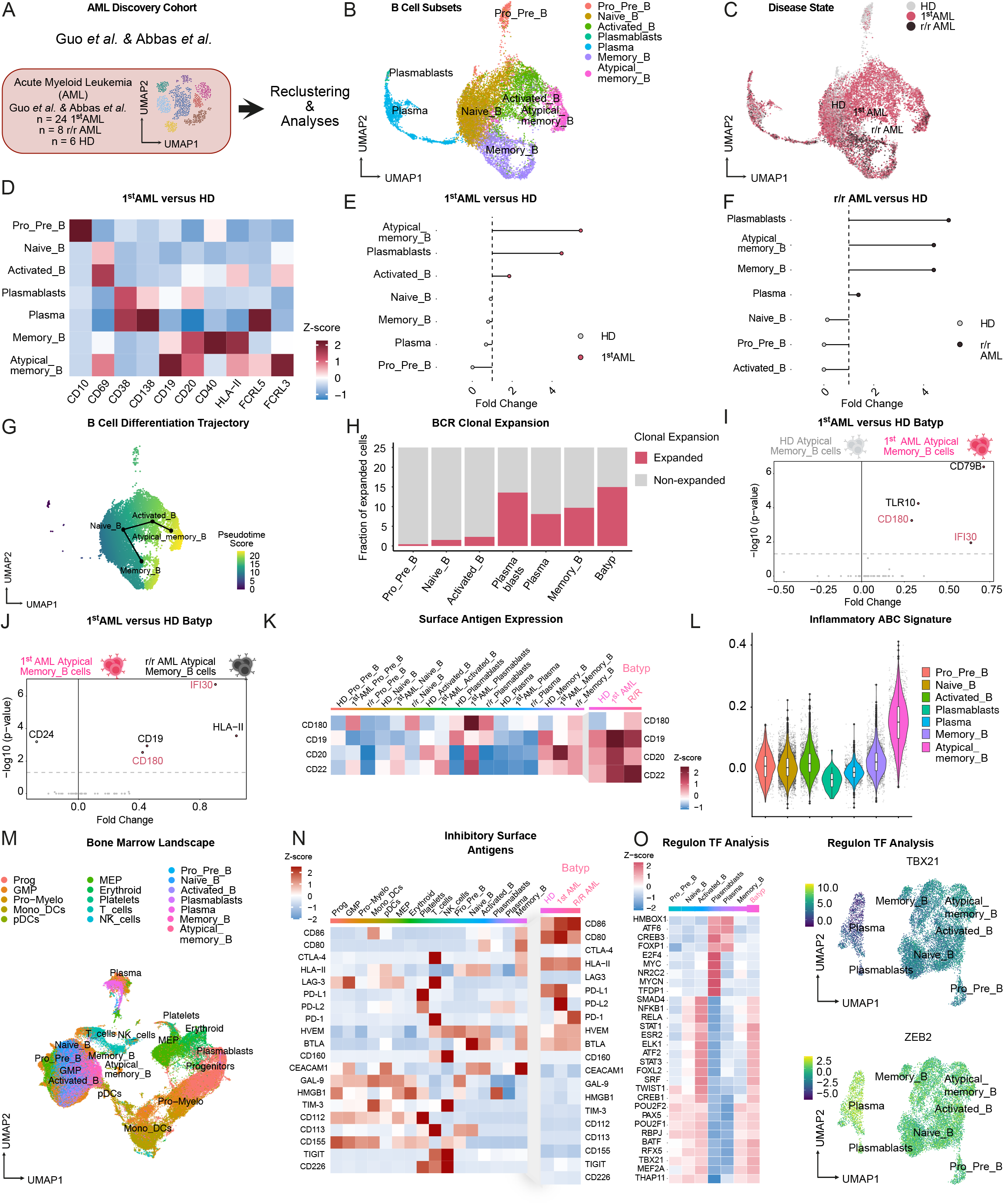
CD180 atypical memory B cells accumulate in AML with an inhibitory phenotype. **(A)** Patient summary and workflow of analyzed dataset of *Guo et al*.^29^ *(n = 12.423 sequenced cells) & Abbas et al..* (**B**) AML cohort of *Guo et al.* ^29^ visualized in an UMAP embedding (bone marrow (BM) samples of n = 24 first diagnosis (1^st^) AML patients, n = 8 relapsed or refractory (r/r) AML patients, n = 6 healthy donors (HD). (**C**) UMAP embedding showing the distribution of B cell subsets across HD, 1^st^ AML and r/r AML. (**D**) Heatmap visualization of the scaled averaged gene expression of the top 10 differentially expressed genes distinguishing the identified B cell subsets. **(E,F)** Lollipop plots showing enrichment of the indicated B cell subsets based on normalized fraction analyses comparing 1^st^ AML versus HD (**E**) and r/r AML versus HD (**F**). (**G**) Pseudotime analysis illustrating B cell differentiation trajectories and corresponding pseudotime scores across identified B cell subsets. (**H**) B cell receptor (BCR) sequencing illustrating clonal expansion of B cell subsets based on shared clonotypes. (**I, J**) Volcano plots showing log fold changes and bonferroni-adjusted *P* values from differential gene-expression analyses (DGE) comparing atypical memory B cells (B_atyp_) from HD and 1^st^ AML (**I**), and from 1^st^ AML and r/r AML (**J**). (**K**) Heatmap showing scaled average gene expression of CD180 compared with hallmark B cell antigens CD19, CD20, and CD22 across identified B cell clusters. (**L)** Violin plot of an inflammatory ABC signature, derived from chronic inflammatory disease-associated ABCs (Holla *et al*.^21^) across AML-associated B cell clusters. **(M**) UMAP embedding of the AML discovery cohort (Abbas, Guo *et al.*^29,72^*, n = 128.094* sequenced single cells). **(N**) Heatmap visualization of the scaled averaged gene expression of hallmark immune inhibitory pathways. Markers were adapted from Chen and Flies^73^. **(O)** DoRothEA^38^ & decoupleR^39^regulon activity analysis of B cell subclusters. Left: heatmap of regulon activity across clusters. Right: UMAP colored by *TBX21* (top) or *ZEB2* (bottom) regulon activity.

B cell subsets differed in relative abundance between AML and HD samples (**Fig. 1C**) and displayed characteristic expression profiles of canonical B cell markers. B_atyp_ were further distinguished by high expression of *CD69*, *FCRL3*, and *FCRL5* (**Fig. 1D**). Cell composition analyses comparing 1^st^ AML versus HD and r/r AML versus HD revealed a selective enrichment of B_atyp_ in AML patients (**Fig. 1E,F; Extended Data Fig. 1A,B**).

To delineate potential development pathways of B_atyp_, we performed Slingshot trajectory analyses and calculated pseudotime scores. Pseudotime mapping suggested a differentiation trajectory from naïve B cells toward either conventional memory or atypical memory B cells, the latter originating from activated B cells (**Fig. 1G**). B cell receptor (BCR) sequencing further demonstrated strong clonal expansion of B_atyp_, plasmablasts and memory B cells in AML patients (**Fig. 1H**).

To identify surface markers specifically upregulated in B_atyp_, we performed DGE analyses comparing B_atyp_ from HD versus AML (**Fig. 1I**) and from 1^st^ AML versus r/r AML (**Fig. 1J**). Among B cell– associated antigens, *CD180* and *IFI30* were consistently upregulated. In contrast to *IFI30*, *CD180* is a transmembrane protein and can therefore serve as a potential surface marker to distinguish B_atyp_^30^. At the single-cell level, *CD180* expression paralleled that of canonical B cell markers (*CD19*, *CD20*, *CD22*) but the expression showed a progressive disease-selective gradient on B_atyp_: minimal in HD and increasing from 1^st^ AML to r/r AML patients (**Fig. 1K**).

As introduced, B_atyp_ are frequently characterized by expression of *ITGAX* (CD11c). In our discovery cohort for example, a ligand–receptor signature from manually curated interactions between atypical memory B cells and leukemic blasts, was associated with adverse outcomes in AML^29^. To interrogate whether a simple four gene signature consisting of the single cell-derived B_atyp_ markers *FCRL3*, *FCRL5*, *ITGAX* and *CD180* would recapitulate this association, we utilized the bulk RNA-sequencing TCGA-LAML cohort (n = 173 AML patients). Similar to the stablished signature, our four gene signature was associated with reduced overall survival in the TCGA-LAML cohort (**Extended Data Fig. 1C**).

We validated these findings in an independent scRNA-seq cohort of 20 patients with AML (1^st^ AML) and 10 HD (**Extended Data Fig. 1D**)^28^. Reference mapping to our AML discovery cohort identified the described seven B cell subsets (**Extended Data Fig. 1E**) with differential distribution between AML and HD (**Extended Data Fig. 1F**). B_atyp_ were again most enriched in AML (**Extended Data Fig. 1G–I**), and trajectory analysis recapitulated the pseudotime progression from activated to B_atyp_ **(Extended Data Fig. 1J**). DGE confirmed the upregulation of *CD180* and *IFI30*, already detected in the first cohort (**Extended Data Fig. 1K**).

B_atyp_ have been most comprehensively described in chronic inflammatory conditions such as SLE and in chronic infectious diseases such as HIV^19^ and malaria^34^. To assess whether AML-associated B_atyp_ share a transcriptional program with these cells, we derived a differential gene expression signature from atypical B cells of patients with malaria, HIV, and autoimmune disease, herein termed atypical B cell (ABC) signature, and scored it across our AML cohorts^21^. Among B cell subsets, the ABC signature was most strongly expressed in B_atyp_ (**Fig. 1L**). The majority of genes upregulated in chronic-inflammatory atypical B cells, were highly expressed in AML-derived B_atyp_ (**Extended Data Figure 1L**).

Together, our analyses across two independent AML cohorts revealed a consistent enrichment of B_atyp_ in the AML bone marrow, whose transcriptional programs resembled those of atypical B cells found in chronic inflammatory and infectious diseases.

### B_atyp_ acquire a T-bet-linked, Breg-like suppressive transcriptional program in AML

As highlighted above, B_atyp_ have been associated with poor patient outcomes in AML, but their mechanism of development and functional characteristics remain poorly defined^28,29^. To investigate potential functions of these cells, we first embedded all cell types from the AML discovery cohort^29^ into a single UMAP (**Fig. 1M)** and quantified expression of hallmark inhibitory and activating molecules. B_atyp_ were positive for *CD86* and *CD80*, consistent with their activated and memory B cell origin (**Fig. 1N**). Interestingly, B_atyp_ had high expression of *PD-L1* and, to a lesser extent, *PD-L2*, as well as strong expression of *HVEM* and *BTLA*. The PD-L1/PD-L2-PD-1 axis and the HVEM-BTLA axis are known to be a potent inhibitor of T cell activity^35^. Expression of activating signaling molecules did not differ substantially from activated or memory B cells (**Extended Data Fig. 2A**).

Given the high expression of T cell-inhibitory ligands by B_atyp_, we applied LIANA (LIgand-receptor ANalysis frAmework) to infer their interactions within the bone marrow microenvironment^36^. At the transcriptomic level, this predicted potential ligand-receptor interactions between B_atyp_, T cells, and AML blasts residing within progenitor compartment. The predicted interactions differed between patients with AML and HD (**Extended Data Fig. 2B**).

In chronic inflammatory and infectious conditions, atypical B cells are closely associated with the transcription factors T-bet (*TBX21*)^34^ and ZEB2^37^. To identify the transcriptional drivers of B_atyp_ in AML, we performed regulon analysis (DoRothEA^38^, decoupleR^39^). The regulon program of B_atyps_ resembled that of activated B cells and memory B cells (**Fig. 1O**), consistent with the inferred differentiation trajectory from activated B cells. At the level of individual transcription factors, *TBX21* (T-bet) was expressed by B_atyp_, with a trend toward higher expression than in other B cell subsets, whereas *ZEB2* showed neither high nor B_atyp_-selective expression (**Fig. 1O, Extended Data Fig. 2C**). Thus, AML-associated B_atyp_ partially recapitulate the T-bet-linked program of canonical atypical B cells, while potentially lacking their prominent ZEB2 association^37^. Next, we performed chemokine-receptor repertoire profiling of AML-associated B_atyp_. B_atyp_ retained CXCR3 and CXCR5, the latter at lower levels than naïve or activated B cells, together with low CXCR4 and comparatively high CCR6 and CCR7 (**Extended Data Fig. 2D**). Overall, at the transcriptomic level, this homing profile can be described as tissue- and inflammation-associated, with retained but reduced CXCR5 expression, consistent with the reported heterogeneity of atypical B cells spanning CXCR5 and CXCR5-low states.

To further classify the B_atyp_ cell population, we characterized the expression of B regulatory cell (Bregs)-associated surface (**Extended Data Fig. 2E, top**) or functional markers (**Extended Data Fig. 2E, bottom)**. AML-associated B_atyp_ showed low expression of *CD24*, *CD38*, *CD1d* and *CD5* and therefore did not conform to the classical transitional (CD24^high^CD38^high^) or B10 (CD5 CD1d^high^) Breg phenotypes^14,24,26^. Yet, B_atyp_ expressed functional Breg markers such as *IL-10*, *TGF-*β, *EBI3* and *PD-L1/PD-L2*, suggesting they acquire a Breg-like immunosuppressive gene program.

Historically, atypical B cells were defined within the CD27 IgD double-negative B cell fraction, and our scRNA-seq data recapitulated their low surface-marker expression of CD27 and IgD (**Extended Data Fig. 2F**). More recent work, however, has revealed a more nuanced, isotype-heterogeneous picture, with atypical B cells in chronic infection and autoimmunity spanning both an unswitched IgD IgM^low^ compartment^21^ and a class-switched, IgG-dominated compartment^40^. Consistent with this heterogeneity, immunoglobulin heavy-chain (IGHC) isotype analysis showed that AML-associated B_atyp_ comprised a dominant unswitched IgM fraction together with class-switched cells (predominantly IgG, with a smaller IgA fraction) and only a minor IgD population (**Extended Data Fig. 2G**).

Together, these data define AML-associated B_atyp_ as a T-bet-linked atypical B cell state that acquires a Breg-like, immunosuppressive transcriptional program.

### AML-derived soluble factors polarize B cells into immunosuppressive B_atyp_

AML is a molecularly heterogeneous disease whose outcome strongly depends on the underlying genetic subtype^41^. To assess whether B_atyp_ transcriptional programs are enriched in specific molecular subtypes, we pooled three bulk RNA-sequencing cohorts (phs001657.v3.p1, BEAT-AML 2.0; GSE6891^42^; GSE146173^43^; n = 1,490 patients with AML; **Extended Data Fig. 3A**) and scored the B_atyp_ signature, with additional weighing of the B-cell compartment by including the pan-B cell markers CD19 and CD20. The B_atyp_ score was highest in core-binding-factor (CBF) AML and RAS-pathway-mutated AML, with a trend toward higher scores in NPM1-mutated AML and in AML with adverse-risk cytogenetics, whereas TP53-mutated and FLT3-ITD AML showed lower scores (**Extended Data Fig. 3B,C**).

To validate these findings at the protein level, we performed multicolor flow cytometry in a discovery cohort including 29 1^st^ AML, 8 r/r AML, and 8 HD bone marrow (BM) samples (**Fig. 2A, see Extended Data Fig. 3D** for gating strategy). We observed a marked enrichment of CD27 IgD cells in 1^st^ AML (**Fig. 2B**), while r/r AML samples had lower frequencies of these cells but higher proportions of naïve B cells (must likely due chemotherapy treatment). Within the CD27 IgD subset, PD-L1 and BTLA expression was elevated in both 1^st^ AML and r/r AML compared with healthy donors, while we did not observe differences in CD86 expression. CD180 was consistently expressed across these B cell fractions (**Fig. 2C**).

**Figure 2:**
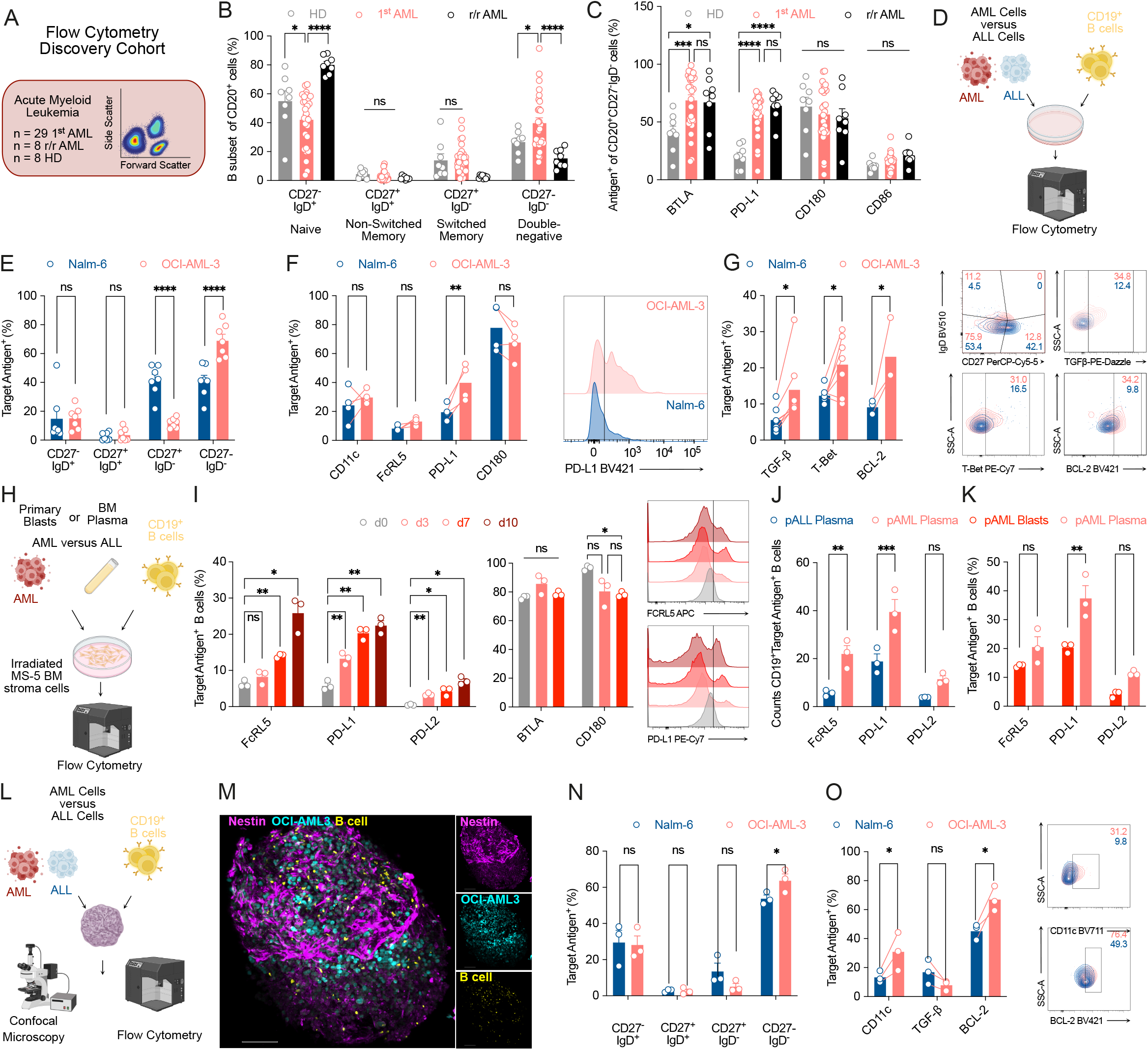
AML-derived soluble factors polarize B cells into immunosuppressive B_atyp_. **(A)** Overview of the flow cytometry discovery cohort. **(B)** Frequencies of B cell subsets in HD, 1^st^ AML, and r/r AML samples determined by flow cytometry. **(C**) Expression frequencies of inhibitory signaling molecules on B_atyp_ (CD20 CD27 IgD ). **(B, C)** Data are mean ± s.e.m.; n = 29 1st AML, n = 8 r/r AML, n = 8 HD. **(D)** Experimental workflow of AML–B cell co-cultures. **(E)** Frequencies of B cell subsets after co-culture with OCI-AML3 (AML) or Nalm-6 (ALL) cell lines, by flow cytometry. **(F, G)** Left: expression of the indicated markers on CD19 B cells. Right: representative flow cytometry plots. **(E-G)** Data are mean ± s.e.m. from n = 4–6 independent donors. **(H)** Experimental workflow of primary AML-B cell co-cultures. **(I)** Left, Middle: Expression of the indicated markers on B cells at the indicated timepoints by flow cytometry. Right: representative flow cytometry plots. **(J)** Counts of target antigen positive B cells after 7 days of polarization in 50% BM pAML or pALL plasma. **(K)** Expression of the indicated markers on B cells co-cultured for 7 days with pAML blasts or BM plasma. **(I–K)** Data are mean ± s.e.m. from n = 3 independent B cell donors. **(L)** Experimental workflow of AML-B cell-BMO triple co-cultures. B cell to tumor cell ratio: 1:1. **(M)** Representative merged confocal z-stack images of AML-B cell–BMO co-cultures. Single-channel images (right) display BMO mesenchymal stromal cells (Nestin, violet), OCI-AML3 tumor cells (turquoise) and B cells (yellow). Scale bar: 100 µm. **(N,O)** Expression of indicated markers on B cells from engrafted BMO measured by flow cytometry. Data are mean ± s.e.m from n = 3 independent donors. Unless otherwise indicated, statistical analysis was performed using two-way ANOVA with Tukey’s multiple comparison test. For all panels: \**P* < 0.05; \*\**P* < 0.01; \*\*\**P* < 0.001; \*\*\*\**P* < 0.0001; ns, not significant.

We next stratified patients by European LeukemiaNet (ELN) 2022 risk classification^41^ or by molecular subtype. B cell subset frequencies did not differ across ELN risk groups (**Extended Data Fig. 3E, left**). CD180 was expressed at slightly lower levels on CD20 CD27 IgD B cells from patients with adverse-risk cytogenetics, whereas BTLA and PD-L1 expression did not differ (**Extended Data Fig. 3E, right**). Likewise, we observed no differences in B cell subset composition or in BTLA, PD-L1, and CD180 expression on CD20 CD27 IgD B cells across NPM1-mutated, CBF, FLT3-ITD, or myelodysplasia-related (AML-MR) AML (**Extended Data Fig. 3F**). Flow cytometric results were confirmed in an independent validation cohort (**Extended Data Fig. 3G**), with increased frequencies of CD27 IgD cells in both peripheral blood and bone marrow of patients with AML (**Extended Data Fig. 3H**). Again, PD-L1 expression was highest on AML-derived CD27 IgD (switched memory) and CD27 IgD (B_atyp_) B cell subsets (**Extended Data Fig. 3I**).

To test whether AML cells actively support B_atyp_ expansion or persistence, we co-cultured MACS-isolated CD19 B cells from healthy donors with AML cell lines or, as a control, with the Nalm-6 acute lymphoblastic leukemia (ALL) cell line (**Fig. 2D**). Co-culture with OCI-AML3 or MOLM-13, and to a lesser extent PL-21, increased donor B cell counts, indicating that AML cells promote B cell proliferation or persistence (**Extended Data Fig. 4A**). We did not observe an increased B cell count in co-cultures with Nalm-6. Phenotypically, co-culture with OCI-AML3 further increased the frequency of CD27 IgD B cells (**Fig. 2E**) and, compared with Nalm-6, induced upregulation of PD-L1, TGF-β, T-Bet, and BCL-2 (**Fig. 2F,G**).

To confirm these findings in a more patient-relevant setting, we established a primary co-culture system in which primary AML blasts (pAML), or bone-marrow plasma (BM plasma) from AML or ALL patients, were co-incubated with healthy-donor CD19 B cells on irradiated MS-5 feeder cells (**Fig. 2H**). B cells cultured on the MS-5 feeder layer alone, without AML, did not survive, indicating that B-cell persistence and phenotype in this system depend on the AML compartment rather than the feeder layer (**Extended Data Fig. 4B)**. Longitudinal profiling of B cells co-cultured with pAML blasts revealed a gradual increase in FCRL5, PD-L1, and PD-L2, whereas BTLA and CD180 were highly expressed throughout, with a minor decrease in CD180 by day 7 (**Fig. 2I**). CD19 expression likewise decreased over time and there were no difference in expression between CD19 and CD180 (**Extended Data Fig. 4C**).

To test whether direct cell–cell contact is required to induce the B_atyp_ phenotype, we cultured CD19 B cells on MS-5 feeders with 50% BM plasma from AML or ALL patients. B-cell counts were markedly higher with pAML plasma than with pALL plasma (**Extended Data Fig. 4D**), and the numbers of FCRL5 , PD-L1 , and PD-L2 B cells were correspondingly higher (**Fig. 2J**). Directly comparing B cells exposed to pAML blasts versus pAML plasma, FCRL5 and PD-L2 did not differ, whereas PD-L1 was higher with plasma (**Fig. 2K**), indicating that soluble factors are sufficient to induce the B_atyp_ phenotype.

To further corroborate these findings in a human model system, we engrafted human iPSC-derived bone-marrow organoids (BMO)^44^ with the AML cell line OCI-AML3 together with HD CD19 B cells at different ratios (**Fig. 2L**). Engraftment of both AML and B cells within the BMO was confirmed by confocal microscopy (**Fig. 2M, Extended Data Fig. 4E**). After seven days, flow cytometry revealed enrichment of CD27 IgD B_atyp_, with elevated expression of CD11c and BCL-2 as well as PD-L2 (**Fig. 2N,O; Extended Data Fig. 4F**), supporting the notion that B_atyp_ can be polarized by AML blasts in the BM environment.

Overall, using various different models of AML cell lines and primary AML blasts, we demonstrate that AML blasts-derived soluble mediators induce the upregulation of an proinflammatory, immunosuppressive and anti-apoptotic B_atyp_-gene program with upregulation of T-bet and inhibitory cell surface markers (FCRL5, PD-L1, PD-L2), cytokines (TGF-β) and anti-apoptotic BCL-2.

### B_atyp_ are highly enriched in patients with AML and MM not responding to CAR T cell therapy

Clinical responses to chimeric antigen receptor (CAR) T cell therapy in AML have been limited, with rates of at best 25 % in recent early studies^10^.

To investigate whether B_atyp_ contribute to this lack of efficacy, we analyzed a scRNA-seq dataset of 11 AML patients treated with CD123 CAR T cells within a clinical trial at the University of Pennsylvania (**Fig. 3A**, NCT03766126^10^). The dataset was successfully reference-mapped to our AML discovery cohort^29^ (**Fig. 3B**) and clustered according to clinical response (complete response, CR; non-complete response, non-CR; **Fig. 3C**). B_atyp_ were strikingly enriched in patients, who did not respond to CD123 CAR T cell therapy (**Fig. 3D-F**). Phenotypically, these B_atyp_ expressed high levels of *CD86*, *CD80*, *PD-L2*, and to a lesser extent *PD-L1*, as well as *HVEM* and *BTLA*, overall recapitulating the above described suppressive immunophenotype (**Extended Data Fig. 5A**). Expression of activating signaling molecules was again similar across different B cell subsets, with a few notable expression differences (**Extended Data Fig. 5B**). Turning to the T cell compartment, unsupervised clustering identified twelve T cell clusters, annotated by differential gene expression, comprising distinct CD4 and CD8 effector states, including a large cluster of exhausted T cells (T_Exh) as well as Tregs and NK-like T cells (NK_T; **Fig. 3G,H**). In non-CR patients, T_Exh was the most enriched cluster, followed by NK_T and cycling T cells (**Fig. 3I**). To examine potential interactions between T cells and B_atyp_, we again performed LIANA analysis. Consistent with previous analyses, the T-B cell interactions were dominated by BTLA and *TGF-*β (**Fig. 3J**).

**Figure 3:**
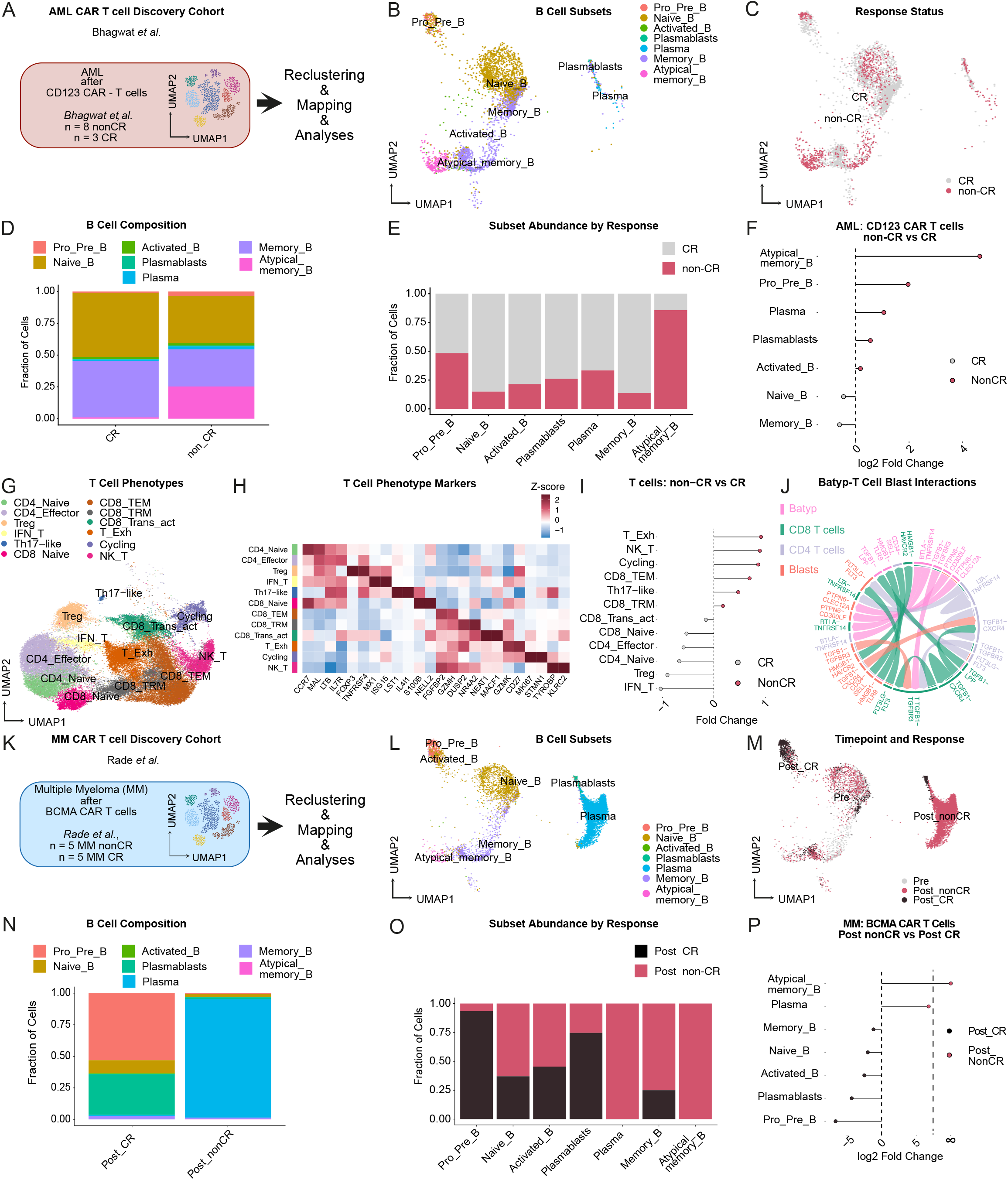
B_atyp_ are enriched in AML and MM patients not responding to CAR T cell therapy. (**A**) Overview of the analyzed scRNA-seq dataset from Bhagwat *et al.*^10^, comprising AML patients treated with CD123 CAR T cells within the clinical trial NCT03766126. (**B**) Reference mapping of B cell subsets from Bhagwat *et al.* to the AML discovery cohort. (**C**) UMAP visualization of AML samples clustered according to treatment response (complete response, CR; non-complete response, non-CR). (**D**) Bar plots showing the relative frequencies of individual B cell subsets in CR versus non-CR cohorts. (**E**) Distribution of B cell subsets in CR and non-CR patients. (**F**) Lollipop plot showing enrichment of B cell subsets in patients achieving CR versus non-CR based on normalized fraction analysis. (**G**) UMAP embedding showing the distribution of T cell subsets. (**H**) Heatmap of the scaled average gene expression of the top two differentially expressed genes distinguishing the identified T cell clusters. (**I**) Lollipop plot showing enrichment of T cell subsets in CR versus non-CR patients, based on normalized fraction analysis. (**J**) Chord diagrams of predicted ligand–receptor interactions between B_atyp_, T cells, and AML blasts in non-CR patients. (**K**) Overview of the analyzed scRNA-seq dataset from Rade *et al.*^45^, comprising multiple myeloma (MM) patients treated with BCMA CAR T cells (*n* = 10 MM patients). (**L**) Reference mapping of B cell subsets from Rade *et al.* to the discovery cohort. (**M**) UMAP visualization of MM samples obtained before (Pre) and after (Post) therapy, clustered according to treatment response (CR versus non-CR). (**N**) Bar plots showing the relative frequencies of individual B cell subsets in Post_CR and Post_NonCR cohorts. (**O**) Distribution of B cell subsets in Post_CR and Post_NonCR patients. (**P**) Lollipop plot showing enrichment of B cell subsets in patients achieving CR versus non-CR based on normalized fraction analyses.

To generalize our findings in other malignancies and CAR T products, we analyzed a second dataset of 10 multiple myeloma (MM) patients treated with B cell maturation antigen (BCMA)-targeted CAR T cells^45^. Reference mapping of bone marrow mononuclear cells (BMMCs) identified similar B cell subsets with expected enrichment of plasma cells, linked to the underlying disease (**Fig. 3K**). Grouping based on treatment response (Post_CR, Post_NonCR) and pre-treatment samples (Pre) revealed that while plasma cells dominated in non-responders, B_atyp_ were exclusively detected in patients failing to respond to BCMA CAR T cell therapy (**Fig. 3L-P**). To validate these findings, we analyzed an independent dataset comprising matched pre- and post-treatment BMMCs from 7 MM patients treated with BCMA CAR T cells^46^ (**Extended Data Fig. 5C**). Despite lower cell numbers, reference mapping identified seven major B cell subsets (**Extended Data Fig. 5D,E**). B_atyp_ frequencies were markedly higher before therapy and decreased in responding patients, linking their abundance to active disease (**Extended Data Fig. 5F-H**). In MM, B_atyp_ exhibited similar expression profiles as in AML, with elevated *CD86*, *PD-L1*, *PD-L2*, *HVEM*, and *BTLA* (**Extended Data Fig. 5I,J**).

Together, these data show that immunosuppressive B_atyp_ are enriched in patients who fail to respond to CAR T cell therapy across hematologic malignancies, and display a conserved inhibitory phenotype in AML and MM.

### B_atyp_ suppress conventional and CAR T cell responses in AML

ScRNA-seq and flow cytometry revealed a consistent enrichment of B_atyp_ in patients with AML and in non-responders to CD123- or BCMA-directed CAR T cell therapy. We also showed that AML-derived soluble mediators are sufficient to polarize B cells into B_atyp_. These cells, in turn, consistently upregulated inhibitory pathways predicted to mediate interactions with T cells. Yet how B cells in general, and B_atyp_ in particular, are spatially organized within the AML bone marrow niche, whether they are positioned to engage T cells, and whether they carry immunosuppressive functions remain uncharacterized.

We therefore performed multiplexed immunofluorescence (mIF) on five primary AML samples, staining for CD180, CD20, CD3, PD-L1, and CD34 to map the spatial organization of AML blasts (CD34 ), B cells (CD20 ), and T cells (CD3 ) (**Fig. 4A**). CD180 was broadly expressed across the bone marrow microenvironment, on both B cells and CD34 blasts (**Fig. 4B, Extended Data Fig. 6A**). CD20 B cells and CD3 T cells were detected in close spatial proximity throughout the marrow in all five patients (median distance to the nearest T cell 17.1–22.0 µm), confirming that B-T cell encounters occur within the AML bone marrow niche (**Fig. 4C, Extended Data Fig. 6B**).

**Figure 4:**
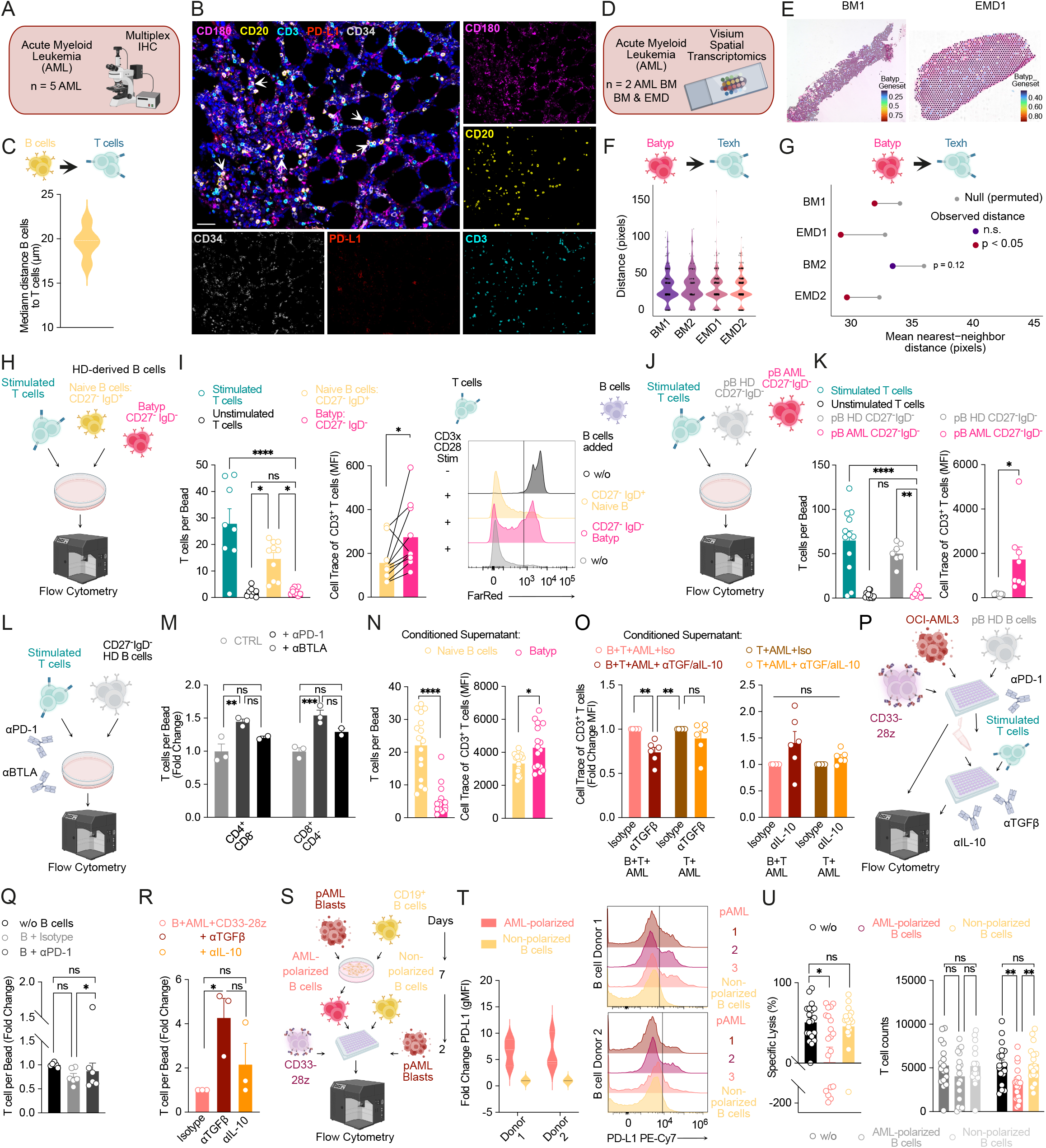
B_atyp_ suppress conventional and CAR T cell responses in AML. **(A)** Overview of the multiplex immunofluorescence (mIF) cohort. (**B**) Representative merged mIF of primary AML bone marrow biopsies. Single channel images (right) show CD180 (violet), CD20 (yellow, B cells), CD3 (turquoise, T cells), PD-L1 (red), and CD34 (grey, AML blasts). Scale bar, 100 µm. (**C**) Violin plot depicting median distance from B to T cells; n = 5 primary AML samples. (**D**) Overview of analyzed spatial transcriptomic cohort^47^. (**E**) Spatial feature plot depicting B_atyp_ signature score expression per Visium spot in primary bone marrow (BM) or extramedullary disease (EMD). **(F**) Violin plot depicting median distance from B_atyp_ to T_exh_ cells. (**G**) Mean nearest-neighbor distance (in pixels) between the B_atyp_ and T_exh_ geneset in the Visium spatial transcriptomic data. (**H**) Experimental workflow of T cell proliferation assays with isolated B cell subsets. B cells were isolated from healthy donors (HD) by magnetic-activated cell sorting (MACS). T cells were labeled with a far red proliferation dye and, unless otherwise indicated, stimulated with CD3/CD28 magnetic beads. **(I**) Flow cytometric quantification of T cell proliferation after co-culture with naïve (CD27 IgD , yellow) or atypical memory (B_atyp_; CD27 IgD , violet) B cells. Stimulated or unstimulated T cells cultured without B cells served as controls. Left: T cell counts per bead. Middle: gMFI of the far red proliferation dye. Statistical analysis by unpaired t-test. Right: representative flow cytometry plot. Data are mean ± s.e.m. from n = 10 independent T cell donors. (**J**) Experimental workflow: B cells isolated from the blood of either HD or patients with AML. (**K**) Flow cytometric quantification of T cell proliferation after co-culture with HD-derived (grey) or AML-derived (violet) B cells. Left: T cell counts per bead. Statistical analysis by unpaired t-test. Right: gMFI of the far red proliferation dye. Data are mean ± s.e.m. from n = 7-8 independent T cell donors. (**L**) Experimental workflow of T cell–B cell co-cultures with αPD-1 and αBTLA checkpoint blockade. (**M**) Flow cytometric quantification of T cell counts (fold change relative to control, CTRL). Data are mean ± s.e.m. from n = 2–3 independent T cell donors. (**N**) Flow cytometric quantification of T cell counts (left) or proliferation dye trace dilution (right) of stimulated T cells co-incubated with conditioned supernatant from naïve B cells (yellow) or B_atyp_ (pink). Data are mean ± s.e.m. of n = 16 T cell-supernatant stimulations. Statistical analysis by unpaired t-test. (**O**) Flow cytometric quantification of T cell proliferation (fold change relative to isotype) of stimulated T cells co-incubated with conditioned supernatants from B+T+AML-triple-co-cultures or T+AML control co-cultures with TGFβ-(αTGFβ, left) or IL-10-neutralizing antibody (αIL-10, right). Data are mean ± s.e.m. of n = 5-6 independent T cell donors. (**P**) Experimental workflow: OCI-AML3 cells and CD33-28z CAR T cells co-cultured with or without HD-derived B cells, with αPD-1 or isotype control. Collected supernatants then co-incubated with stimulated T cells with αTGFβ or αIL-10 or isotype control. (**Q**) Flow cytometric quantification of T cell counts (fold change relative to the non–B cell condition). Data are mean ± s.e.m. from n = 6 independent donors. (**R**) Flow cytometric quantification of T cell counts (fold change relative to isotype control) co-cultured supernatant of CAR-AML-B cell co-cultures with αTGFβ or αIL-10 or isotype control. (**S**) Experimental workflow: B cells were polarized with pAML blasts for 7 days, then co-cultured with CD33-28z CAR or untransduced T cells (UT) for 48 h at an E:T ratio of 1:10. (**T**) PD-L1 expression on polarized versus non-polarized B cells. (**U**) Flow-cytometric quantification of CD33-28z–mediated lysis of pAML blasts (left) and T cell counts (right). Data are mean ± s.e.m. from n = 19 B cell and T cell donors. Unless otherwise indicated, statistical analysis used two-way ANOVA with Tukey’s multiple comparison test. For all panels: \**P* < 0.05; \*\**P* < 0.01; \*\*\**P* < 0.001; \*\*\*\**P* < 0.0001; ns, not significant.

To extend these findings to B_atyp_, we analyzed a published Visium spatial transcriptomic dataset of bone-marrow (BM) and extramedullary disease (EMD) biopsies from two AML patients^47^ (**Fig. 4D**). We scored a B_atyp_ signature (*FCRL3, FCRL5, CD180, CD19, CD20, ITGAX*) and an exhausted T cell signature (*IL10, HAVCR2, GRB2, PTPN2, CD3E*), which was upregulated on T cells with non-CR treated with CD123 CAR T cells, across the Visium spots (**Fig. 4E, Extended Data Fig. 6C**). The median B_atyp_-T_exh_ cell distance was comparable between BM and EMD (**Fig. 4F**). Nearest-neighbor analysis showed significant B_atyp_-T_exh_ cell co-localization in the two EMD sample and one bone marrow sample, with a similar trend in the remaining samples (**Fig. 4G**).

Next, to investigate whether B_atyp_ inhibit T cells, we isolated naïve (CD27^−^IgD ) and atypical memory (CD27 IgD ) B cell subsets from healthy donor and co-cultured them with autologous, activated T cells after stimulation with CD3-CD28 activation beads (**Fig. 4H**). T cell proliferation drastically reduced in the presence of B_atyp_ compared to naïve B cells, as seen by a reduction in overall T cell counts and an increased geometric mean fluorescence intensity (gMFI) of the proliferation dye, indicating lower T cell proliferation (**Fig. 4I**). Stimulated or unstimulated T cells (not co-cultured with B cells) were used as controls. T cells co-cultured with naïve B cells expressed higher levels of PD-1, whereas CD25 and CD69 were comparable across conditions, consistent with their higher proliferation (**Extended Data Fig. 6D**). We next asked whether AML-derived B_atyp_ behave similarly. CD27 IgD B cells isolated from peripheral blood (PB) or bone marrow (BM) of patients with AML strongly reduced proliferation of co-cultured activated T cells, without differences between PB-and BM-derived cells (**Extended Data Fig. 6E,F**). Comparing HD- and AML-derived B_atyp_ head-to-head, AML-derived B_atyp_ suppressed T cell proliferation more potently (**Fig. 4J,K**), without changes in T cell activation markers (**Extended Data Fig. 6G**).

Because B_atyp_ expressed high levels of PD-L1, PD-L2, and BTLA, we asked whether these axes contribute to the suppression of T cell proliferation. To this end, we tested whether the inhibition could be reversed with the anti-PD-1 antibody nivolumab (αPD-1) or the anti-BTLA antibody tifcemalimab (αBTLA; **Fig. 4L**). Addition of αPD-1 increased T cell proliferation ∼1.5-fold, and αBTLA showed a similar trend (**Fig. 4M**). We did not observe differences in the expression of activation markers (CD69, 4-1BB, CD25) on T cells (**Extended Data Fig. 6H**). Suppression by B_atyp_ did not exclusively require direct cell contact: B_atyp_-conditioned supernatant reduced T cell proliferation (**Fig. 4N**). Supernatant from B cell–T cell–AML co-cultures suppressed proliferation in the same way, and this was partially reversed by TGF-β, but not IL-10, neutralization (**Fig. 4O**). This was dependent on B cell-derived TGF-β, as we did not see a reversal of suppression in T-AML co-cultures.

Given that AML cells polarize B cells toward B_atyp_ and that B_atyp_ were enriched in patients not responding to CAR T cell therapy, we tested whether AML-associated B cells inhibit CAR T cell expansion. We co-cultured OCI-AML3 cells with second-generation CD28-costimulated anti-CD33 CAR T cells (CD33-28z) in the presence or absence of HD-derived B cells, with αPD-1 or isotype control (**Fig. 4P**). CD33-28z CAR T cell numbers decreased in the presence of B cells and isotype control, but not when αPD-1 was added (**Fig. 4Q**), indicating that polarized B cells suppress CAR T cells through the PD-1–PD-L1 axis. Transferring supernatant from these co-cultures onto stimulated T cells reproduced the soluble arm: only αTGF-β but, not αIL-10, increased T cell proliferation (**Fig. 4R**).

To confirm this mechanism in a more patient-relevant setting, we polarized HD-derived B cells with primary AML blasts from three patients for 7 days and then co-cultured the polarized B cells with CD33-28z CAR T cells. Freshly isolated, non-polarized B cells and CAR T cells alone served as controls (**Fig. 4S**). As before, co-culture with pAML blasts upregulated PD-L1, confirming successful polarization (**Fig. 4T**). Strikingly, CD33-28z CAR T cells co-cultured with AML-polarized B cells showed reduced lysis capacity, albeit with considerable heterogeneity across the different AML, B cell, and T cell donors (**Fig. 4U**). Notably, CD33-28z CAR T cell counts were reduced only in the presence of polarized, but not unpolarized, B cells, whereas untransduced T cells (UT; negative control) were not impacted (**Fig. 4U).** We did not observe differences in PD-1 expression on the T cells **(Extended Data Fig. 4I).**

Together, B_atyp_ suppress conventional T cells and CD33-directed CAR T cells through the PD-1–PD-L1 axis and soluble TGF-β, reducing proliferation, expansion and cytotoxicity.

### Interferon, TLR and CD70–CD27 signals link AML and CAR T cells to B_atyp_ polarization

While our results indicate that B_atyp_ are polarized within the AML bone marrow microenvironment through soluble mediators, the underlying molecular mechanism remains unclear. We therefore returned to the AML scRNA-seq discovery cohort and performed pathway enrichment analysis between AML-derived B_atyp_ in comparison to other B cell subsets, as well as between AML blasts and HD myeloid cells (progenitors, promyelocytes, MonoDC). AML-derived B_atyp_ upregulated several cytokine-signaling pathways (TNF-α signaling via NF-κB, IFN-γ response, IFN-α response, IL-2/STAT5 signaling, IL-6/JAK/STAT3 signaling) and apoptosis-and p53-related programs (**Extended Data Fig. 7A,B**), broadly recapitulating the polarization experiments. Malignant myeloid cells showed a comparable upregulation of TNF-α, IFN-α, and IFN-γ response pathways (**Extended Data Fig. 7C–E**), consistent with malignant AML cells providing critical signals.

The prevailing model holds that the atypical, T-bet CD11c B cell (ABC) program is instructed by at least two inputs: an innate signal one through TLR7 (or TLR9) that licenses the B cell, followed by a signal two from the surrounding cytokine milieu. Within TLR-activated B cells, both IFN-γ and IL-21 can directly induce T-bet, whereas CD11c is driven preferentially by IL-21 and largely independently of T-bet. IL-4 acts as a contextual modulator, opposing IL-21, but enhancing IFN-γ– driven T-bet induction^48^.

To explore the responsible inputs more broadly, we leveraged a recently published perturbation atlas of human immune responses, in which PBMCs were stimulated *in vitro* with 90 individual cytokines^49^ (**Fig. 5A**). Using this resource, we asked which cytokine inputs activate defined modules of the ABC program, focusing on the B cell compartment. We grouped *NFKB1*, *BCL2* and *IL10* as an NF-κB-dependent module, and *TBX21 (*T-Bet) together with *PDL1* as an IFN/STAT1-dependent module, noting that full T-bet induction additionally requires innate input. Because CD11c can be induced by IL-21 independently of T-bet, we retained it as a separate module. Additional ABC-associated genes (*FCRL3*, *TLR9*, *TLR10*) were analyzed individually.

**Figure 5:**
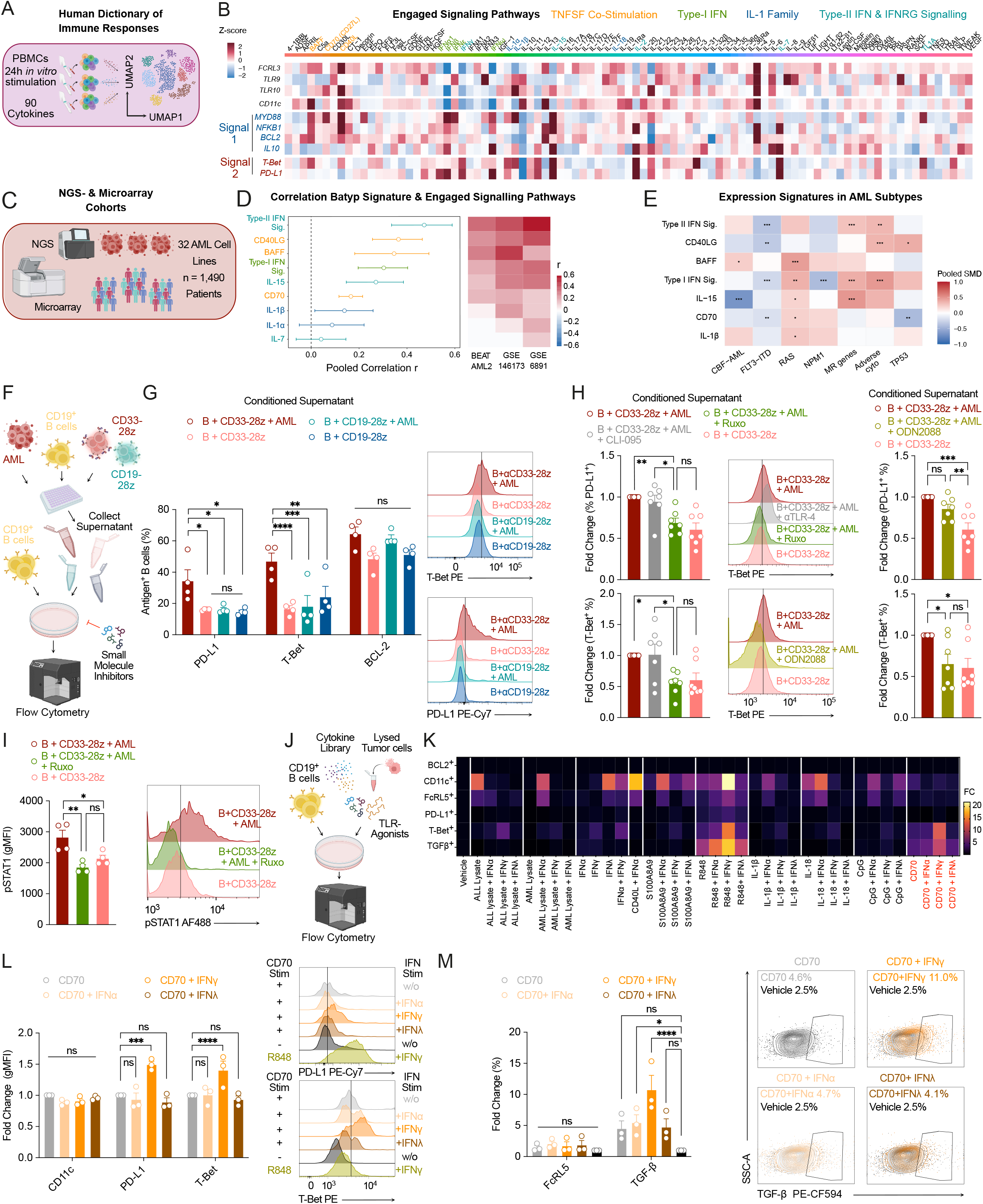
Interferon, TLR and CD70–CD27 signals link AML and CAR T cells to B_atyp_ polarization. **(A)** Summary of the human cytokine dictionary perturbation dataset^74^. (**B**) Heatmap of ABC– associated downstream signaling modules induced in HD PBMCs by each stimulus. (**C**) Overview of the bulk RNA-sequencing datasets ^52^ and microarray AML cohorts. (**D**) Forest plot of the spearman correlation coefficient (r) between the B_atyp_ signature and the indicated cytokines or signatures across AML cohorts (phs001657.v3.p1, BEAT-AML 2.0; GSE6891^42^; GSE146173^43^). (**E**) Expression signatures across molecularly defined AML subtypes. (**F**) Experimental workflow of B cell cultures stimulated with supernatants from the indicated CAR T cell–AML supernatant co-cultures. (**G**) Left: Expression of the indicated markers on B cells stimulated with the indicated supernatants, by flow cytometry. Data are mean ± s.e.m. of n = 3–4 B cell–AML–CAR co-cultures. Right: representative flow cytometry. **(H)** Left: Expression of PD-L1 (top) or T-Bet (bottom) on B cells stimulated with the indicated supernatants and treated with ruxolitinib (Ruxo) or TLR4 antagonist CLI-095, by flow cytometry. Middle: Representative flow cytometry plot of T-Bet expression on B cells. Right: B cells treated with TLR7-9 antagonist ODN2088. Data are mean ± s.e.m. of n = 7-8 B cell–AML–CAR supernatant co-cultures. (**I**) pSTAT1 phosphoflow analysis of B cells stimulated with indicated supernatants with or without Ruxo treatment. Data are mean ± s.e.m. of n = 4 B cell–AML–CAR co-cultures. (**J**) Experimental workflow of B cell cytokine and agonist screening. (**K**) Heatmap illustrating expression of indicated markers as fold change normalized to vehicle control. Data are mean of n = 3 different B cell donors. (**L,M**) Left: Expression of indicated markers by flow cytometry. Right: Representative flow cytometry plot. Data are mean of n = 3 different B cell donors. Unless otherwise indicated, statistical analysis used two-way ANOVA with Tukey’s multiple comparison test. For all panels: \**P* < 0.05; \*\**P* < 0.01; \*\*\**P* < 0.001; \*\*\*\**P* < 0.0001; ns, not significant.

This analysis revealed a complex picture, with distinct ligands inducing distinct components of the program, yet several inputs stood out (**Fig. 5B**). TNFSF ligands linked to B cell homeostasis such as BAFF, CD70 (CD27L), and CD40L strongly induced the signal-1 module (*MYD88*, *NFKB1*, *BCL2, IL-10*). Type-I interferons (type-I-IFNs) activated both modules. The signal-2 module was further induced by lymphoid cytokines IL-2, IL-7, IL-15, and IFN-γ. Because B cells express IL-7Rα only at low levels, little to no IL-15 receptor, and often lack the high-affinity IL-2 receptor complex, the most likely explanation in this mixed PBMC system is that these cytokines act on T and NK cells to elicit IFN-γ, which in turn drives the ABC program in B cells. To confirm this hypothesis, we examined cytokine receptor expression across B cell subsets in our AML discovery cohort.

B_atyp_ had high expression of *TLR7–9*, of both IFN-γ receptor chains (*IFNGR1*, *IFNGR2*) and of complete receptor pairs for IL-21 (*IL21R, IL2RG*), IL-27 (*IL27RA, IL6ST*) and TNF (*TNFRSF1A, TNFRSF1B*) (**Extended Data Fig. 7F**). Interestingly, B_atyp_ also expressed both chains of the type III interferon receptor (*IFNLR1, IL10RB*), possibly indicating a role for these cytokines in ABC biology.

To identify the relevant factors in AML, we then analyzed a bulk RNA-seq dataset of human cell lines^50–52^ and asked which of these pathways are more highly expressed in AML compared to Nalm-6 cells, which were used as a control during our polarization experiments. AML cell lines expressed higher levels of the TNFSF ligands *BAFF*, *CD70*, and *CD40LG*, as well as higher levels of IL-1-family proteins, *type-I IFNs, IL-7*, and *IL-15* (**Extended Data Fig. 7G**). We then correlated these signaling families with the B_atyp_ signature used earlier to describe B_atyp_ abundance in AML patients (**Extended Data Fig. 3A-C**). Across our pooled cohort of n = 1,490 AML patients, the B_atyp_ signature correlated most strongly with the *type-II IFN* signature, and also with *CD40LG*, *BAFF*, the *type-I IFN signature*, *IL-15* and *CD70*. *IL-1*α, *IL-1*β, and *IL-7* correlated more weakly (**Fig. 5D**). The pattern varied with molecular subtype (**Fig. 5E**).

Given the consistent enrichment of B_atyp_ in CAR T cell-treated patients, we asked whether activated CAR T cells can directly drive the B_atyp_ phenotype. To this end, we collected supernatants from co-cultures in which CD33-28z or CD19-28z (negative control) CAR T cells were cultured with OCI-AML3 cells and B cells, or with B cells alone, and used them to stimulate HD-derived B cells (**Fig. 5F**). After 72 hours, only supernatants from CD33-28z CAR T cell–OCI-AML3–B cell co-cultures, but not from CAR T cell–B cell co-cultures lacking AML cells, induced the B_atyp_ program, with upregulation of PD-L1 and T-Bet (**Fig. 5G**). Baseline expression of these markers in unstimulated B cells (day 0) was low, confirming that their upregulation was induced by the supernatant rather than reflecting a pre-existing phenotype (**Extended Data Fig. 7H**).

To delineate the involved signaling pathways, we next repeated this experiment in the presence of the TLR4 inhibitor CLI-095, the TLR7–9 antagonist ODN2088, or the JAK1/2 inhibitor ruxolitinib. Ruxolitinib and ODN2088, but not CLI-095, reduced expression of both PD-L1 and T-bet (**Fig. 5H**), indicating that TLR and JAK/STAT signaling are both required to induce the ABC program. Phospho-flow cytometry confirmed that conditioned supernatant induced STAT1 phosphorylation, which ruxolitinib abolished (**Fig. 5I**).

To identify additional ABC inducers, we screened cytokine and TLR agonist combinations in HD-derived B cells (**Fig. 5K**). Necrotic tumor cells, generated by repeated freeze-thaw cycles, were also plated in combination with type I, II or III interferons, to test whether the effect of CAR T cell– conditioned medium was driven by DAMPs released from dying cells.

As previously described^22^, the combination of the TLR7/TLR8 agonist R848 with IFN-γ led to the strongest upregulation of the ABC program, indicated by induction of CD11c, FcRL5, T-bet and TGF-β. Combining R848 with IFN-α or IFN-λ also induced FcRL5, PD-L1, T-bet and TGF-β, albeit more weakly than IFN-γ and without affecting CD11c expression (**Extended Data Fig. 7I,J**). In contrast, neither AML nor ALL lysates induced the ABC program when combined with IFN-α, IFN-γ or IFN-λ (**Fig. 5K**).Yet, freeze-thaw treatment induces primarily necrotic death, whereas CAR T cell– mediated killing induces apoptosis, and the released material may therefore differ in composition and in the extent to which nucleic acids remain protected from degradation.

Intriguingly, treating B cells with recombinant, plate-bound CD70 in combination with IFN-γ also strongly induced PD-L1 and T-bet (**Fig. 5K,L**), as well as TGF-β (**Fig. 5K,M**). CD70 is constitutively expressed on AML blasts^53^ and on activated T and B cells (**Extended Data Fig. 8K,N**). As TLR signaling, CD27 converges on NF-κB through TRAF-dependent pathways, activating NF-κB and JNK via TRAF2, TRAF5 and NIK^54^.

In summary, our data show that CD33-directed CAR T cells can induce an immunosuppressive ABC program in B cells through soluble mediators acting on TLR7–9 and interferon signaling. In addition, we identify CD70–CD27 signaling as a non-canonical input that, together with IFN-γ, induces T-bet in B cells.

### CD180 is a candidate target co-expressed on AML blasts and B_atyp_

As indicated above, at single cell levels, CD180 was the only surface protein we found upregulated on B_atyp_ between HD and AML patients. CD180 is a TLR-related surface molecule expressed on both B cells and myeloid cells^30^. Multiplexed immunofluorescence staining of primary AML samples already indicated expression of CD180 also on CD34^+^ AML Blasts (**Fig. 4B, Extended Data Fig. 6A**).

To examine its expression on AML blasts and to identify additional potential target antigens which are shared between AML and B cells, we analyzed a scRNA-seq dataset comprising 16 AML patients with integrated mutational profiling, allowing clear distinction between healthy and malignant blasts^55^ (**Fig. 6A**). Reanalysis and reclustering of the entire scRNA-seq dataset identified 21 distinct cell populations. Consistent with previous reports and the molecular heterogeneity of AML, we observed six malignant cell populations spanning different differentiation states, including HSC-like, Prog-like, GMP-like, ProMono-like, Mono-like, and cDC-like clusters (**Fig. 6B, Extended Data Fig. 8A,B**). In most cases, AML and HD samples were readily distinguishable (**Fig. 6C**).

**Figure 6:**
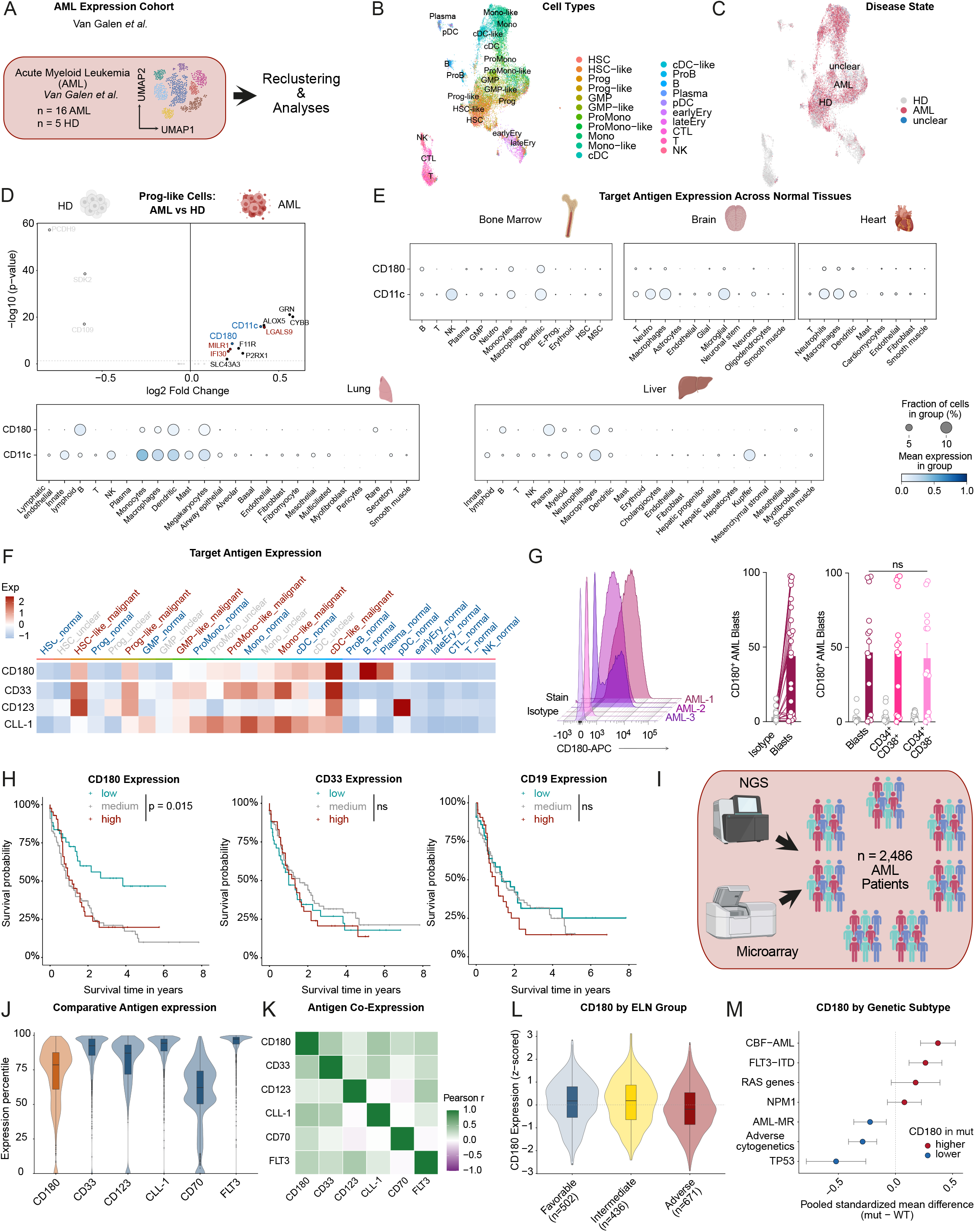
CD180 is expressed on AML blasts and associated with adverse prognosis. (**A**) Overview of the analyzed scRNA-seq dataset from van Galen *et al.*^55^ (n = 21.667 sequenced cells) including integrated mutational profiling to distinguish healthy and malignant AML blasts. (**B**) UMAP embedding showing 21 transcriptionally distinct cell populations, including six malignant clusters (HSC-like, Prog-like, GMP-like, ProMono-like, Mono-like, and cDC-like). (**C**) Visualization of AML and HD samples across the UMAP embedding. (**D**) Volcano plot showing log fold change and bonferroni-adjusted *P* values from DGE between Prog vs Prog-like. Blue: target genes meeting all screening criteria (transmembrane localization, B cell expression >= 6 %, < 2 % expression on T cells, limited off-tumor expression). Red: genes excluded due to off-tumor expression. Black: intracellular or secreted proteins. Grey: genes upregulated in healthy BM controls. (**E**) Comparative scRNA-seq off-target expression screening for candidate antigens CD180 and ITGAX (CD11c) using a cross-organ off-target transcriptomic atlas^56^ (COOTA; ∼ 2.5 million single cells, 11 organs). (**F**) Heatmap showing scaled average expression of CD180 compared to hallmark AML target antigens (CD33, CD123, CLL-1) across healthy and malignant cell clusters. (**G**) Flow cytometric validation of CD180 surface expression on primary AML blasts from patient samples. Left: Representative histograms showing expression of CD180 compared to isotype control stainings. Middle: Quantification of CD180 expression on AML blasts compared to isotype control staining. Data are mean ± s.e.m. from n = 23 AML patients. Right: Quantification of CD180 expression across leukemic subpopulations, demonstrating comparable levels in AML blasts and leukemia-initiating cells (LICs; CD34 CD38 and CD34 CD38 ). Data are mean ± s.e.m from *n* = 13 AML patients. Statistical analysis was performed using ordinary one-way ANOVA with Tukey’s multiple comparison test. (**H**) Kaplan– Meier survival analyses of the TCGA-LAML cohort (*n* = 173 AML patients) comparing overall survival of patients with high, medium, and low expression of CD180 (left), CD33 (middle left), CLL-1 (middle right), and CD19 (right). Statistical significance was calculated with log-rank test. (**I**) Overview of the pooled bulk RNA-sequencing cohort (n = 2,486 patients with AML), combining BEAT-AML 2.0 (phs001657.v3.p1), GSE6891^42^ and GSE146173^43^, GSE37642^75^, GSE68833^76^, GSE232130^77^, GSE199451^78^. (**J**) Expression of the indicated target antigens in the pooled AML cohort. (**K**) Heatmap of pairwise Pearson correlations (r) between the indicated AML antigens. (**L,M**) CD180 expression across European LeukemiaNet (ELN) risk groups (**L**) or molecular subtypes (**M**). For all panels: \**P* < 0.05; \*\**P* < 0.01; \*\*\**P* < 0.001; \*\*\*\**P* < 0.0001; ns, not significant.

We next performed differential gene expression (DGE) analyses between corresponding healthy and malignant populations (e.g., HSC vs. HSC-like, Prog vs. Prog-like), focusing on transmembrane proteins with shared expression between AML and B cells. Among malignant Prog-like cells, only *CD11c* and *CD180* were identified as transmembrane proteins upregulated on AML blasts with shared expression on B cells (**Fig. 6D**). *CD11c* was also upregulated in HSC-like and Mono-like clusters (**Extended Data Fig. 8C–G**).

To predict potential off-target expression, we applied our recently developed single-cell cross-organ off-target transcriptomic atlas (COOTA^6,56^). While *CD11c* and *CD180* exhibited similar immune cell profiles, *CD11c* showed broader expression across multiple tissues, including glial cells in the brain, neutrophils in the heart, and Kupffer cells in the liver. In contrast, CD180 was expressed primarily on immune cells of the myeloid and B cell lineage (**Fig. 6E**).

At the single-cell level, *CD180* expression patterns on AML blasts were comparable to hallmark AML antigens *CD33*, *CD123*, and *CLL-1* (**Fig. 6F**). Flow cytometry confirmed CD180 expression on primary AML blasts (**Fig. 6G**). Importantly, CD180 expression did not differ between AML blasts and leukemia-initiating cells (LICs) residing within the CD34 CD38 and CD34 CD38 compartments^57^ (**Fig. 6G, right**). High *CD180* expression was associated with reduced overall survival in patients with AML. In contrast, most of the known AML-associated target antigens (*CD33, CLL-1*) or B cell antigens (*CD19, CD20, CD22*) did not discriminate poor patient outcomes (**Fig. 6H, Extended Data Fig. 8H).** Only *CD123* showed a similar but less pronounced effect, likely reflecting its known role in promoting leukemic proliferation **(Extended Data Fig. 8H**)^58^.

To validate these expression patterns in independent cohorts and to benchmark CD180 against the established AML targets *CD33*, *CD123*, *CLL-1*, *CD70*, and *FLT3*, we analyzed two additional scRNA-seq cohorts, a five adult patients^59^ and a combined pediatric–adult AML dataset (SCPCP000007; **Extended Data Fig. 8I,J and 8L,M**). In both, *CD180* was highly expressed in the malignant progenitor compartment (**Extended Data Fig. 8K,N**).

For a broader view of translational relevance, we added three bulk RNA-seq cohorts to our initial cohort, yielding a combined 2,486 patients (**Fig. 6I**). *CD180* was highly expressed in primary AML, slightly lower than *CD33*, *CD123*, *CLL-1*, and *FLT3*, but higher than *CD70* (**Fig. 6J**), and did not differ between bone marrow and blood samples (**Extended Data Fig. 9A**). By co-expression analysis, *CD180* was independent of the other target antigens (**Fig. 6K**). Across molecular subtypes, *CD180* was higher in CBF-AML and FLT3-ITD AML, with a trend toward higher expression in RAS- and NPM1-mutated AML, and lower in AML-MR, adverse-cytogenetics, and TP53-mutated AML, consistent with lower expression in ELN-2022 adverse-risk AML (**Fig. 6L,M, Extended Data Fig. 9B**). Finally, *CD180* expression did not differ between first diagnosis and r/r AML , at either the bulk (**Extended Data Fig. 9C,D**) or single cell level (**Extended Data Fig. 9E,F**).

Together, CD180 is a shared surface antigen on AML blasts, including leukemia-initiating cells, and on immunosuppressive B_atyp_, with an off-target profile largely confined to myeloid and B cells, nominating it as a candidate target to address both compartments simultaneously.

### CD180-28z CAR T cells simultaneously lyse AML tumor cells and AML-associated B cells

Our data indicate that B_atyp_ play a pivotal role in shaping an immunosuppressive tumor microenvironment in hematologic malignancies, suppressing T cell responses and accordingly, high abundance correlates with failure to respond to CAR T cell therapy in patients. As we could identify CD180 as unique shared antigen between AML blasts and B_atyp_, we next hypothesized that simultaneous targeting of these two compartments may mediate therapeutic benefits.

First, we generated a novel second-generation anti-CD180 CAR construct containing a CD8 hinge (CD8h) and a CD28 co-stimulatory domain, and expressed it in primary human T cells. Anti-CD33 (CD33-28z) and anti-CD19 (CD19-28z)-CAR constructs with identical configurations served as controls (**Fig. 7A**). CD180 CAR T cells specifically recognized recombinant human CD180 protein, as evidenced by concentration-dependent T cell activation. In contrast, CD19-28z and untransduced T cells (UT) showed no activation (**Fig. 7B**).

**Figure 7:**
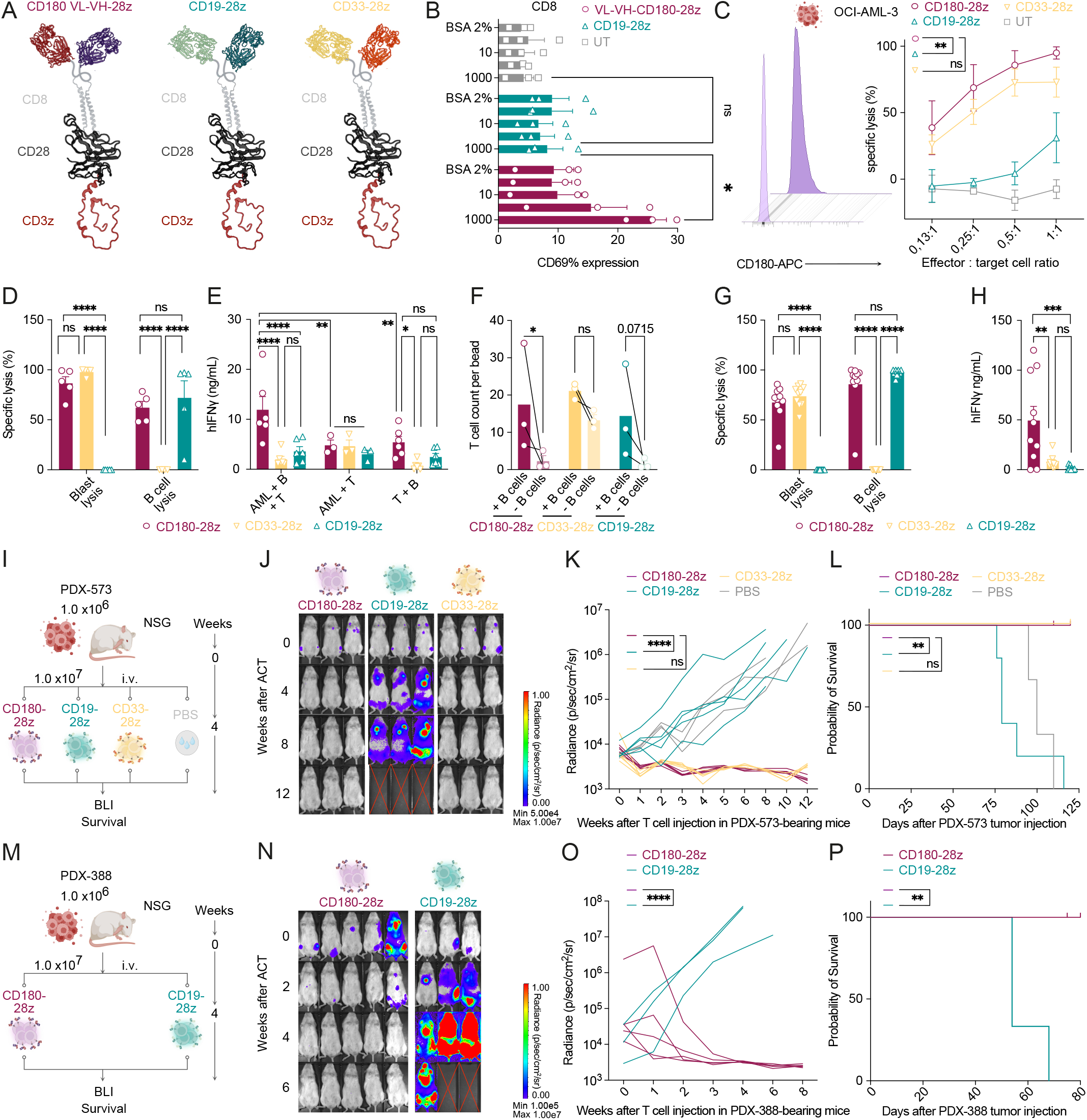
CD180-28z CAR T cells simultaneously target AML blasts and AML-associated B cells. (**A**) Schematic representation of CAR constructs: anti-CD180 (aCD180-VL-VH-28z, maroon), anti-CD19 (CD19-28z, teal), and anti-CD33 (CD33-28z, cantaloupe). (**B**) CD69 expression on CD8 T cells (aCD180-VL-VH-28z, maroon; CD19-28z, teal; untransduced (UT), gray) after incubation on Fc-immobilized recombinant CD180 protein or 2% BSA control. Data are mean ± s.e.m. from *n* = 3 independent donors. (**C**) Left: Flow cytometric detection of CD180 expression on OCI-AML3 AML cells (representative of *n* = 3 independent experiments). Right: Specific lysis of OCI-AML3 AML cells by CD180-28z, CD33-28z, CD19-28z CAR T cells, or UT controls at indicated effector-to-target (E:T) ratios after 48 hours, as determined by bioluminescence imaging (BLI). (**D–F**) Triple co-cultures of OCI-AML3 AML cells and CD19 B cells with the indicated CAR T cells. Data are mean ± s.e.m. from *n* = 5 independent donors. (**D**) Lysis determined by BLI. (**E**) IFNγ secretion measured by ELISA. Data are mean ± s.e.m. from *n* = 6 donors. Statistical analysis by one-way ANOVA with Tukey’s multiple comparison test. (**F**) Quantification of total T cell counts in triple co-cultures by flow cytometry (data are mean ± from *n* = 3 donors). (**G,H**) Specific lysis (**G**) and IFNγ secretion (**H**) in co-cultures of CD180-28z, CD33-28z, and CD19-28z CAR T cells with primary AML samples from BM or pB AML samples. Data are mean ± s.e.m. from *n* = 10 independent donors. (**I–L**) Experimental design (**I**), representative BLI images (**J**), tumor burden quantification (**K**), and Kaplan– Meier survival curves (**L**) of NSG mice engrafted with AML PDX-573 and treated with the indicated number of CAR T cells (*n* = 5 mice per group). (**M–P**) Experimental design (**M**), representative BLI images (**N**), tumor burden quantification (**O**), and Kaplan–Meier survival curves (**P**) of NSG mice engrafted with AML PDX-388 and treated with the indicated number of CAR T cells (*n* = 3–5 mice per group). Unless otherwise indicated, statistical significance was determined using two-way ANOVA with Tukey’s multiple comparison correction or log-rank test for Kaplan-Meier-Curves. For all panels: \**P* < 0.05; \*\**P* < 0.01; \*\*\**P* < 0.001; \*\*\*\**P* < 0.0001; ns, not significant.

To identify the most effective CAR configuration, we compared four CD180 CAR variants differing in scFv orientation and co-stimulatory domains (**Extended Data Fig. 10A**). All constructs were efficiently expressed in primary T cells (**Extended Data Fig. 10B,C**) and lysed MOLM-13 AML cells with similar potency to CD33-28z CAR T cells (**Extended Data Fig. 10D,E**). Given their overall comparable cytotoxicity and proliferative capacity, CD180-VL-VH configuration (hereafter termed CD180-28z CAR T cells) was used for all subsequent experiments.

We then assessed the therapeutic potential of CD180-28z CAR T cells against additional AML cell lines (OCI-AML3, MV4-11, THP-1), all of which expressed CD180. CD180-28z CAR T cells efficiently lysed all AML lines at low effector-to-target (E:T) ratios, performing comparably to CD33-28z CAR T cells (**Fig. 7C, Extended Data Fig. 10E,F**). To test the hypothesized dual-targeting strategy, we performed triple co-cultures of OCI-AML3 cells, CD19 B cells, and CAR T cells. CD180-28z CAR T cells effectively lysed both AML blasts and B cells, similar to CD33-28z and CD19-28z controls on the individual cell types, but secreted higher amounts of IFNγ, indicating superior activation (**Fig. 7D,E**). B cell co-culture enhanced proliferation of CD180-28z CAR T cells, similar to the effect observed with B cell-targeting CD19-28z CAR T cells (**Fig. 7F**).

In co-cultures with primary AML bone marrow samples, CD180-28z CAR T cells again efficiently lysed AML blasts and AML-associated B cells, comparable to CD33-28z and CD19-28z CAR T cells, respectively (**Fig. 7G**), while secreting higher levels of IFNγ (**Fig. 7H**).

We next examined the efficacy of CD180-28z CAR T cells *in vivo*. Mv4-11 (**Extended Data Fig. 11A-D**), THP-1 (**Extended Data Fig. 11E-H)**, and MOLM-13 (**Extended Data Fig. 11I-L)** AML cell lines expressing firefly luciferase were injected into NSG mice. Following CAR T cell treatment, CD180-28z CAR T cells effectively controlled tumor growth across all xenograft models and prolonged overall survival (**Extended Data Fig. 11B-L**). In contrast, CD19-28z CAR T cells, which were used as a non-binding negative control, failed to control tumor progression, and mice rapidly reached humane endpoints.

We further tested CD180-28z CAR T cells in two cytogenetically distinct AML patient-derived xenograft (PDX) models (PDX-573 and PDX-388; **Fig. 7I,M**). Of note, PDX-388 was derived from a patient with r/r AML and therefore allows us to assess the efficacy of CD180-28z CAR T cells in relapsed disease. After leukemia establishment, mice were treated with CAR T cells four weeks post-engraftment. In both PDX models, CD180-28z CAR T cells rapidly eliminated AML blasts, leading to complete tumor clearance in all treated animals (CR 5/5; **Fig. 7J–L, 7N–P**).

In summary, CD180-28z CAR T cells effectively target both AML blasts and AML-associated B cells and demonstrate potent anti-tumor activity *in vitro* and i*n vivo*.

### CD180-28z CAR T cells spare healthy hematopoietic stem and progenitor cells

In addition to limited efficacy, on-target off-tumor toxicity remains a major obstacle in AML-targeted immunotherapies. In our scRNA-seq–based off-tumor expression mapping, CD180 was predominantly expressed on myeloid and B cells (**Fig. 6E**). To further characterize its developmental expression pattern, we performed Slingshot trajectory analyses using the AML dataset from von Galen *et al.*^55^, which includes all stages of hematopoietic and immune cell development—from hematopoietic stem cells (HSCs) to terminally differentiated plasma cells (**Extended Data Fig. 12A,D**).

CD180 expression followed a gradient pattern, increasing along the B cell lineage, with highest levels on mature B and plasma cells but minimal expression on healthy HSCs (**Extended Data Fig. 12B,C**). Similarly, along the myeloid trajectory, CD180 expression rose from progenitor stages toward differentiated monocytic (Mono) and conventional dendritic cells (cDCs), again with negligible expression on HSCs (**Extended Data Fig. 12E,F**).

At the protein level, flow cytometry confirmed minimal CD180 expression on healthy hematopoietic stem and progenitor cells (HSPCs), whereas CD33 was strongly expressed (**Extended Data Fig. 13A,B**). When co-cultured with healthy HSPCs, CD180-28z CAR T cells neither induced HSPC lysis nor secreted high levels of pro-inflammatory cytokines. In contrast, CD33-28z CAR T cells caused marked HSPC lysis and robust IFNγ secretion (**Extended Data Fig. 13C,D**).

We next examined CD180 on healthy immune cells. Consistent with the scRNA-seq data, CD180 was highly expressed on healthy B cells and on CD14 CD16 classical (Mo), CD14 CD16 intermediate (Int Mo), and CD14 CD16 non-classical monocytes (NC; **Extended Data Fig. 13E**). In contrast to the scRNA-Seq data, protein validation did not reveal high expression of CD180 expression on plasma cells of healthy donors (**Extended Data Fig. 13F**). In co-cultures with HD-derived PBMCs, CD180-28z CAR T cells, like CD33-28z CAR T cells, depleted classical monocytes, and CD19 B cells were reduced by both CD180-28z and CD19-28z CAR T cells. T and NK cell numbers were not reduced (**Extended Data Fig. 13G**).

To assess the risk of serious on-target/off-tumor toxicity -in particular severe hyperinflammation, given that CD180 belongs to the TLR family^30^ -we used a PBMC-humanized mouse model, co-injecting 20 million HD-derived PBMCs with high numbers of CD180-28z, CD33-28z, or CD19-28z CAR T cells (**Extended Data Fig. 13H**). No weight loss, a sensitive parameter of toxicity, was observed in any group (**Extended Data Fig. 13I**). Longitudinal serum cytokine profiling showed the highest IFN-γ secretion by CD180-28z CAR T cells at day 3, whereas at later timepoints serum IFN-γ was higher in CD19-28z–treated mice. All other cytokines were only moderately elevated, without differences between products (**Extended Data Fig. 13J**). Two weeks after infusion, analysis of spleen and bone marrow showed complete B cell depletion by CD180-28z and CD19-28z, but not CD33-28z, CAR T cells (**Extended Data Fig. 13K**). CAR T cell expansion did not differ between products in either organ (**Extended Data Fig. 13L**). CD19-28z CAR T cells had the highest expression of PD- 1 and CD25 (**Extended Data Fig. 13M**). Activation markers on all T cells (gated on CAR-negative), were not elevated in mice treated with CD180-28z CAR T cells, yet CD25 expression was moderately higher in mice injected with CD33-28z or CD19-28z CAR T cells (**Extended Data Fig. 13N**).

Together, CD180-28z CAR T cells spared HSPCs and did not trigger hyperinflammation, while sharing the expected B cell– and monocyte-directed on-target activity of CD180 targeting.

### CD180 CAR T cells deplete B_atyp_ and harness B cell co-stimulation for anti-leukemic efficacy *in vivo*

While CD180-28z CAR T cells depleted conventional CD19 B cells both *in vitro* and *in vivo*, and eliminated AML-associated CD19 B cells in primary co-cultures, it remained unclear if they can specifically deplete B_atyp_, and whether such CAR T cells are protected from B_atyp-_mediated suppression.

Therefore, we polarized HD-derived B cells into B_atyp_ by co-culture with OCI-AML3 cells (see also **Fig. 2D**). After 7 days, polarized B cells were co-cultured with either CD180-28z or CD33-28z CAR T cells (**Fig. 8A**). CD180-28z, but not CD33-28z, CAR T cells eliminated the polarized B_atyp_ (**Fig. 8B**). To confirm this in a primary setting, we polarized B cells with primary AML blasts (see also **Fig. 4R**). After 5 days, we set up co-cultures of CD180-28z or CD33-28z CAR T cells with primary AML blasts as targets, in the presence of either polarized B_atyp_ or non-polarized B cells (**Fig. 8C**). At a low effector-to-target (E:T) ratio of 0.25:1, the addition of polarized B_atyp_ specifically reduced blast lysis by CD33-28z CAR T cells, whereas lysis by CD180-28z CAR T cells was unaffected (**Fig. 8D**).

**Figure 8:**
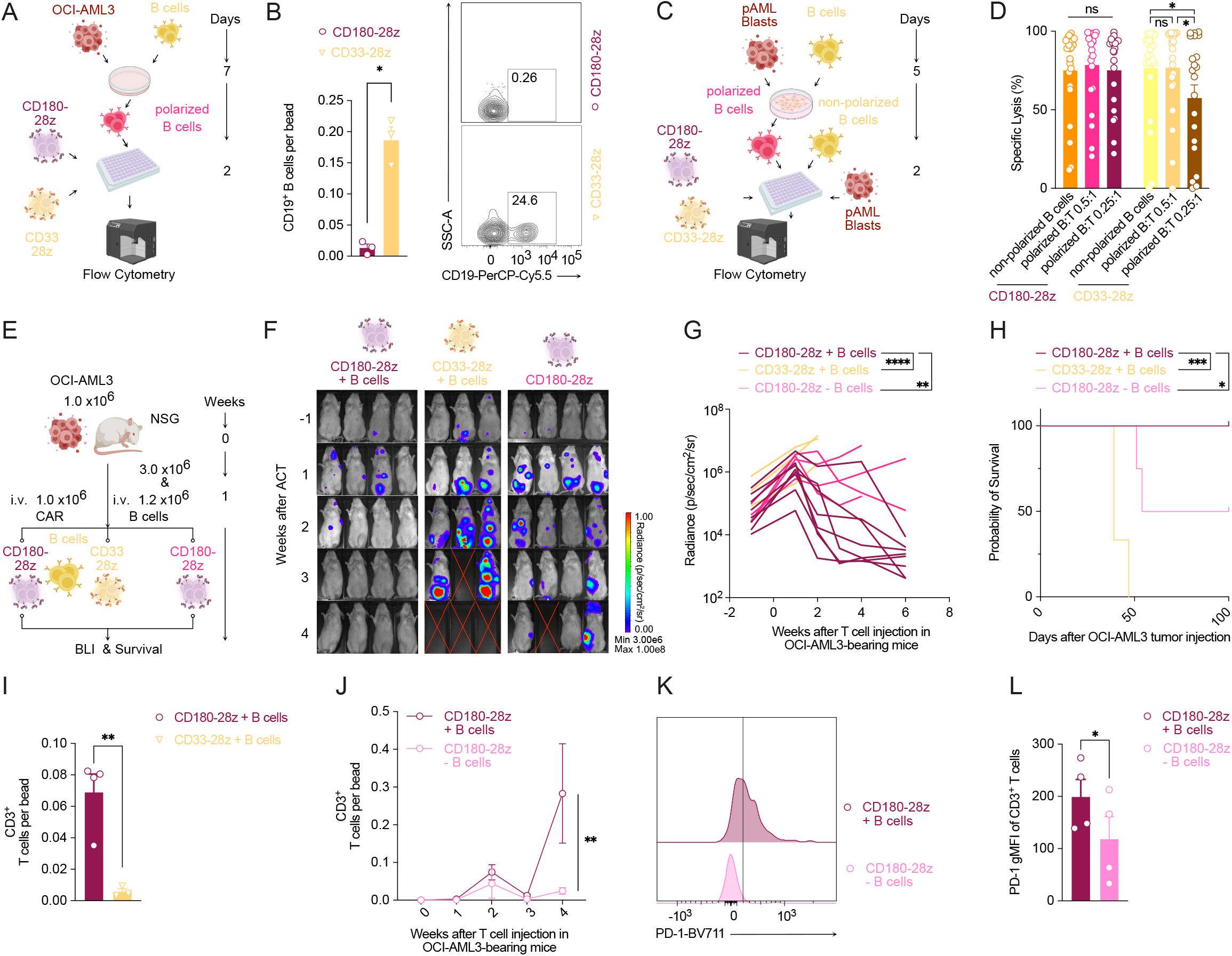
CD180 CAR T cells deplete B_atyp_ and harness B cell co-stimulation for anti-leukemic efficacy *in vivo.* (**A**) Overview of experimental design. (**B**) Left: Counts of AML-polarized B cells quantified by flow cytometry. Right: Representative flow cytometric image. Data are mean ± s.e.m. from *n* = 3 independent donors. Statistical analysis by unpaired *t*-test. (**C**) Experimental workflow: B cells were polarized with pAML blasts for 7 days, then co-cultured with CD33-28z CAR or untransduced T cells (UT) for 48 h at an E:T ratio of 1:2 or 1:4. (**D**) Flow cytometric quantification of CD180-28z or CD33-28z –mediated lysis of pAML blasts. Data are mean ± s.e.m. from n = 22 B cell and T cell donors. (**E–H**) Experimental design (**E**), representative BLI images (**F**), tumor burden quantification (**G**), and Kaplan–Meier survival curves (**H**) of NSG mice engrafted with OCI-AML3 AML tumor cells and CD19 B cells, treated with the indicated CAR T cells (*n* = 3–7 mice per group, pooled from two independent experiments). (**I**) Expansion of CD180-28z or CD33-28z CAR T cells in peripheral blood on day 9 after CAR T cell injection, determined by flow cytometry. Data are mean ± s.e.m. from *n*= 3–4 mice. Statistical analysis by unpaired *t*-test. (**J**) Expansion of CD180-28z CAR T cells in B cell–humanized or non-humanized NSG mice, determined in peripheral blood by flow cytometry. (**K,L**) Quantification of PD-1 expression on CAR T cells as a marker of T cell activation. (**K**) Representative flow cytometric staining. (**L**) Quantification of PD-1 gMFI on CD3^+^ T cells. (**J-L**) Data are mean ± s.e.m. from *n*= 4 mice. Statistical analysis by paired *t*-test. Unless otherwise indicated, statistical significance was determined using two-way ANOVA with Tukey’s multiple comparison correction or log-rank test for Kaplan-Meier-Curves. For all panels: \**P* < 0.05; \*\**P* < 0.01; \*\*\**P* < 0.001; \*\*\*\**P* < 0.0001; ns, not significant.

To analyze *in vivo* efficacy, we next developed a B cell–humanized AML mouse model. OCI-AML3 tumor cells were first injected into NSG mice, followed two weeks later by infusion of HD–derived B cells. Subsequently, B cell–humanized, OCI-AML3 tumor-bearing mice received low-dose CD180-28z or CD33-28z CAR T cells (10 per mouse; **Fig. 8E**). Non-B cell humanized, tumor-bearing mice treated with CD180-28z CAR T cells served as a control. CD180-28z CAR T cells outperformed CD33-28z CAR T cells, achieving near-complete tumor eradication, whereas CD33-28z–treated mice rapidly progressed (**Fig. 8F-H**). While CD33 is homogenously expressed on OCI-AML3 tumor cells, antigen heterogeneity or CAR-mediated internalization of CD33 cannot be excluded^6^. Importantly, co-injection of B cells did not reduce, but rather increased efficacy of CD180-28z CAR T cell therapy, as shown by improved tumor control and prolonged overall survival (**Fig. 8F-H).**

We next quantified CAR T cell expansion and activation to elucidate the underlying mechanism. In B cell–humanized mice, CD180-28z CAR T cells showed greater expansion than CD33-28z CAR T cells, with elevated T cell counts in peripheral blood by day 9 (**Fig. 8I**). Interestingly, CD180-28z CAR T cells expanded more robustly and showed higher activation in B cell–humanized compared with non-humanized mice (**Fig. 8J-L**).

In summary, CD180-28z CAR T cells recognize and eliminate B_atyp_, are resistant to B_atyp_-mediated suppression of blast killing at low effector-to-target ratios, and come with enhanced expansion, activation, and anti-tumor efficacy upon engagement of the B cell compartment in humanized models.

## Discussion

AML has long been viewed largely through the lens of myeloid- or Treg-driven immune suppression, with comparatively little attention paid to the adaptive B cell compartment^60^. Here we show that atypical memory B cells (B_atyp_) constitute an immunosuppressive population within the AML bone marrow, and that they are not merely present but actively induced by the leukemia itself. CAR T cells amplify this induction, thereby reinforcing an immunosuppressive state within the bone marrow microenvironment. Building on earlier transcriptional descriptions of atypical B cell states in AML^28,29^, we combined cross-cohort single cell and flow cytometry analyses with functional studies of patient-derived cells to define how these cells arise, how they impact the efficacy of CAR T cell therapy, and how they can be therapeutically exploited through their shared expression of CD180 with leukemic blasts.

Mechanistically, B_atyp_ are polarized *in situ* by soluble factors from AML blasts. These induce a proinflammatory, immunosuppressive program comprising T-bet, PD-L1, PD-L2 and BTLA, together with TGF-β, alongside anti-apoptotic pathways such as BCL-2. This polarization was further amplified by supernatants from CAR T cell–AML co-cultures. Converging transcriptional, cytokine-perturbation and patient-level analyses implicate interferon signaling and TNF superfamily ligands as the central drivers of B_atyp_ polarization in AML. Through functional experiments, we also identify central roles for TLR7–9 and JAK/STAT signaling in driving this phenotype, as it has been reported in chronic inflammatory conditions. In addition, we identify the CD70–CD27 axis as a previously unrecognized, non-canonical route to induce an atypical B cell reprogramming. Our analyses further revealed that these AML-derived B_atyp_ share key transcriptional pathways with atypical B cell populations described in chronic inflammatory diseases such as HIV, malaria, and SLE^19,21,22,34,40^.

Our analysis highlighted that across multiple patient cohorts, B_atyp_ abundance correlated with clinical non-response to CAR T cell therapy in AML and MM. AML-associated B_atyp_ upregulated inhibitory molecules relative to their healthy donor counterparts and we could demonstrate their ability to potently suppress T cell proliferation and anti-CD33 CAR T cell responses. Although the clinical associations may partly reflect microenvironmental inflammation, our functional findings implicate the adaptive B cell compartment in shaping therapeutic outcomes.

As previously reported, the immunosuppressive B cell state appears to be shaped by environmental cues rather than encoded by a fixed genetic program^14^. This notion is directly mirrored in our data, in which AML blasts and CAR T cells can continuously polarize B cells toward a suppressive B_atyp_ phenotype. Unlike regulatory T cells, whose identity is anchored by the lineage-defining transcription factor FOXP3, these cells lack a comparable hallmark regulator. Our data show that B_atyp_ express high levels of inhibitory ligands and receptors, including PD-L1, PD-L2 and BTLA, and that T cell proliferative capacity can be partially restored by PD-1 blockade. In addition, conditioned supernatant from B_atyp_ alone inhibited T cell proliferation, and this effect was reversed by TGF-β-blockade. Beyond these two mechanisms, B_atyp_ expressed high levels of CD39 and CD73, raising the possibility that adenosine generation contributes to their inhibitory effect. Together our data argue for a broad suppressive impact of B_atyp_, not driven by a single mechanism but rather their combination. This provides a strong rationale for B_atyp_ depletion strategy against individual pathway inhibition.

CD19-targeted CAR T cells have recently shown promising, preliminary efficacy signals in RUNX1::RUNX1T1-mutated AML aberrantly expressing CD19^61^. It is tempting to speculate that some of the activity seen there, may have been driven to some extent by B_atyp_ depletion and B cell co-engagement. In any case, this suggests that B cell depletion is clinically manageable in AML and prompted us to herein explore a dual-targeting approach. This became possible, by the unique expression pattern of CD180, which showed high expression on both AML-associated B_atyp_ and AML blasts. Using our targeting approach, we demonstrate that CD180-directed CAR T cells efficiently deplete B cells — both B_atyp_ and their healthy counterparts — *in vitro* and *in vivo* and that they are resistant to B_atyp_-mediated suppression, especially at low E:T ratios. Notably, beyond merely depleting an immunosuppressive B cell population, in line with recent reports from other groups^62^, we observed a benefit of CAR-B cell interactions itself for T cell performance.

Overall, a direct and expected consequence of CD180-targeted immunotherapies is B cell aplasia. While clinically relevant, such aplasia is a well-precedented and manageable toxicity of B cell– directed cellular therapy: it can be supported by immunoglobulin replacement, and long-term follow-up indicates that patients can tolerate a near-absence of circulating B cells for up to a decade without prohibitive complications^63^.

Beyond the B cell compartment, CD180 is expressed, albeit at lower levels, on macrophages, dendritic cells and megakaryocytes. This off-tumor expression raises the possibility of sustained myeloid depletion and, potentially, of serious infectious complications^64^. This risk is not unique to CD180. Given the myeloid origin of AML, nearly all established AML antigens are also expressed on healthy myeloid cells, with few exceptions such as CD7 or CD56, which themselves carry substantial toxicities, as recently illustrated for anti-CD7 CAR T cells in the clinic^65,66^. Against the dismal prognosis of r/r AML and the considerable morbidity of allo-SCT, the true therapeutic index of CD180-directed therapies will ultimately need to be established in carefully designed clinical trials^67^. Such clinical evaluation is, in fact, already beginning: a first trial of CD180-directed CAR T cells for r/r CD180-positive hematologic malignancies has recently been registered (NCT07106749). To our knowledge, however, no results, preclinical rationale, or evidence of potency from this trial have been reported, and a biological rationale for CD180-directed therapy had not previously been described, rendering its interpretation speculative at present.

Besides CD180, our characterization nominates several alternative or complementary antigens: combining established AML targets (CD33, CD123) with B cell antigens (CD19, CD20, CD22, CD79a) is conceptually attractive but most likely constrained by the sustained hematotoxicity expected from high CD33/CD123 expression on HSPCs^4,68^. In contrast, CD180 combines minimal toxicity towards healthy HSPCs with activity against B_atyp_. As confirmed by our studies, beyond CD180, we provide further candidate antigens, including CD79a, CD11c (also present on primary AML blasts; **Fig. 7D**), FcRL5, and TLR10, whose evaluation will require dedicated follow-up studies.

Our approach builds on, rather than departs from, an established body of work that seeks to co-target malignant cells together with immunosuppressive elements of the tumor microenvironment. CAR T cells directed against immunosuppressive myeloid populations, for instance, have been used to relieve tumor-induced immune suppression and to simultaneously engage malignant and regulatory immune compartments^69,70^. Most recently, this principle has been extended to a single shared antigen: GPNMB, co-expressed on glioblastoma cells and immunosuppressive tumor-associated macrophages, enables anti-GPNMB CAR T cells to couple tumor control with microenvironmental reprogramming through one target^71^. Notably, these efforts have centered predominantly on the myeloid compartment. Our study extends this established dual-compartment framework to the B cell compartment, identifying CD180 as an antigen co-expressed on AML blasts and immunosuppressive B_atyp_, and adds a further dimension, in that B cell engagement not only removes a suppressive population but can also provide co-stimulatory support to CAR T cells^62^.

In conclusion, our study identifies atypical memory B cells as a major immunosuppressive population in AML and characterizes CD180 as a dual-function target that couples the depletion of a suppressive B cell compartment with enhanced CAR T cell activity. By showing that B cell engagement can act as a physiological amplifier of CAR T cell responses, our findings offer a mechanistic and translational framework for exploiting B–T cell interactions to counter immune resistance, with potential relevance across hematologic malignancies and, possibly, solid tumors.

## Acknowledgement and Funding

The work leading to this publication was supported by the PRIME programme of the German Academic Exchange Service (DAAD) with funds from the German Federal Ministry of Education and Research (BMBF). This study was further supported by the Förderprogramm für Forschung und Lehre der Medizinischen Fakultät der LMU (grant number 1138 to A.G.), the DKTK School of Oncology (to A.G.), the Bavarian Cancer Research Center (BZKF; A. Gottschlich and TANGO to S.K.), the Deutsche Forschungsgemeinschaft (DFG, grant number GO 3823/1-1 to A.G.; grant numbers KO5055-2-1, KO5055/3-1 to S.K., TRR332/B01 to S.F.-W. and C.K), the Else Kröner-Fresenius-Stiftung (grant numbers 2023_EKEA.19 to A. Gottschlich and IOLIN to A.G., A.O. and S.K.), German Cancer Aid (grant number: 70115514 to A. Gottschlich and AvantCAR.de, 70117182 to S. Kobold), the Deutsche José Carreras Leukämie-Stiftung (grant number DJCLS 01 R/2025 to A.G. and S.K.) the international doctoral program ‘i-Target: immunotargeting of cancer’ (funded by the Elite Network of Bavaria; to S.K.), the Melanoma Research Alliance (grant number 409510 to S.K.), Marie Sklodowska-Curie Training Network for Optimizing Adoptive T Cell Therapy of Cancer (funded by the Horizon 2020 programme of the European Union; grant 955575 to S.K.), Marie Sklodowska-Curie Training Network for Tracking and Controlling Therapeutic Immune Cells in Cancer (funded by the Horizon Programme of the EU; grant 101168810 to S.K.), the Wilhelm-Sander-Stiftung (to S.K.), Ernst Jung Stiftung (to S.K.), Institutional Strategy LMUexcellent of LMU Munich (within the framework of the German Excellence Initiative; to S. Kobold), the Go-Bio-Initiative (to S.K.), the m4-Award of the Bavarian Ministry for Economic Affairs (to S.K.), Bundesministerium für Bildung und Forschung (to S.K.), the EUROSTAR-Programm (to S.K.), and the European Research Council (Starting Grant 756017, PoC Grant 101100460 and CoG 101124203 to S.K.), SFB-TRR 338/3 2026 –452881907, by the CRC 1784 (ID 560243407; B07 to S.K.) (both to S.K.), Fritz-Bender Foundation (to S.K.), Hector Foundation (to S.K. and C.K.), Bavarian Research Foundation (BAYCELLATOR to S.K.), the Monika-Kutzner Foundation (S.K.), the Bruno and Helene Jöster Foundation (360° CAR to S.K.), the Dr. Rurainski Foundation (to S.K.), the Reinhard Frank Foundation (S.F.-W.), the Stihl Foundation (C.K.) and the Constanze and Dr. Brigitte Wegener Foundation (to S.K.). The study was further supported by the Bavarian Transformation and Research Foundation (BAYCELLator AZ1568-22 to S.K., and Scholarship DOK-195-24 to S.D.). J.H. was supported by a doctoral scholarship (Promotionsstipendium) from the Studienstiftung des deutschen Volkes. N.K. is supported by the International Myeloma Society and the Paula and Rodger Riney Foundation Career Development Award. M.T. is funded by the Volkswagen Foundation (project OntoTime). C.M. has received funding from the European Research Council (ERC) under the European Union’s Horizon 2020 research and innovation programme (grant agreement number 866411). V.I., R.G., T.L., A.Ö. and S.M. were supported by a grant from the Förderprogramm fu r Forschung und Lehre (FöFoLe) of the Ludwig Maximilian University (LMU) of Munich. S.S. declares research funding by the Else Kröner-Fresenius Clinician Scientist Program Cancer Immunotherapy and IOLIN, the Munich Clinician Scientist Program (MCSP), the DKTK School of Oncology, the SFB-TRR 338 (start-up funding) and BZKF (start-up funding 2025). M.R., M.K., M.M. and K.R. received funding from the European Union (CERTAINTY, Grant Agreement 101136379). Views and opinions expressed are those of the author(s) only and do not necessarily reflect those of the European Union or the Health and Digital Executive Agency. Neither the European Union nor the granting authority can be held responsible for them.

Cytometry data were either obtained in the Core Facility Flow Cytometry (campus site downtown) of the University Hospital, LMU Munich using a BD LSRFortessa™ II, Beckman Coulter CytoFLEX or Cytek Aurora or the iFlow Core Facility of the University Hospital, LMU Munich (INST 409/225-1 FUGG). *In vivo* imaging device was funded by the Deutsche Forschungsgemeinschaft (DFG, German Research Foundation) – INST 409/231-1.

Figure illustrations were created with Biorender.com under a paid subscription. The authors acknowledge the use of large language models (LLMs) to assist in improving the clarity, flow, and grammar of the manuscript. All scientific content, interpretations, and conclusions were conceived, written, and verified by the authors. We acknowledge Life Science Editors for their editing services.

## Author Contributions

A.G. and S.K. conceived and designed the study. A.G., T.A.J., V.I., R.G., L.R., S.N., E.C., F.B., S.F.-W., I.G., D.B., A.O., R.D.M., L.Y.B., C.A.T., G.V.H., S.St., W.d.H., J.H., P.P., M.Tr., B.Ra., P.Z., S.D., P.S., S.So., P.X.M., G.H. and C.Z. performed or assisted with *in vitro* and *in vivo* experiments. A.G., V.K., C.Ra., M.Th., R.B., M.Ra., M.Kre., N.K. and C.M. performed computational and bioinformatic analyses. A.G., V.K., T.A.J., V.I., R.G., L.R., C.Ra., M.Th. and R.B. analysed and interpreted data. S.L., M.Ru. and C.B. performed and interpreted multiplex imaging and histopathological analyses. T.H., K.S., C.Ra., W.F., M.Kru., I.P. and M.M. provided clinical datasets, patient cohorts and primary patient material. J.T.T., D.S., H.L., A.D., A.S.B., S.G., C.K., K.R., P.B. and S.G.A. and provided critical reagents, models or infrastructure. V.H., M.F. and T.M. provided conceptual advice and critical feedback on study design and interpretation. A.G., M.v.B., M.S., S.E. and S.K. supervised the research. A.G., C.M., M.v.B., S.E. and S.K. acquired funding and provided overall project support. A.G., V.K., T.A.J. and S.K. conceptualized the figures and wrote the manuscript. All authors discussed the results, provided critical feedback and approved the final version of the manuscript.

---

**Extended Data Figure 1:**
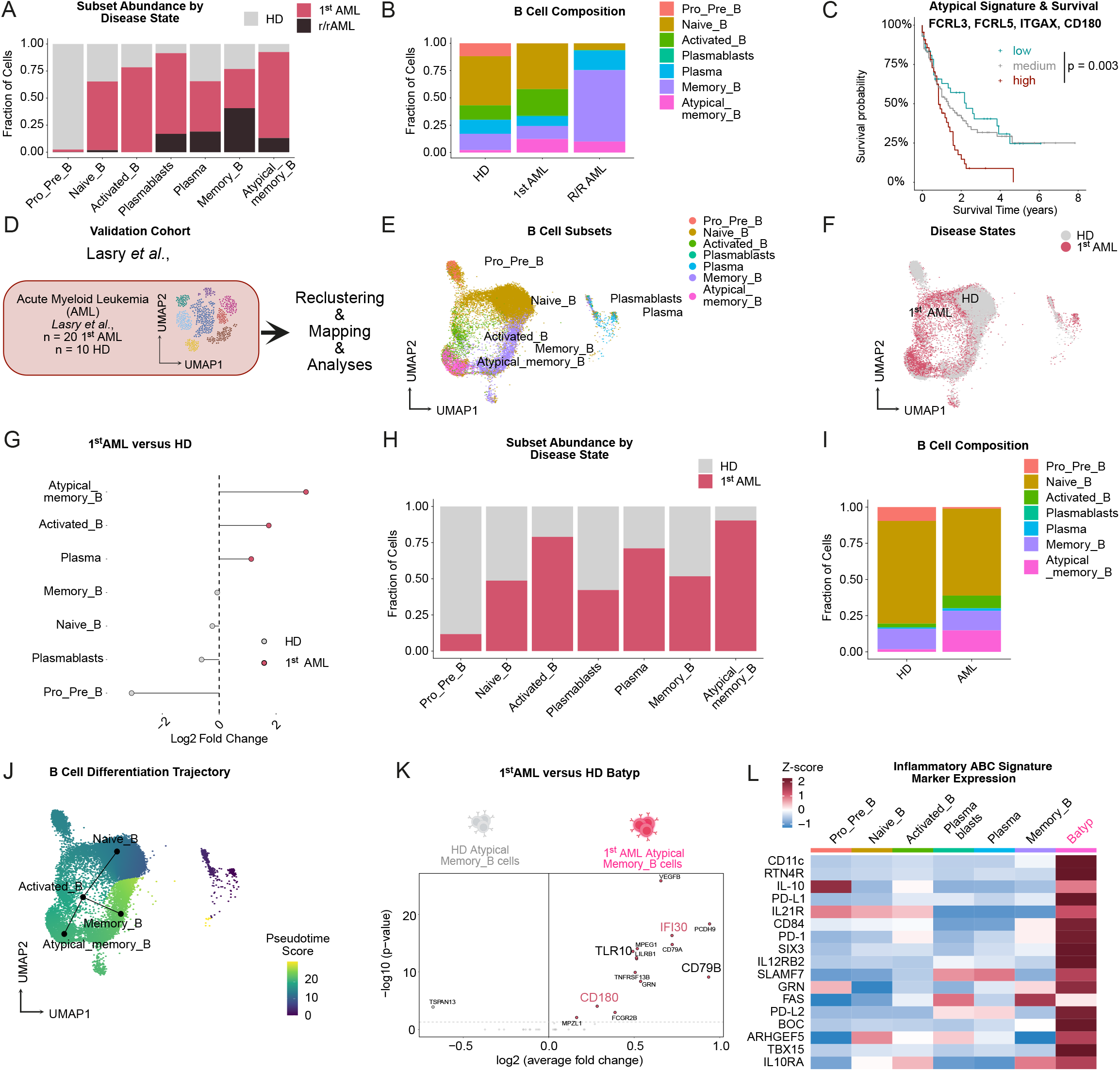
Validation of B_atyp_ enrichment and CD180 expression across independent AML cohorts. (**A,B**) Bar plots showing the distribution of B cell subsets between HD, 1^st^ AML, and r/r AML. (**A**) Frequencies of individual B cell subsets across groups. (**B**) Distribution of B cell clusters across HD, 1^st^ AML, and r/r AML samples. (**C**) Kaplan–Meier survival analysis of patients with AML from the TCGA-LAML cohort (*n* = 173 AML patients) stratified by high, medium, or low expression of a four-gene atypical B cell signature. (**D**) Summary of analyzed patient cohort from Lasry *et al.*^28^, including n = 20 1^st^ AML and n = 10 HD bone marrow samples. (**E**) UMAP embedding of the Lasry *et al.*^28^ dataset after reference mapping to the B cell subsets of the discovery cohort (Guo *et al.*^29^). (**F**) Distribution of identified B cell subsets between HD and AML samples visualized in an UMAP embedding. (**G**) Lollipop plot showing enrichment of B cell subsets in AML compared with HD based on normalized fraction analysis. (**H,I**) Bar plots showing the relative frequencies (**H**) and overall distribution (**I**) of B cell subsets between HD and AML samples. (**J**) Slingshot pseudotime analysis of B cell subsets, with the inferred trajectory from activated B cells toward B_atyp_. (**K**) Volcano plot depicting log fold changes and bonferroni-adjusted P values from DGE between B_atyp_ from HD and AML. **(L)** Heatmap visualization of top-ranked upregulated ABC markers (from Holla et al.^21^) across AML-associated B cell clusters.

**Extended Data Figure 2:**
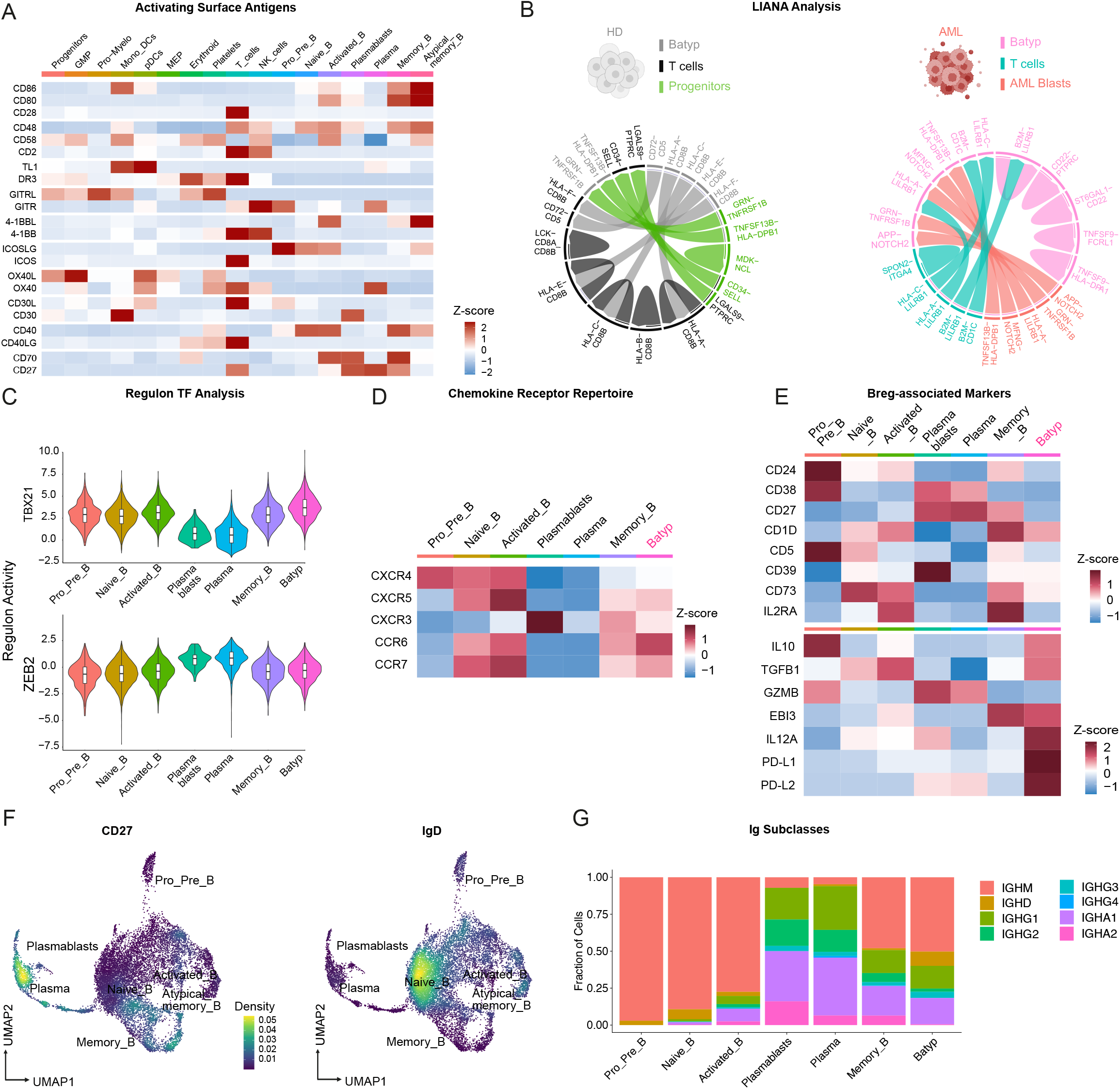
B_atyp_ display an inhibitory, regulatory B cell-like phenotype. **(A)** Heatmap showing scaled average gene expression of hallmark activation pathways. Marker genes were adapted from Chen and Flies^73^. **(B)** Chord diagrams of predicted ligand–receptor interactions between B_atyp_, T cells, and AML blasts in patients with AML (right) versus HD (left). **(C)** Violin plot illustrating DoRothEA^38^ & decoupleR^39^ regulon activity analysis of B cell subclusters. **(D,E)** Heatmap visualization of chemokine receptor repertoire (**D**), Breg-associated surface markers (**E**, top) or functional markers (**E**, bottom) or (**E**) across AML-associated B cell clusters. **(F)** Density plots demonstrating CD27 and IgD expression on distinct B cell subsets visualized within an UMAP embedding. **(G)** Bar plots showing the distribution of Ig subclasses across AML-associated B cell clusters.

**Extended Data Figure 3:**
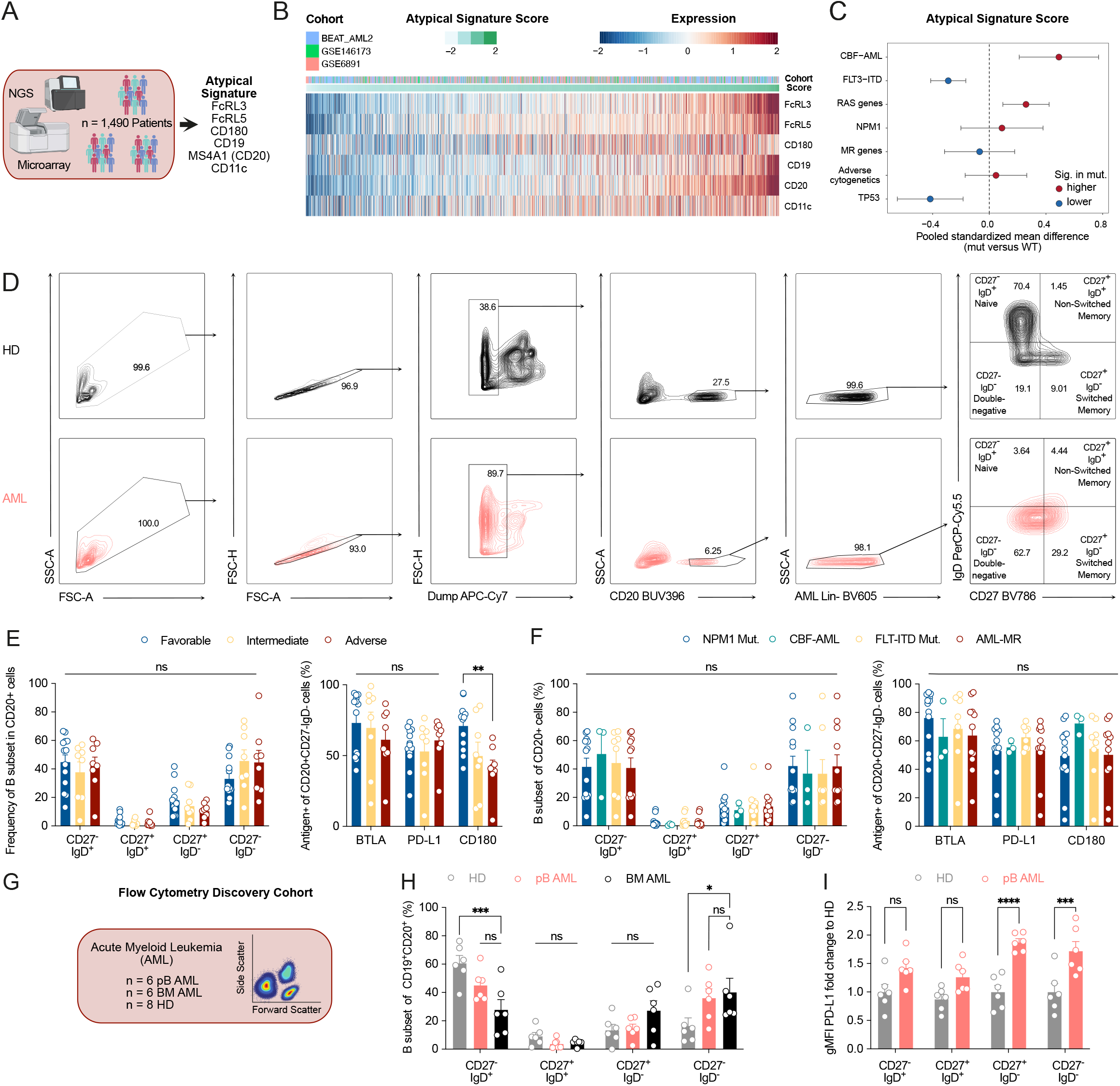
B_atyp_ enrichment and inhibitory phenotype across AML subtypes. **(A)** Overview of the pooled bulk RNA-sequencing cohort (n = 1,490 patients with AML), combining BEAT-AML 2.0 (phs001657.v3.p1), GSE6891^42^ and GSE146173^43^. (**B**) Heatmap visualization of expression of atypical signature (score and single component) across cohorts. (**C**) Forest plot of pooled standardized mean differences (SMD) in the atypical signature score (mutated versus unmutated) across the indicated AML subtypes. (**D**) Overview of the flow cytometric gating strategy used to identify B_atyp_. Top: healthy donors (HD); bottom: patients with AML. (**E,F)** Frequencies of B cell subsets and expression frequencies of inhibitory signaling molecules on B_atyp_ stratified by ELN 2022 classification (**E**) or AML subtype (**F**). Data are mean ± s.e.m; n = 3-13 AML patients. (**G**) Overview of the flow cytometry validation cohort. (**H**) Frequencies of B cell subsets in HD peripheral blood (pB) and in pB AML (pB, red) or BM (black) AML samples as determined by flow cytometry. (**I**) Expression frequencies of inhibitory signaling molecules on individual B cell subsets. (**H,I**) Data are mean ± s.e.m from n = 6 HD, n = 6 pB AML, n = 6 BM AML. Unless otherwise indicated, statistical analysis was performed using two-way ANOVA with Tukey’s multiple comparison test. For all panels: \**P* < 0.05; \*\**P* < 0.01; \*\*\**P* < 0.001; \*\*\*\**P* < 0.0001; ns, not significant.

**Extended Data Figure 4:**
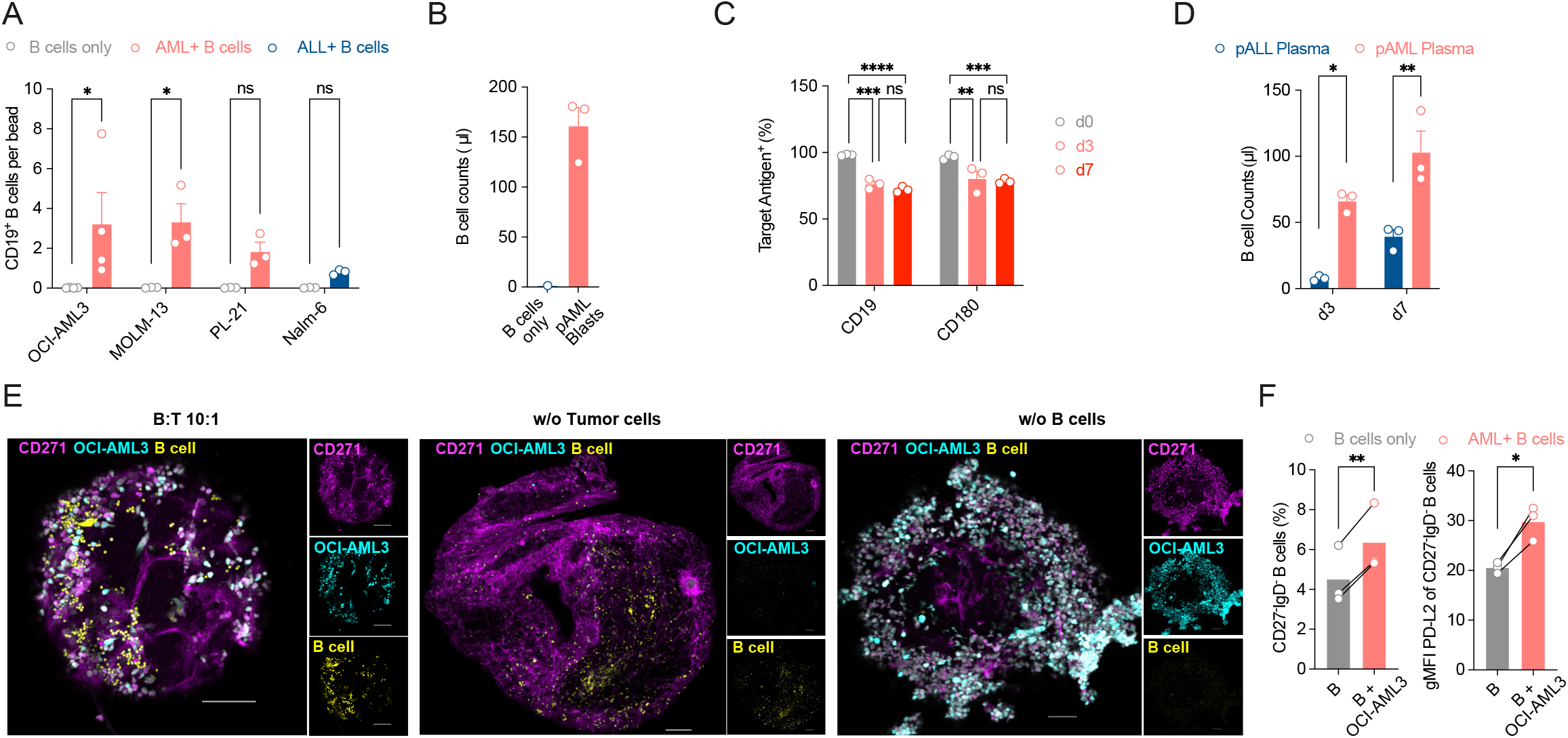
AML blasts support B cell survival and B_atyp_ polarization across model systems. (**A**) Quantification of CD19 B cell counts recovered from co-cultures with the indicated AML-or ALL cell lines. Data represent mean ± s.e.m. from n = 5–6 biological replicates. (**B**) B cell counts after three day culture on MS-5 feeder cells. (**C**) Expression of the indicated markers on B cells in primary AML-B cell co-cultures at the indicated timepoints by flow cytometry. (**D**) B cell counts after 3 (d3) or 7 (d7) days of culture in 50% pAML or pALL bone marrow plasma. **(E)** Representative merged confocal z-stack images of B cell–BMO co-cultures. B cell to tumor cell ratio: 10:1. Single-channel images (right) display BMO mesenchymal stromal cells (CD271, violet), OCI-AML3 tumor cells (turquoise) and B cells (yellow). Scale bar: 100 µm; Control conditions lacking either OCI-AML3 (middle) or B cells (right) are also depicted. Scale bar: 50 µm. (**F**) Flow cytometric quantification of B cell subsets (left) and expression of immune inhibitory cell markers (right) on B cells from engrafted BMO. Data are mean ± s.e.m from n = 3 independent donors. Statistical analysis was performed using paired t-test. Unless otherwise indicated, statistical analysis was performed using two-way ANOVA with Tukey’s multiple comparison test. For all panels: \**P* < 0.05; \*\**P* < 0.01; \*\*\**P* < 0.001; \*\*\*\**P* < 0.0001; ns, not significant.

**Extended Data Figure 5:**
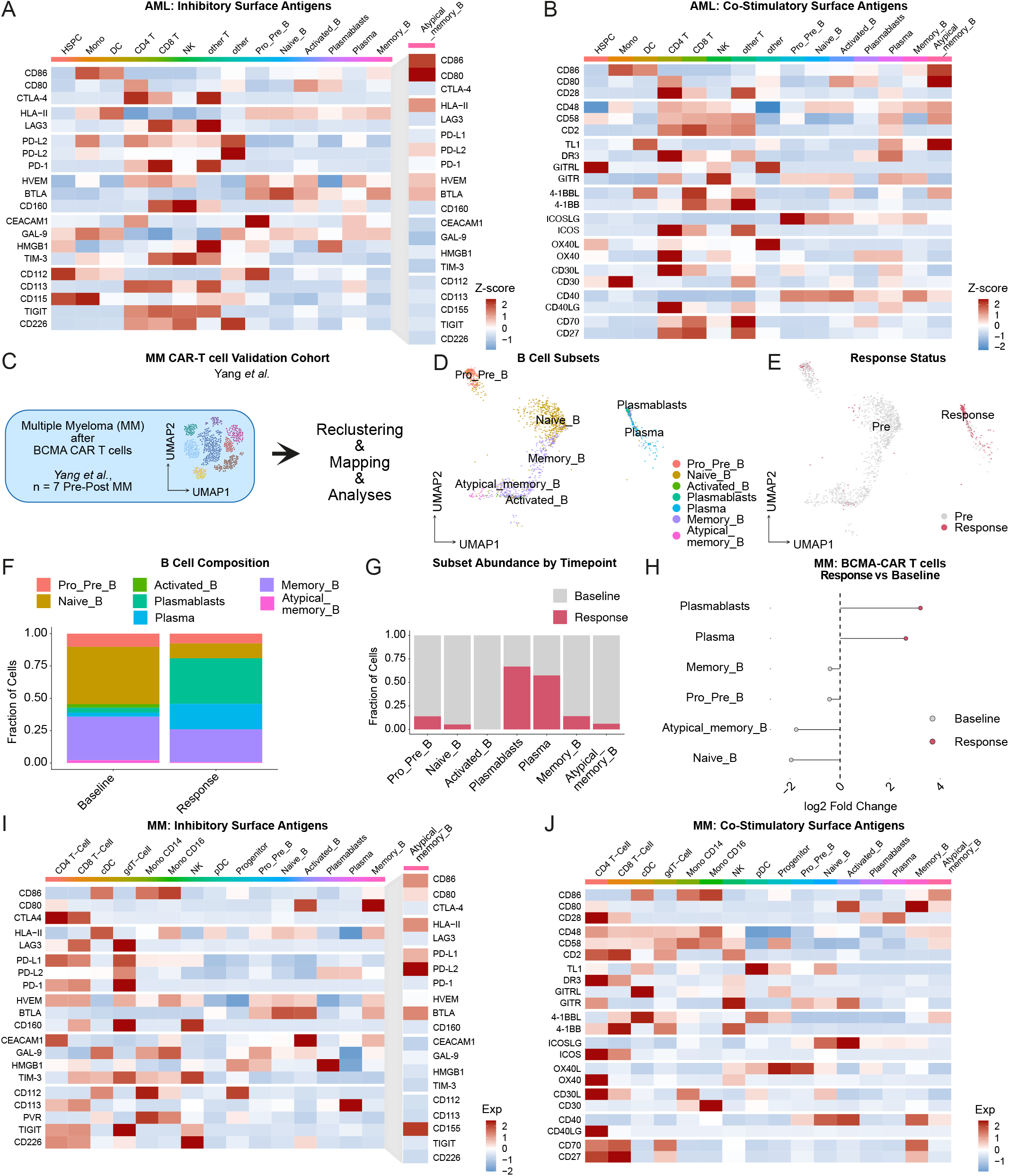
B_atyp_ exhibit conserved immunoinhibitory phenotypes across AML and MM CAR T cell cohorts. (**A,B**) Heatmap visualization of scaled average gene expression of hallmark immune inhibitory (**A**) or activating (**B**) signaling pathways in the scRNA-seq dataset from Bhagwat *et al.* (*NCT03766126*)^10^. Marker selection adapted from Chen and Flies^73^. (**C**) Overview of the analyzed scRNA-seq dataset from Yang *et al.,* comprising MM patients treated with BCMA CAR T cells (*n* = 7 patients with paired biopsies before and after therapy). (**D**) Reference mapping of B cell subsets from Yang *et al.* to the discovery cohort (Guo *et al.*). (**E**) UMAP visualization of MM samples obtained before and after BCMA CAR T cell therapy. (**F**) Bar plots showing relative frequencies of individual B cell subsets before (baseline) and after (response) therapy. (**G**) Distribution of B cell subsets before and after BCMA CAR T cell therapy. (**H**) Lollipop plot showing enrichment of B cell subsets before versus after BCMA CAR T cell therapy based on differential normalized fraction analyses. (**I,J**) Heatmap visualization of scaled average gene expression of hallmark immune inhibitory (**I**) or activating (**J**) signaling pathways in the scRNA-seq dataset from Rade *et al*.^45^.

**Extended Data Figure 6:**
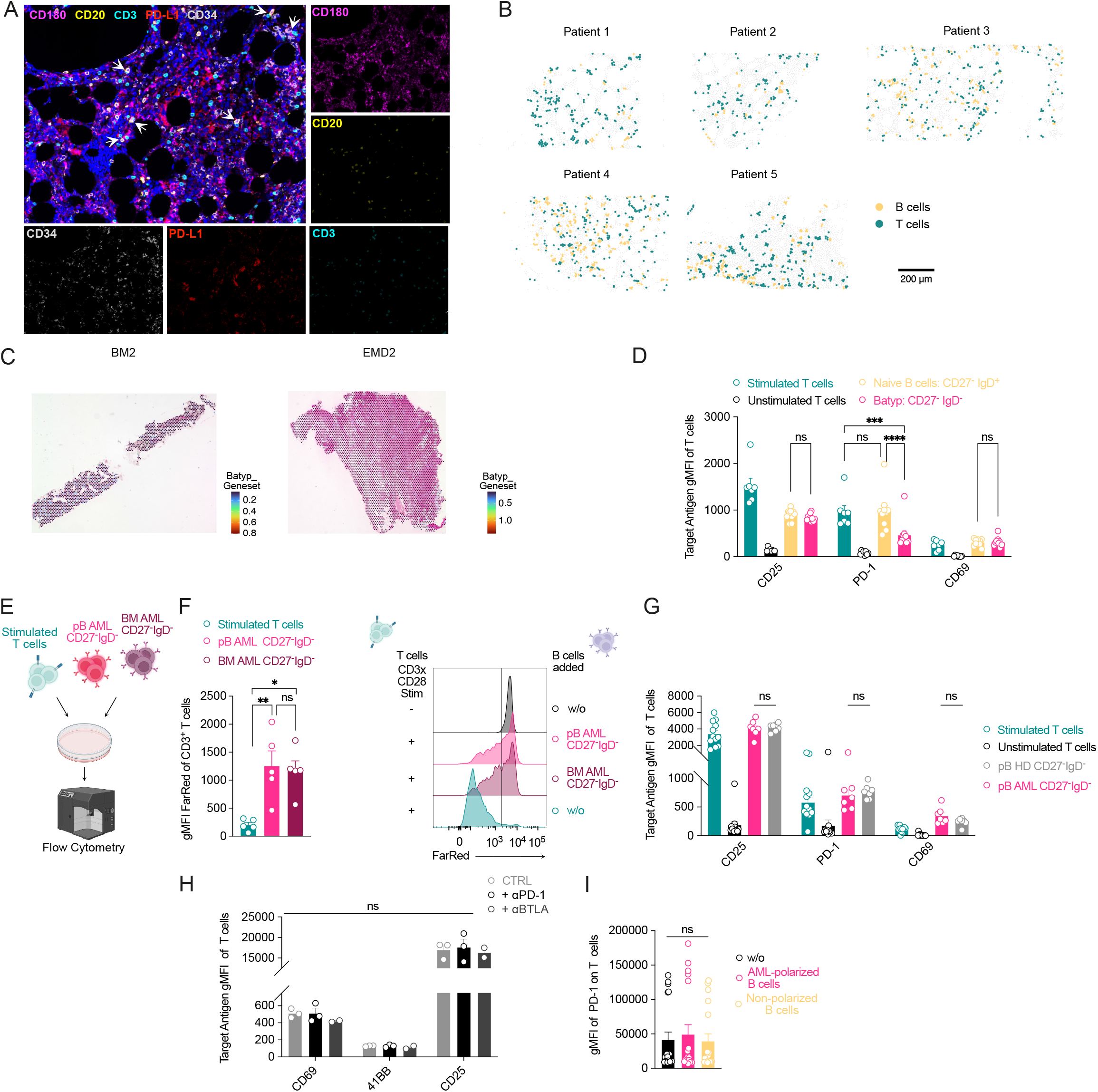
B_atyp_ suppress conventional and CAR T cell responses in AML. **(A**) Representative merged mIF of primary AML bone-marrow biopsies. Single channel images (right) show CD180 (violet), CD20 (yellow, B cells), CD3 (turquoise, T cells), PD-L1 (red), and CD34 (grey, AML blasts). Scale bar, 40 µm. **(B**) Distribution of B cells (yellow) and T cells (turquoise) across mIF slides. (**C**) Distribution of a B_atyp_ geneset across Visium spots in primary bone marrow (BM) or extramedullary disease (EMD). **(D)** Expression of indicated T cell markers after T-B cell co-culture quantified by flow cytometry. Data are mean ± s.e.m. from n = 7–10. (**E**) Experimental workflow: B cells isolated from cryopreserved AML peripheral blood (pB) or bone marrow (BM). (**F**) Left: Dilution of the proliferation dye in stimulated T cells co-cultured with pB-or BM-derived B_atyp_. Data are mean ±s.e.m. from n = 7-12 independent donors. Right: Representative flow cytometry plot. (**G**) Expression of indicated T cell markers after T-B cell co-culture quantified by flow cytometry. Data are mean ± s.e.m. from n = 7-12 independent T cells donors. **(H)** Expression of indicated T cell markers after T-B cell co-culture quantified by flow cytometry. Data are mean ± s.e.m. from 2-3 independent T cells donors. (**I**) Expression of PD-1 on CD33-28z CAR T cells in pAML-polarized B cell co-cultures. Data are mean ± s.e.m. from n = 19 B cell and T cell donors. Unless otherwise indicated, statistical analysis used two-way ANOVA with Tukey’s multiple comparison test. For all panels: \**P* < 0.05; \*\**P* < 0.01; \*\*\**P* < 0.001; \*\*\*\**P* < 0.0001; ns, not significant.

**Extended Data Fig. 7:**
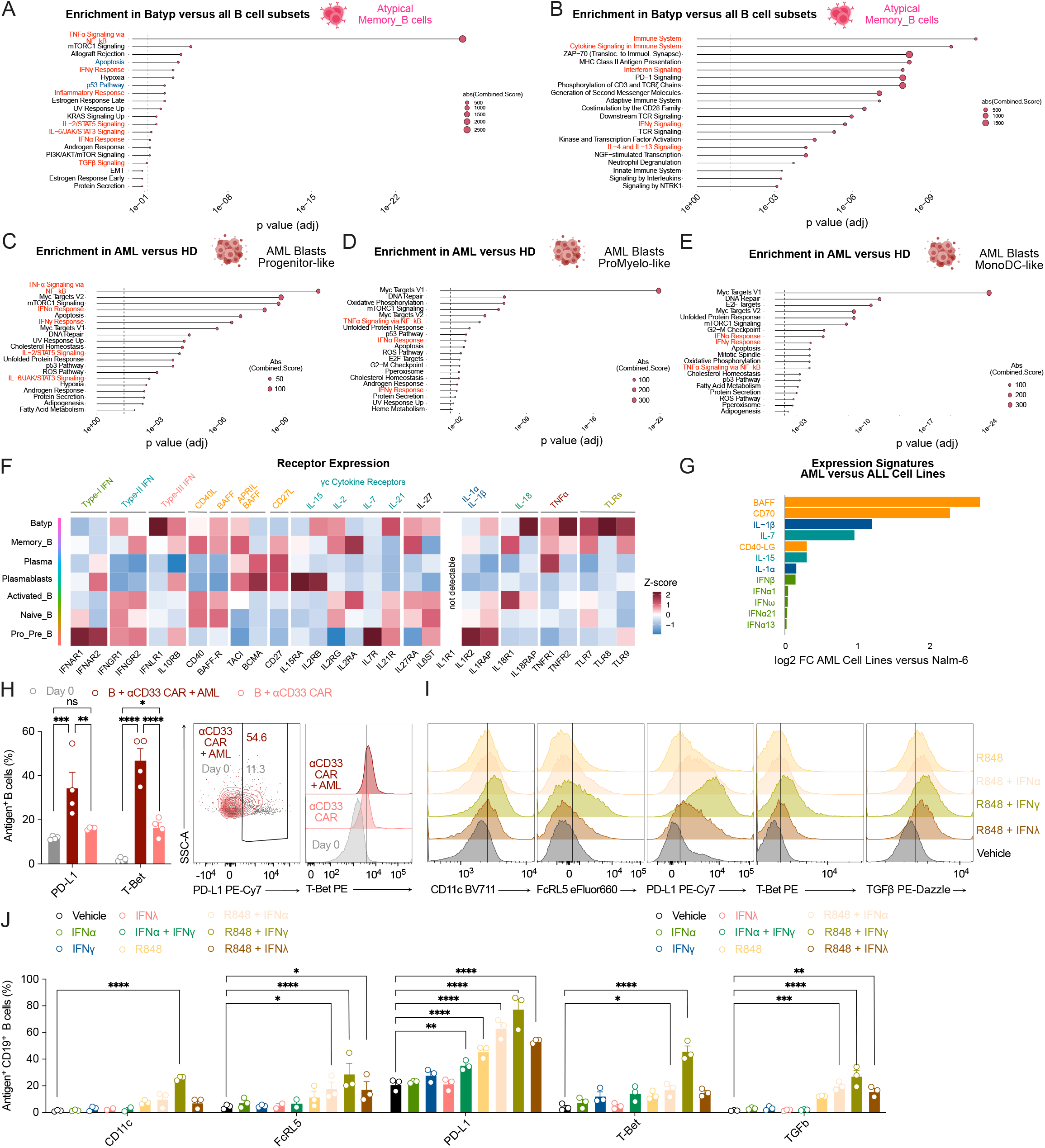
Pathway enrichment in malignant myeloid cells and TLR-and cytokine-driven B_atyp_ polarization. **(A, B)** Pathway enrichment analysis (MSigDB Hallmark, **A**; Reactome, **B**) of pathways enriched in B_atyp_ versus all other B cell subsets of the AML discovery cohort. **(C-E)** Pathway enrichment analysis of the indicated malignant cell populations (Progenitor-like, A; ProMyelo-like, B; MonoDC-like; MSigDB Hallmark) compared to HD. (**F)** Heatmap illustrating expression of indicated cytokine receptors on AML-associated B cell subsets of AML scRNA-seq discovery cohort. (**G**) Log fold-change in expression of indicated markers between pooled AML cell lines (n = 32) and Nalm-6. (**H**) Left: Expression of the indicated markers on CAR–AML supernatant-stimulated B cells (72 h) versus unstimulated B cells at day 0, by flow cytometry. Right: representative flow cytometry plots. Data are mean ± s.e.m. of n = 3–4 B cell–AML–CAR co-cultures. (**I,J**) Expression of the indicated markers on B cells stimulated with indicated TLR-cytokine combinations. (**I**) Representative flow cytometry plot. (**J**) Expression of indicated markers by flow cytometry. Data are mean of n = 3 different B cell donors. Unless otherwise indicated, statistical analysis used two-way ANOVA with Tukey’s multiple comparison test. For all panels: *P < 0.05; **P < 0.01; ***P < 0.001; ****P < 0.0001; ns, not significant.

**Extended Data Figure 8:**
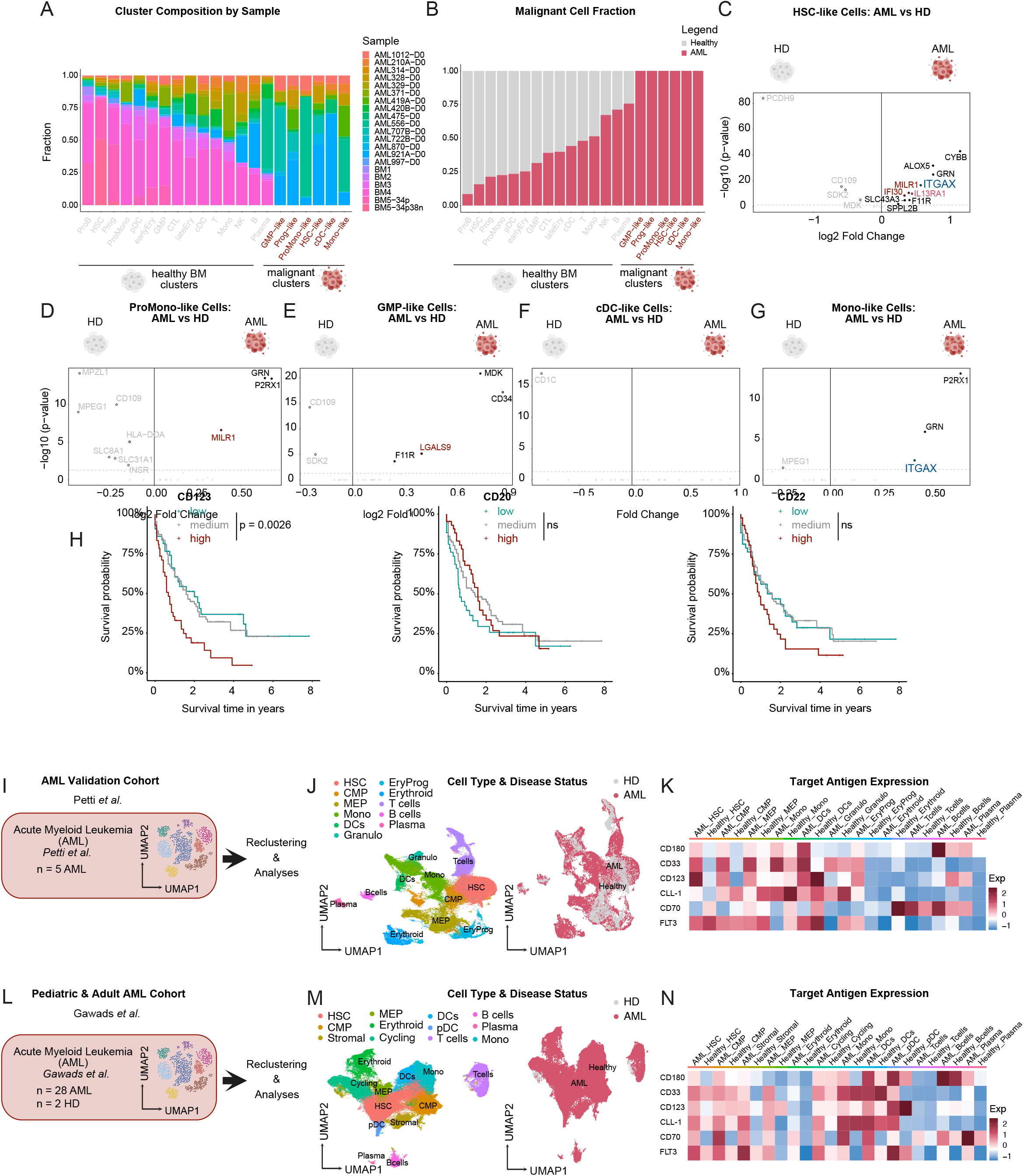
CD180 is expressed on AML blasts and associated with adverse prognosis. (**A,B**) Bar plots showing the distribution of cell clusters across AML patients and healthy controls (**A**) and the relative frequencies of cell subsets between AML and healthy samples (**B**). (**C– G**) Volcano plot showing log fold change and bonferroni-adjusted *P* values from differential gene expression (DGE) analyses between healthy and malignant cell populations: HSC versus HSC-like (**C**), ProMono versus ProMono-like (**D**), GMP versus GMP-like (**E**), cDC versus cDC-like (**F**), Mono versus Mono-like (**G**). Blue: target genes meeting all screening criteria (transmembrane localization, B cell expression >= 6 %, < 2 % expression on T cells, limited off-tumor expression). Red: genes excluded due to off-tumor expression. Black: intracellular or secreted proteins. Grey: genes upregulated in healthy BM controls. (**H**) Kaplan–Meier survival analyses of the TCGA-LAML cohort (*n* = 173) comparing overall survival of patients with high, medium, and low expression of CD123 (left), CD20 (middle) CD22 (right). Statistical significance was calculated with log-rank test. **(I)** Overview of the analyzed scRNA-seq dataset from Petti *et al*.^59^. (**J**) UMAP embedding of immune cell clusters (left) and sample origin (AML versus HD). **(K)** Heatmap showing scaled average expression of CD180 compared to hallmark AML target antigens (CD33, CD123, CLL-1, CD70, FLT3) across healthy and malignant cell clusters. **(L)** Overview of the analyzed scRNA-seq dataset from SCPCP000007. (**M**) UMAP embedding of immune cell clusters (left) and sample origin (AML versus HD). **(N)** Heatmap showing scaled average expression of CD180 compared to hallmark AML target antigens (CD33, CD123, CLL-1, CD70, FLT3) across healthy and malignant cell clusters.

**Extended Data Figure 9:**
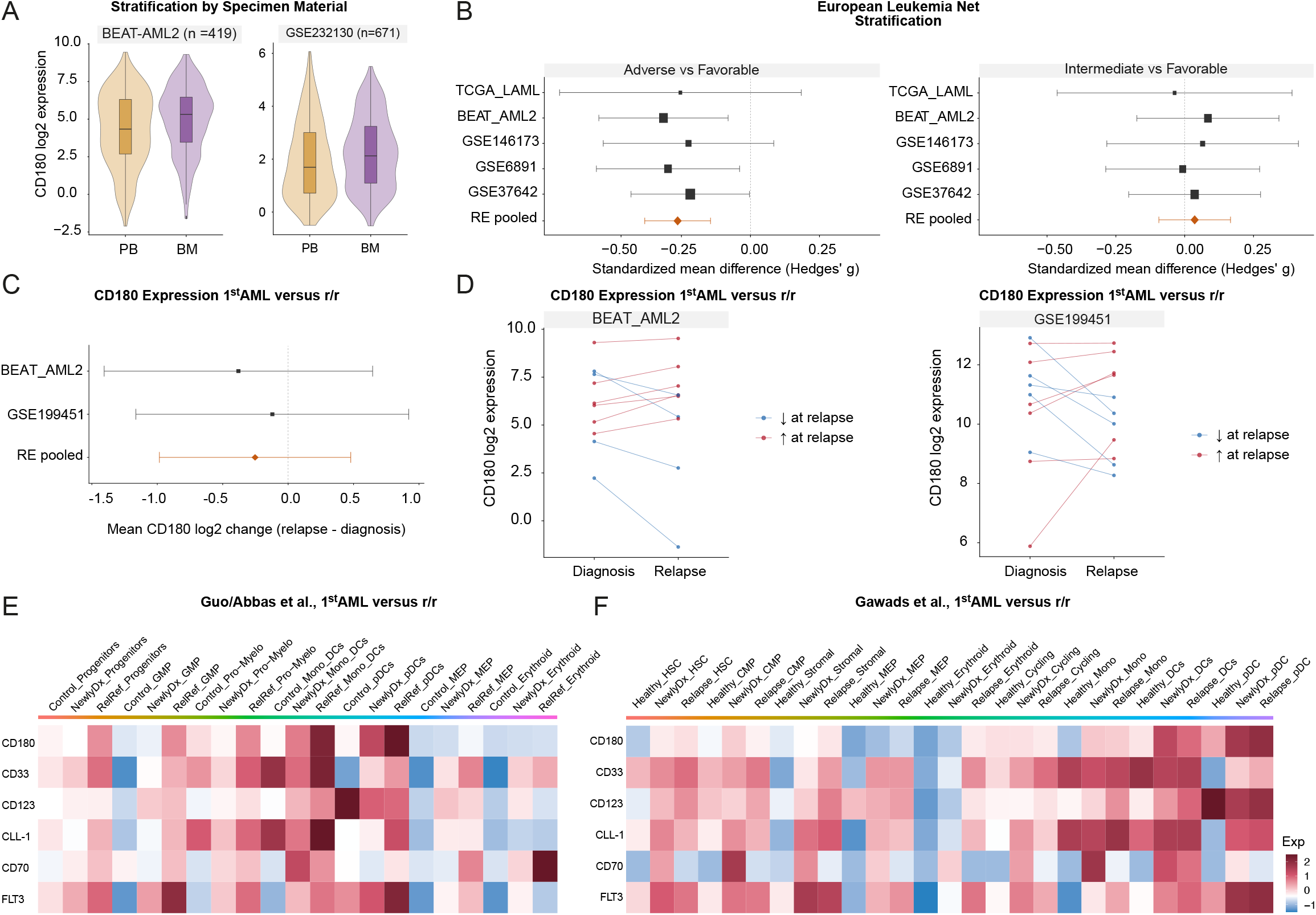
CD180 expression across cohorts and disease stages. (**A**) CD180 expression across peripheral blood (pB) and bone marrow (BM) samples in the indicated bulk sequencing cohorts. (**B**) Forest plot of the standardized mean difference in CD180 expression (Hedges’ g) between ELN adverse and favorable (left) or intermediate and favorable (right) risk groups across cohorts. (**C,D**) Mean difference in CD180 expression between paired first-diagnosis and r/r AML samples. (**E,F**) Heatmap of scaled average CD180 expression compared with hallmark AML target antigens (CD33, CD123, CLL-1, CD70, FLT3) across healthy and malignant cell clusters and disease stages (Guo et al.^29^, **E**; SCPCP000007, **F**).

**Extended Data Figure 10:**
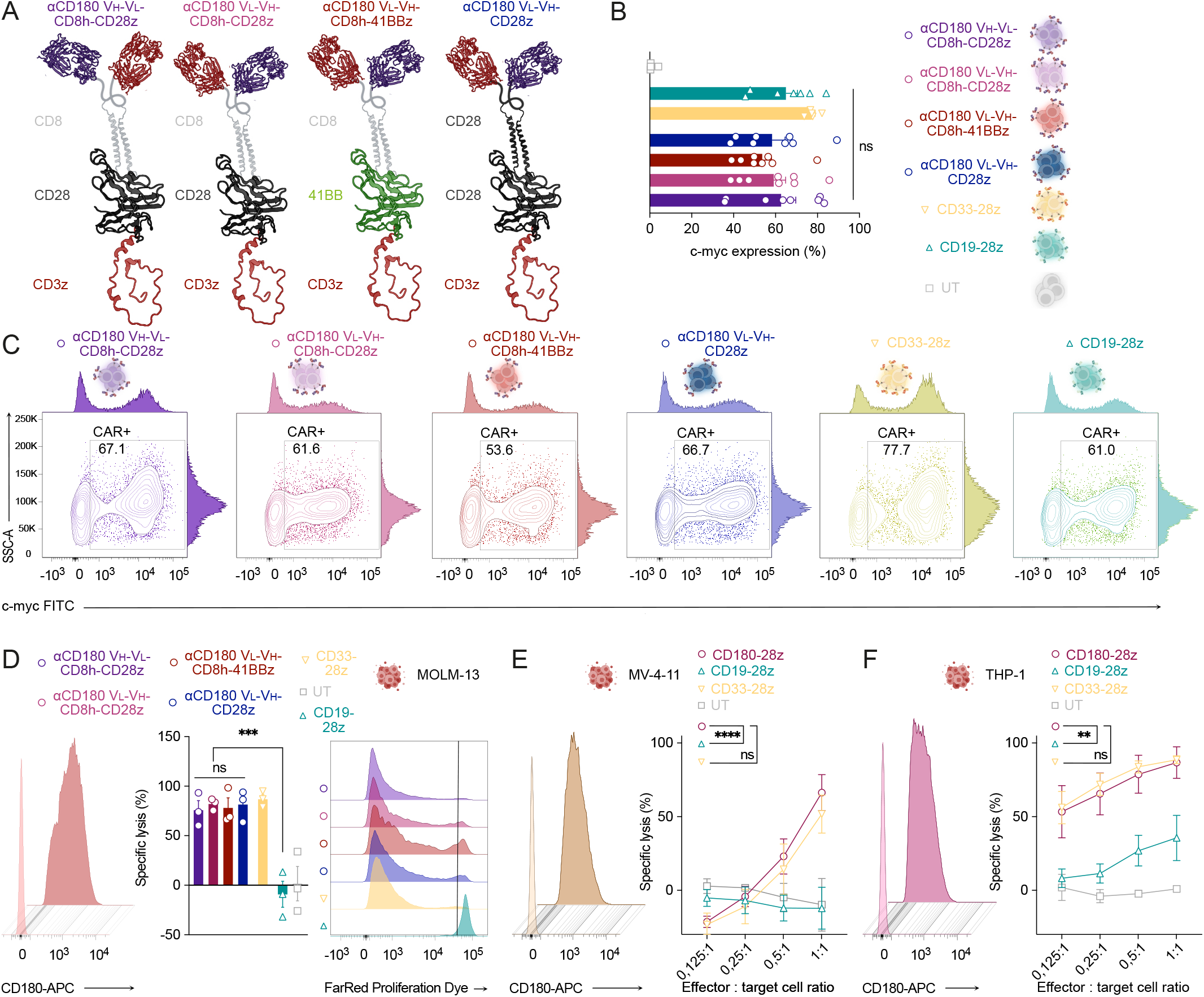
Optimization and *in vitro* characterization of CD180 CAR constructs. (**A**) Schematic overview of the composition of different anti-CD180 CAR configurations. (**B,C**) Summary of transduction efficiencies (**B**) and representative flow cytometric image (**C**) of CD180 CAR constructs compared with CD33 and CD19 control CAR T cells. Data are mean ± s.e.m. from *n* = 8 independent donors. Statistical analysis was performed using one-way ANOVA with Tukey’s multiple comparison correction. (**D**) Left: Flow cytometric detection of CD180 expression on MOLM-13 AML cells. Middle: Specific lysis of MOLM-13 cells by the indicated CAR T cells after 48 hours, measured by BLI. Right: Antigen-specific proliferation of CAR T cells in co-cultures with MOLM-13 cells, determined by FarRed dye dilution after 7 days. Data are mean ± s.e.m. from *n* = 3 independent donors. Statistical analysis was performed using one-way ANOVA with Tukey’s multiple comparison correction. (**E,F**) Left: Flow cytometric detection of CD180 expression on MV4-11 (**E**) and THP-1 (**F**) AML cell lines (representative of *n* = 3 independent experiments). Right: Specific lysis of MV4-11 (**E**) and THP-1 (**F**) cells by CD180-28z, CD33-28z, CD19-28z CAR T cells, or untransduced (UT) controls at indicated effector-to-target (E:T) ratios after 48 hours, determined by BLI. Data are mean ± s.e.m. from *n* = 3 independent donors. Statistical analysis by two-way ANOVA with Tukey’s multiple comparison test. For all panels: \**P* < 0.05; \*\**P* < 0.01; \*\*\**P* < 0.001; \*\*\*\**P* < 0.0001; ns, not significant.

**Extended Data Figure 11:**
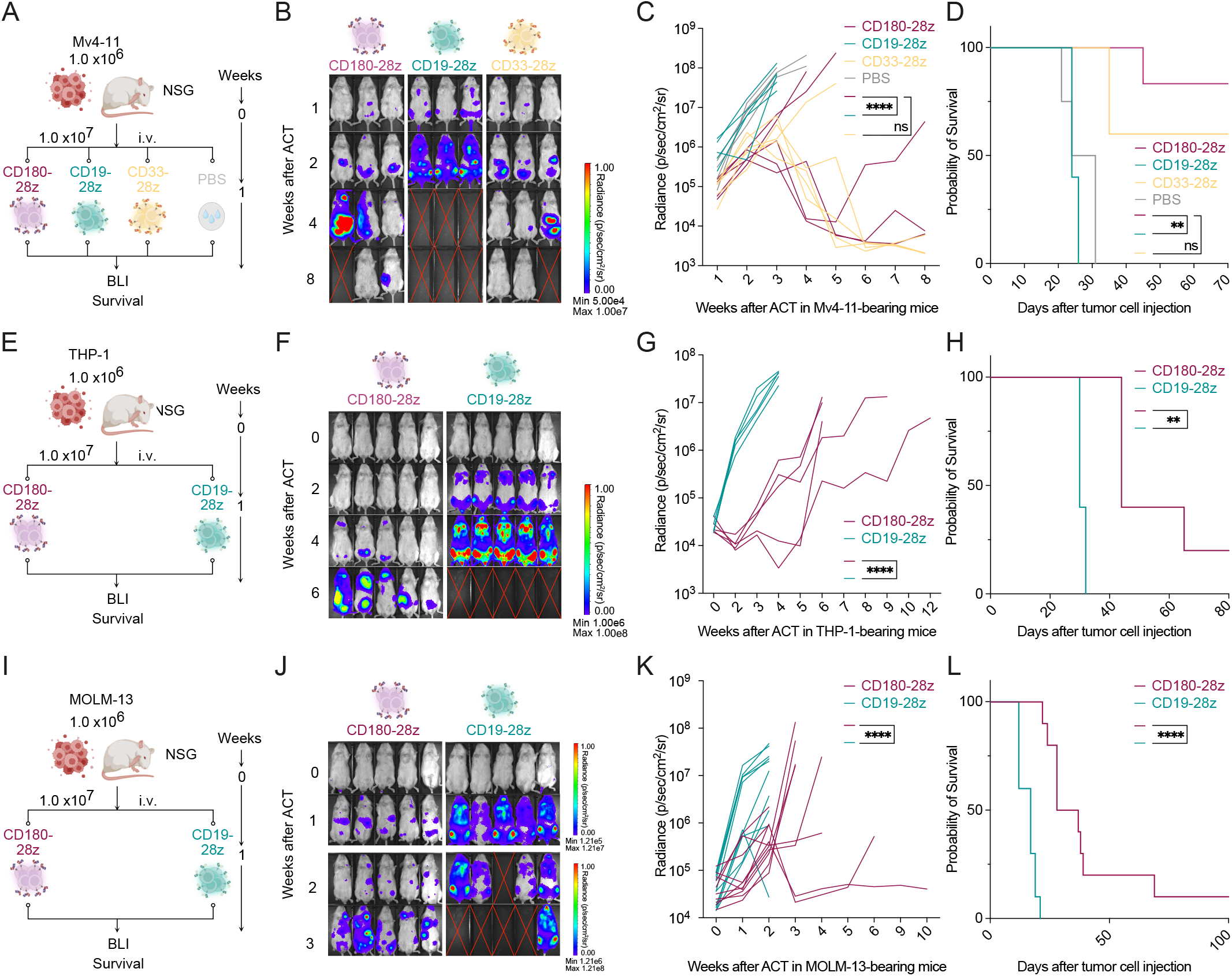
CD180-28z CAR T cells exhibit potent anti-tumor activity across multiple AML xenograft models. (**A–D**) Experimental design (**A**), representative BLI images (**B**), tumor burden quantification (**C**), and Kaplan–Meier survival curves (**D**) of NSG mice engrafted with Mv4-11 AML tumor cells and treated with the indicated number of CAR T cells (*n* = 5 mice per group). (**E–H**) Experimental design (**E**), representative BLI images (**F**), tumor burden quantification (**G**), and Kaplan–Meier survival curves (**H**) of NSG mice engrafted with THP-1 AML tumor cells and treated with the indicated number of CAR T cells (*n* = 5 mice per group). (**I–L**) Experimental design (**I**), representative BLI images (**J**), tumor burden quantification (**K**), and Kaplan–Meier survival curves (**L**) of NSG mice engrafted with MOLM-13 AML tumor cells and treated with the indicated number of CAR T cells (*n* = 10 mice per group of two independent experiment). (**A-L**) Statistical analysis by two-way ANOVA with Tukey’s multiple comparison test or log-rank test for Kaplan-Meier-Curves. For all panels: \**P* < 0.05; \*\**P* < 0.01; \*\*\**P* < 0.001; \*\*\*\**P* < 0.0001; ns, not significant.

**Extended Data Figure 12:**
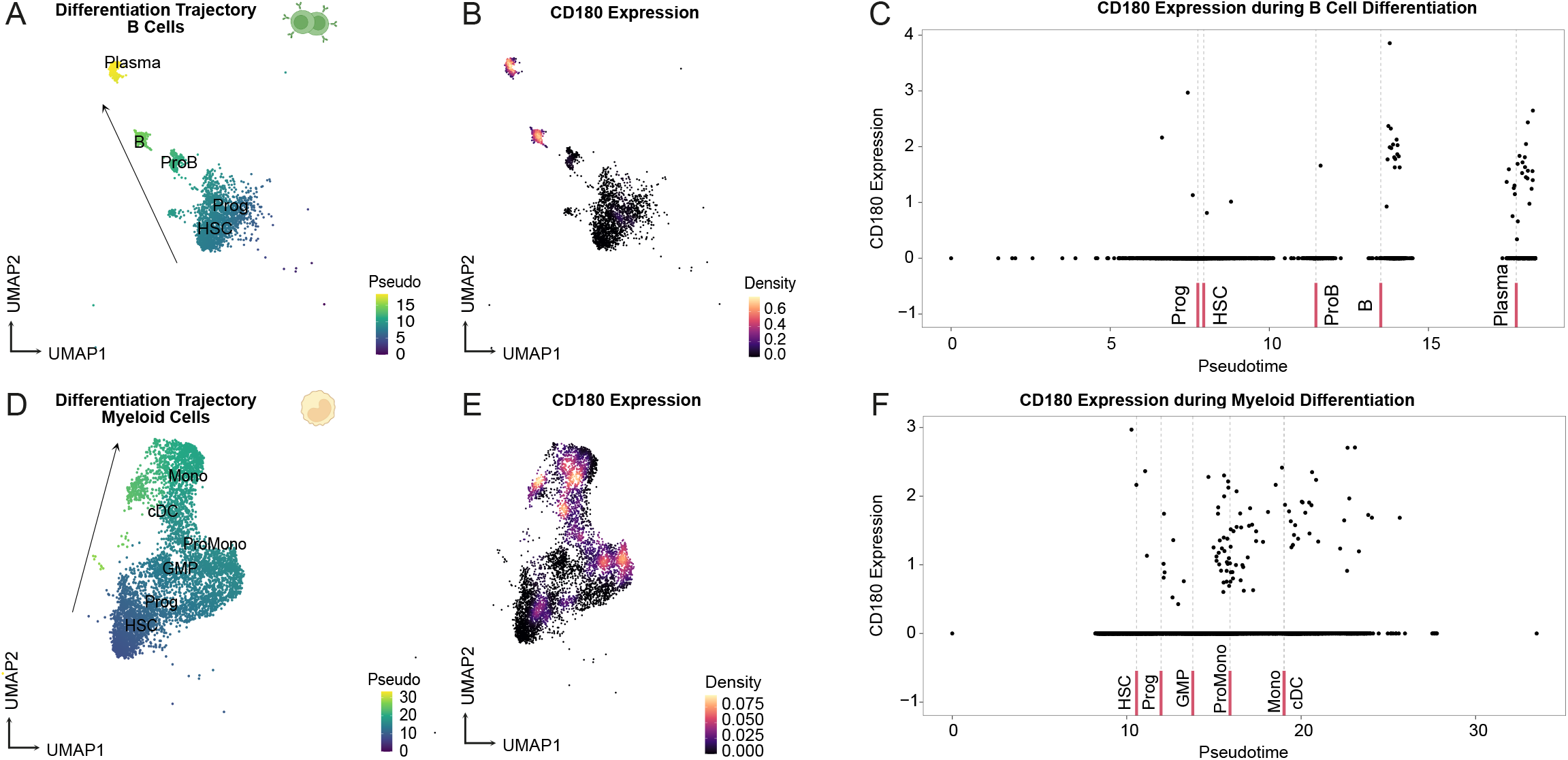
CD180 expression across hematopoietic development and safety assessment of CD180-28z CAR T cells. (**A**) B-cell development trajectory calculated using Slingshot R package in the AML scRNA-seq cohort of van Galen *et al.*^55^ (*n* = 21.667 cells). (**B, C**) CD180 expression during B-cell development visualized by UMAP (**B**) or dot plot (**C**). (**D**) Myeloid development trajectory calculated using Slingshot R package in the AML scRNA-seq cohort of Galen *et al.*^55^ (**E, F**) CD180 expression during myeloid differentiation visualized by UMAP (**E**) or dot plot (**F**).

**Extended Data Figure 13:**
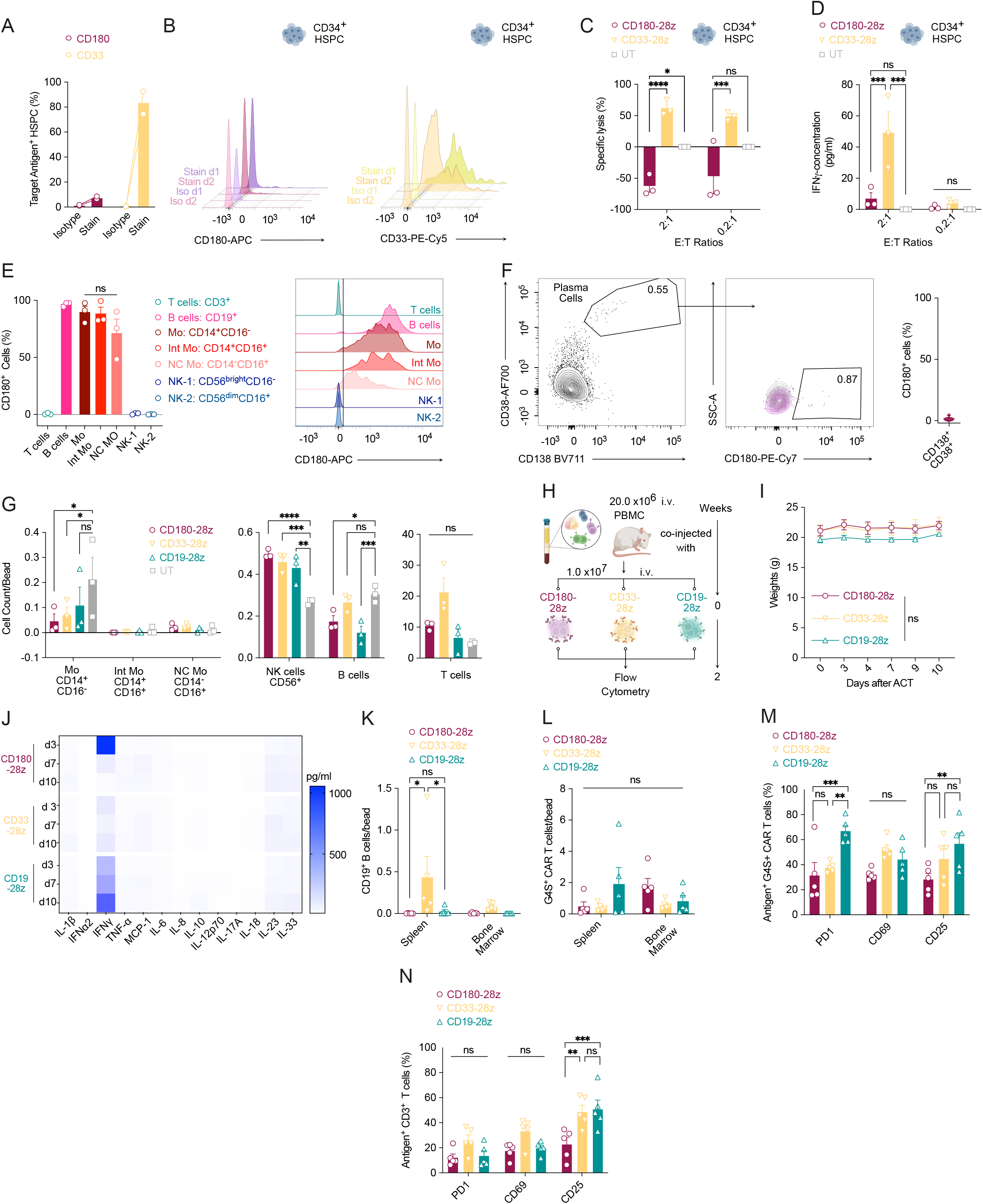
CD180 expression across hematopoietic development and safety assessment of CD180-28z CAR T cells. (**A,B**) Expression of CD180 (maroon) versus CD33 (cantaloupe) on CD34^+^ MACS-enriched healthy hematopoietic stem and progenitor cells (HSPCs) determined by flow cytometry. (**A**) Quantification of expression. (**B**) Representative flow-cytometric plots for CD180 (left) or CD33 (right). (**C,D**) Co-culture of CD180-28z, CD33-28z, or untransduced (UT) T cells with healthy HSPCs for 24 h at the indicated effector-to-target (E:T) ratios. Lysis of HSPCs (**C**) and secretion of IFNγ (**D**) were determined by flow cytometry or ELISA, respectively. Data are mean ± s.e.m. from *n* = 3 independent donors. (**E**) CD180 expression on HD-derived immune cell subsets. Left: quantification. Right: representative flow-cytometric plot. Data from n =3 independent donors. (**F**) CD180 expression on HD-derived bone marrow plasma cells. Left: gating strategy. Right: quantification. Data from n =9 different healthy bone marrow samples. (**G**) Co-culture of CD180-28z, CD33-28z, CD19-28z, or UT T cells with healthy PBMCs for 24 h at the indicated E:T ratios; PBMC lysis by flow cytometry. Data are mean ± s.e.m. from n = 3 independent donors. (**H**) Experimental design of the PBMC-humanized mouse model. (**I**) Body weight after adoptive T cell transfer (ACT). (**J**) Heatmap of serum cytokine levels (LEGENDplex) in mice treated with CD180-28z, CD19-28z, or CD33-28z CAR T cells. (**K-N**) Quantification of immune cell subsets 2 weeks after ACT. (**K,L**) B cell (**K**) and CAR T cell (**L**) counts in the indicated organs. (**M,N**) Expression of the indicated T cell markers on transferred CAR T cells (**M**) or all T cells (**N**). (**G-M**) *n* = 5 mice per group. Unless otherwise indicated, statistical significance was determined using two-way ANOVA with Tukey’s multiple comparison correction. For all panels: \**P* < 0.05; \*\**P* < 0.01; \*\*\**P* < 0.001; \*\*\*\**P* < 0.0001; ns, not significant.

## Methods

### Single cell analysis

The following open accessible datasets were retrieved through the corresponding repositories: Abbas et al.^72^ and Guo et al.^29^ (with accession number GSE198052 and GSE239721, atypical B cells characterization), van Galen et al.^55^ (GSE116256, AML characterization), Lasry et al.^28^ (GSE185381, validation for the atypical B cells differentiation state), Rade et al.^45^ (GSE234261, CAR-T cell-treatment response in the context of multiple myeloma), Yang et al.^46^ (GSE271915, CAR-T cell-treatment response in the context of multiple myeloma) and Bhagwat et al.^10^ (National Center for Biotechnology Information Database of Genotypes and Phenotypes accession number: phs001707.v4). Oesinghaus et al^74^: https://www.parsebiosciences.com/datasets/10-million-human-pbmcs-in-a-single-exp.

The filtered UMI count matrices, and, in some cases, BCR-sequencing data were used to compose the analysis object using Seurat R package (v. 4.1.0)^79^ by discarding the cells expressing fewer than 500 and more than 5000 features and/or >15% mitochondrial reads. Subsequently, the PCA was constructed using the detected variable features. To leverage the patient-and batch-related bias, the objects were integrated using the Harmony R package (Version 0.1.0)^80^. Briefly, Harmony integrates single-cell RNA-sequencing datasets by iteratively projecting cells into a shared low-dimensional space where batch effects are minimized. It performs soft clustering with a diversity-promoting penalty, computes batch-specific correction vectors within clusters, and refines these corrections over successive iterations until cells group by biological rather than technical variation.

In the next step, unbiased k-nearest neighbors and cluster identification was performed using the FindNeighbors and FindClusters functions of the Seurat R package. A two-dimensional Uniform Manifold Approximation and Projection (UMAP) was generated to visualize the cellular embedding. The Abbas et al. dataset was used as a reference for the subsequent identification of B cell subpopulations in the following datasets. For this purpose, query datasets were projected on to the Abbas et al. dataset using the symphony R package (v. 0.1.1)^81^. Briefly, Symphony builds on a harmonized reference atlas by compressing it into a minimal, portable set of parameters (e.g. PCA loadings, cluster centroids, mixture model terms) and maps query cells into the fixed reference embedding by estimating their positions via the compressed reference model. This approach facilitates scalable and reproducible cross-dataset annotation.

## Data analysis

The differential gene expression (DGE) analysis was performed by using the FindMarkers function of the Seurat R Package. The Bonferroni correction was used to adjust the results with the p-adjusted threshold of 0.05. The DGE comparisons were conducted within the clusters of interest to avoid the cell distribution-related bias. The graph-based pseudotime analysis was performed with the slingshot R package (v. 1.8.0)^82^. The CellCycleScoring function of the Seurat R package was used for cell cycle state identification. The visualization plots were constructed with ggplot2 (v. 3.3.5) and Seurat R packages.

Transcription factor (TF) regulon activity was inferred for the integrated AML/B cell atlas following library-size normalization and computation of cluster-level pseudobulk expression profiles. TF–target gene interactions were obtained from the human DoRothEA(v. 1.22.0) regulon (dorothea_hs), restricted to confidence levels A–C (the curated, high-to-medium-confidence subset), yielding a signed regulatory network in which each interaction is annotated with a mode of regulation (activating/repressing). TF activity scores were then estimated per cluster from this network using decoupleR (v. 2.16.0).

For the interaction analysis, we computed pseudobulk average expression profiles and cross-referenced these with cluster-wise differential expression results to identify literature-curated AML blast-, T cell-, and B cell-intrinsic marker genes that were both expressed above baseline and upregulated in AML relative to control within each cluster of interest. Cell–cell communication was inferred using the LIANA framework (v 0.1.14), applied separately to AML and control cells after splitting the integrated object by condition. For each condition, liana_wrap ran five complementary ligand–receptor inference methods in parallel — NATMI (edge specificity), Connectome (ligand– receptor co-expression weight), a log-fold-change–based method, SingleCellSignalR/SCA (LR interaction score), and CellPhoneDB (permutation-based mean expression) — against the curated OmniPath “Consensus” ligand–receptor resource, and results were combined per condition into a consensus ranking. Differential cell–cell communication between AML and control was quantified by pivoting the combined aggregate-rank table and calculating a log2 fold-change of aggregate rank between conditions; interactions were classified as AML-enriched, control-enriched, or shared based on a rank significance threshold of 0.05 in one or both conditions.

## Survival analyses

Bulk RNA sequencing and corresponding clinical data for adult newly diagnosed AML patients were obtained from the TCGA-LAML cohort (n = 173) via the Genomic Data Commons (GDC). Only primary tumor samples with available overall survival (OS) data were included. Gene expression values were normalized to transcripts per million (TPM) and log -transformed [log (TPM + 1)]. For samples with multiple entries, the primary aliquot was selected.

For the atypical memory B cell (B_atyp_) gene, a per-sample signature score was calculated as the mean of z-scaled log expression values of the significantly upregulated DEGs from the single-cell analysis. For single-gene analyses (*CD180, CD33, CD123, CLL-1, CD19, CD20, CD22*), normalized expression values were used directly. Patients were stratified into high, medium, or low expression groups according to interquartile range of the respective gene or signature score within the cohort.

Overall survival was defined as the time from diagnosis to death, with surviving patients censored at last follow-up. Kaplan–Meier survival curves were generated using the R packages survival and survminer, and statistical significance was assessed using two-sided log-rank tests. Hazard ratios (HR) and 95% confidence intervals were derived from Cox proportional hazards models comparing medium and high expression groups against the low expression reference. Proportional hazards assumptions were verified using Schoenfeld residuals. All analyses were performed in R (version 4.0.5) using TCGAbiolinks, GSVA, and ggplot2.

### Generation of single cell atlas for off-tumor expression screening

Single-cell atlas was generated and used as previously described^56^. All preprocessing and analysis steps of scRNA-Seq data analysis for off-tumor target screening were run in python 3 using Scanpy v.1.8.2+^83^ and anndata v.0.7.8+^84^ except stated otherwise. All scRNA-Seq figures were plotted using matplotlib and seaborn.

Single cell transcriptomic datasets were obtained either via sfaira^85^, cellxgene (https://cellxgene.cziscience.com/) or from various depositories listed in the respective original studies. A detailed description of used datasets can be found in Supplementary table 1.

For each dataset, low-quality cells were removed based on total UMI count, gene distributions and fraction of mitochondria-encoded genes per sample after visual inspection (see provided Notebook). Genes that were detected in less than 20 cells were excluded for further analysis. Normalization of UMI counts of each cell was carried out using the implemented R-based SCRAN algorithm ^86,87^. Briefly, size factors were estimated upon clustering the data using the Louvain algorithm implemented in Scanpy (tl.louvain) before running ComputeSumFactors with min.mean = 0.1. Size factors were then used for cell normalization. Finally, the data was log-transformed (log(count+1)). Cell annotation labels were provided by the authors of the respective study and were carefully inspected and harmonized between datasets upon close inspection of marker gene expression. The normalized cell x gene matrix was then used for plotting mean expression values across cell types.

### Bulk RNA sequencing

All analyses were performed in R (version 5.4.3) using the tidyverse, data.table, GEOquery, TCGAbiolinks, cBioPortalData, Biobase, org.Hs.eg.db/AnnotationDbi, hgu133plus2.db / hgu133a.db / hgu133b.db, metafor, uwot, and pheatmap/ggplot2 packages.

Publicly available AML transcriptomic datasets were obtained. Of those, The Cancer Genome Atlas AML (TCGA-LAML), BEAT-AML 2.0 (phs001657.v3.p1), GSE146173^43^, GSE232130^77^ (Leucegene), and GSE199451^78^ offered bulk RNASeq data. In addition, three array based (Affymetrix platforms) cohorts were included: GSE6891^42^, GSE37642^75^, and GSE68833^76^. For TCGA-LAML, expression matrices were assembled from per-sample transcript quantification files (log2[TPM+1]). For Beat AML, expression and mutation data were obtained as packaged study bundles via the cBioPortalData R package (cBioDataPack), which retrieves the maintained study archives; the RNA-seq assay used was mrna_seq_rpkm. GEO cohorts were retrieved with GEOquery. Clinicopathological annotation for GSE37642 and GSE146173 was supplemented with unpublished data.

#### Exclusion criteria

The following samples were excluded from all analyses: all APL cases (PML-RARA-positive / t15;17); the GSE6891 mononuclear-cell batch; Sixteen non-AML samples from Leucegene; and all leukapheresis specimens, which constitute a distinct specimen type with aberrant antigen profiles. Analyses of diagnostic disease were restricted to diagnostic specimens; relapse and remission/residual specimens were retained only for the paired diagnosis–relapse analysis.

RNA-seq raw counts were converted to log2 counts-per-million (log2 CPM), and microarray intensities were log2-transformed where required. Genes were resolved by official symbol for symbol-keyed matrices, by Ensembl gene identifier () for Ensembl-keyed matrices (GSE146173, Leucegene), and by platform-specific probe sets for the microarray cohorts; probes or transcripts mapping to the same gene were summarized by their mean. Because the cohorts were generated on different platforms, datasets were processed and analyzed independently and combined only at the level of summary statistics (see below). Each cohort underwent quality control by principal component analysis (PCA) on the top 2,000 high-variance genes. PCA was inspected for batch artefacts, specimen-type effects and biological structure (ELN risk). Several cohort-specific artefacts were identified and corrected: In GSE6891, 76 samples derived from unfractionated bone-marrow mononuclear cells (a distinct submission batch) separated strongly on PC1 and were excluded, restoring a normal variance structure on the remaining blast-enriched samples. In GSE146173, samples from two clinical trials (AMLCG-1999 and AMLCG-2008) showed a CD180 offset of +0.85 log2 that was confirmed technical rather than biological (within-trial analysis showed no corresponding disease-state effect) and was entangled with ELN risk. This was corrected by within-trial median centring of CD180. In the Leucegene SuperSeries, both sequencing instrument (HiSeq 2000 vs NovaSeq 6000) and library strandedness produced technical shifts in antigen expression. These were corrected by three-group median centring (NovaSeq, HiSeq-reverse, HiSeq-unstranded). Distributions were verified to be indistinguishable after correction. A cross-cohort uniform manifold approximation and projection (UMAP) was additionally constructed by harmonising all RNA-seq cohorts to common gene symbols, applying per-cohort per-gene z-scoring, selecting the top 2,000 high-variance genes and embedding with uwot (n_neighbors=30, min_dist=0.3). The cross-cohort UMAP showed thorough intermixing of all RNA-seq cohorts within a shared expression space with a coherent cross-cohort ELN gradient and biologically defined residual clusters (core-binding-factor and FLT3-ITD-associated AML), supporting the validity of within-cohort normalization for cross-cohort comparison.

Mutation and cytogenetic status were harmonized across cohorts with available molecular data (TCGA-LAML, BEAT-AML, GSE146173, GSE6891, GSE37642). Samples were annotated by mutational/cytogenetic subgroup (CBF-AML, *FLT3-ITD*, *RAS*, *NPM1*, MR genes, adverse cytogenetics, TP53), with harmonized mutant-versus-wild-type group labels. European LeukemiaNet 2022 (ELN-2022) risk was assigned by a rule-based classifier applied uniformly across cohorts.

Expression of CD180 was compared against five established or emerging AML immunotherapy target antigens: *CD33*, *IL3RA (CD123)*, *CLEC12A (CLL-1)*, *CD70* and *FLT3*. To compare absolute expression level across genes and cohorts, each antigen was expressed as its per-sample percentile rank against the whole transcriptome of that sample. Percentiles were pooled across all diagnostic samples and compared between antigens. Raw log2 expression per cohort was additionally shown. Within each cohort, pairwise Pearson correlations were computed among the six antigens on diagnostic samples. Per-cohort correlations were pooled across acohorts by Fisher z-transformation followed by random-effects meta-analysis; I² quantified heterogeneity. Results were displayed as a pooled correlation heatmap and as a per-cohort correlation forest plot.

#### ELN risk group

For visualization by ELN risk group CD180 was z-scored within each cohort and pooled across the cohorts with available mutation data. For quantification, the within-cohort standardized mean difference (Hedges’ g) in CD180 was computed for Adverse-vs-Favorable and Intermediate-vs-Favorable contrasts using escalc (measure=“SMD”), and pooled by random-effects meta-analysis (REML) using metafor::rma, with heterogeneity quantified by I².

#### Molecular subtypes

To further assess the relationship to molecular subtypes, the within cohort standardized mean difference (mutated − wild-type) in CD180 was computed for each of seven molecular contrasts (NPM1, FLT3-ITD, TP53, MR composite, RAS composite, CBF composite, and adverse-cytogenetics composite), and pooled by random-effects meta-analysis. Cohorts contributing fewer than three samples in either group for a given contrast were excluded from that contrast.

#### Specimen type

Within-cohort standardized mean difference (bone marrow − peripheral blood) in CD180 was computed for BEAT-AML and Leucegene on diagnostic specimens, and pooled by random-effects meta-analysis.

#### Diagnosis versus relapse

Patients with CD180 expression at both diagnosis and relapse were identified (n=21 expression-paired patients: BEAT-AML2 n=10, GSE199451 n=11). The within-patient change in CD180 (relapse − diagnosis, raw log2 within cohort) was assessed by paired Wilcoxon signed-rank test per cohort and pooled, and the per-cohort mean change was pooled by random-effects meta-analysis.

An atypical signature (*FCRL3*, *FCRL5*, *CD180*, *CD19*, *MS4A1*, *ITGAX,* **Extended Data Fig. 3A**) was scored per sample as the mean of gene-wise z-scores (row-scaled expression across samples) within each cohort. Interferon pathway activity was quantified using response signatures rather than the interferon ligands themselves. A type-I IFN response signature (interferon-stimulated genes, *ISG; IFI27, IFI44, IFI44L, IFIT1, IFIT3, ISG15, RSAD2, SIGLEC1, USP18, MX1, MX2, OAS1, OAS2, OAS3, IFI6, LY6E, HERC5, SPATS2L*) and a type-II IFN (IFN-γ) response signature (*CXCL9, CXCL10, CXCL11, GBP1, GBP4, GBP5, IDO1, IRF1, CIITA, HLA-DRA, TAP1, PSMB9, UBD, SOCS1, WARS1, ANKRD22*), as well as individual candidate ligands (e.g., *TNFSF13B/BAFF, CD70, CD40LG, IL15, IL1A, IL1B, IL7*), were scored analogously (**Fig. 5D**).

#### Correlation and meta-analysis

For each molecular feature and subgroup defined above, the standardized mean difference (Cohen’s d, mutant − wild-type) was calculated, and group differences were tested by two-sided Wilcoxon rank-sum tests with Benjamini–Hochberg correction. Results were visualized per cohort and as combined heatmaps.

#### AML cell-line expression

Constitutive expression of candidate ligands in AML cell lines was analyzed using bulk RNA-seq data from the Cancer Cell Line Encyclopedia/DepMap (log2(TPM+1)) accessed via the Bioconductor depmap package. A panel of AML lines was compared with the B-ALL line Nalm-6 as a contrast. Type-I IFN genes were additionally aggregated into a per-line module score (log2 of summed TPM). Differential expression between AML lines and Nalm-6 was expressed as the log2 fold-change of mean log2(TPM+1) values.

### Spatial Transcriptomic

Publicly available spatial transcriptomics data from the bone marrow and extramedullary AML cohort of Dasdemir et al. were used for this analysis^47^.

Spatially resolved transcriptomes were generated using the 10x Genomics Visium platform (standard and CytAssist chemistries) for bone marrow (BM) and extramedullary (EM) tumor sections from two AML patients, yielding six slides in total; one low-quality slide (EM2, v1 chemistry) was excluded prior to downstream analysis, leaving five slides for the main pipeline. Spots with zero total counts were removed, and the mitochondrial read fraction was calculated per spot. Quality thresholds were defined per sample using an adaptive, log-scale median-absolute-deviation (MAD) approach: spots below (median − 3×MAD) UMI counts or detected features were excluded, with absolute floors of 50 counts and 25 features as a backstop; spots exceeding (median + 3×MAD) mitochondrial content were additionally removed on a per-sample basis.

Filtered spots were normalized with SCTransform (Spatial assay), and integration features were selected with SelectIntegrationFeatures (3,000 features), restricted to genes with fitted SCT residuals present in every sample. Samples were normalized (NormalizeData, log-normalization) and integrated by canonical correlation analysis (CCA-based FindIntegrationAnchors/IntegrateData).

Composite gene-signature scores for the atypical memory B cell (B_atyp_; *FCRL3, FCRL5, CD180, CD19, MS4A1, ITGAX*) and T cell (*IL10, HAVCR2, GRB2, PTPN2, CD3E*) programs were computed per spot using AddModuleScore on the Spatial assay, and spatial distributions of individual markers and composite scores were visualized with SpatialFeaturePlot.

To assess spatial co-localization, for each signature the top decile (90th percentile) of spots by score within each slide was defined as “positive” for that signature. Pairwise nearest-neighbor Euclidean distances (on Visium tissue pixel coordinates, GetTissueCoordinates) from each positive spot of one signature to the nearest positive spot of a second signature were calculated, and compared against a null distribution generated by 100 permutations of randomly drawn, size-matched spot sets within the same slide.

### Cell lines

Human AML cell lines MOLM-13, Mv4-11, THP-1 and OCI-AML3 were purchased from ATCC (USA). Cells were transduced with a pCDH-EF1a-eFly-eGFP plasmid^88^ purchased from addgene (plasmid #104834, deposited by I. Jeremias) as recently described^6^. Enhanced green-fluorescent protein (eGFP) positive cells were sorted with a BD FACSAria™ III Cell Sorter.

Short tandem repeat (STR) typing was used to verify the cellular origin of all cell lines used . Cells were regularly tested for mycoplasma contamination by polymerase chain reaction (PCR) as previously described^89^. Cells were cultured as previously described in RPMI containing 10 to 20 % FBS, 2 mM L-glutamine, 100 U/ml penicillin and 100 µg/ml streptomycin. All cells were grown at 37°C in a humidified incubator with 5 % CO_2_. Cells were stored at -80°C or in liquid nitrogen after addition of freezing medium (90 % FCS and 10 % DMSO).

### Animal experiments

Animal experiments were approved by our local regulatory agency (Regierung von Oberbayern, ROB) and carried out according to the regulations set by the regulatory agency. C57Bl/6 and NOD.Cg-Prkdc^scid^ Il2rg^tm1WjI/SzJ^ (NSG) or NOD-*Prkdc^scid^-IL2rg^Tm^*^1^/Rj were purchased from Charles River Laboratories (Sulzfeld, Germany), Janvier (St Berthevin, France), or bred at our local facilities. Mice were housed in specific-pathogen free facilities.

For AML models, human AML cell lines MOLM-13, MV4-11, THP-1 and OCI-AML3 or murine C1498 AML cells were injected intravenously (i.v.) at the indicated cell numbers. Tumor progression was monitored using the bioluminescence imaging (BLI) platform Lumina X5 (PerkinElmer, USA) as recently described^6,90^. T and B cells were injected i.v. at the indicated cell numbers and time points.

The AML PDX models have been extensively characterized^91^ and were kindly provided by I. Jeremias. Adoptive T cell transfer was carried out at the indicated cell numbers and time points. BLI images displayed in the figures are representative images. Red cross indicates that mice were removed from the experiment due to reaching the pre-defined abortion criteria set by our local regulatory agency as a humane surrogate parameter for survival.

### AML blast isolation and culture

The use of primary AML blasts were approved by the Institutional Review Board of the Ludwig-Maximilians-Universität (LMU) or the Ethics Committee of the Medical Association of Hamburg after written informed consent. Isolation of AML blasts and primary AML cultures have been extensively described in prior publications ^6,90,92,93^.

### Flow cytometry

Flow cytometric analysis was performed using a LSRFortessa™ II, a Beckman Coulter CytoFLEX LX or a Cytek™ Aurora spectral analyzer. Flow cytometric analyses were performed as recently described^6,89^. In short, cells were harvested, centrifuged at 200 to 400 g for 5 min at 4°C and then washed twice with PBS. Cells were then incubated with human TrueStain FcX™ (Biolegend, USA) and stained with a fixable viability dye (eFluor™ 780, eBioscience, USA) for at least 10 min at room temperature or 4°C. After two additional washing steps, cells were stained with the respective antibodies for 30 min at 4°C. Prior to analyses, cells were again washed with PBS.

Cells from the flow cytometry discovery cohort were stained with anti-human PD-L2 (clone 24F.10C12), anti-human CD27 (clone O323), anti-human CD138 (clone MI15), anti-human CD86 (clone IT2.2), anti-human CD24 (clone ML5), anti-human IgD (clone IA6-2), anti-human PD-1 (clone A17188B), anti-human CD180 (clone MHR73-11), anti-human BTLA (clone MIH26), anti-human FcRL5 (clone 509f6), anti-human CD38 (clone HB-7), anti-human HVEM (clone 122), anti-human CD20 (clone 2H7) and anti-human PD-L1 (clone MIH1).

The cells from the flow cytometry validation cohort were stained with anti-human CD45 (clone HI30), anti-human CD19 (clone HIB19), anti-human CD20 (clone S8015E), anti-human CD27 (clone M-T271), anti-human IgD (clone IA6-2), anti-human HVEM (clone 122), anti-human CD86 (clone IT2.2), anti-human PD-1 (clone EH12.2H7), anti-human PD-L1 (clone 29E.2A3), anti-human PD-L2 (clone 24F.10C12), anti-human CD180 (clone MHR73-11) and anti-human CD33 (clone WM53).

For B cell polarization cells were stained with anti-human CD19 (clone HIB19, BV785), anti-human CD45 (clone 2D1, PerCP/Cy5.5), anti-human CD38 (clone HB-7, BV605), anti-human CD11c (clone Bu15, BV711), anti-human IgD (clone IA6-2, BV510), anti-human CD20 (clone 2H7, APC), anti-human CD27 (clone O323, AF700 or clone LG3A10, BV421), anti-human CD274/PD-L1 (clone 29E.2A3, PE/Cy7) and anti-human CD180/RP105 (clone MHR73-11, PE or PE/Cy7) and anti-human CD307e/FcRL5 (clone 509f6, eFluor660; eBioscience).

Primary AML blasts were stained with following antibodies: anti-human CD45 (clone HI30), anti-human CD33 (clone P67.6; WM53, Invitrogen/eBioscience), anti-human CD34 (clone 561), anti-CD38 (clone HB-7) and anti-human CD180 (clone MHR73-11) or mouse IgG1 control (clone MOPC-21).

Transduction efficiencies were determined by staining with anti-c-myc-FITC (clone SH1-26E7.1.6; Miltenyi Biotec). Activation of T cells was determined by measurement of the early activation marker CD69 (clone FN50) or PD-1 (clone EH12.2H7).

For intracellular staining, Brefeldin A was added for the final 4h of culture. Cells were fixed and permeabilized with the eBioscience™ Foxp3/Transcription Factor Fixation/Permeabilization kit according to the manufacturer’s instructions and stained with anti-T-bet (clone 4B10, PE or PE/Cy7), anti-human Bcl-2 (clone 100, BV421) and anti-human TGF-β1 (clone TW4-9E7, PE-CF594; BD Horizon).

For phospho-flow cytometry, cells were fixed directly in the culture plate by adding an equal volume of 4% PFA, washed and resuspended in ice-cold 96% methanol (BD Phosflow™), and stained with anti-pSTAT1 (Tyr701, AF488; Cell Signaling Technology).

CAR surface expression was detected with anti-c-myc (clone SH1-26E7.1.6; Miltenyi Biotec) or an anti-G4S linker antibody (clone E702V). Target antigen expression on CAR T cell targets was assessed with anti-human CD180 (clone MHR73-11), anti-human CD33 (clone P67.6) and anti-human CD19 (clone HIB19), each with the corresponding isotype control.

For analyses of HSPC and of PBMC, cells were stained with anti-human CD34 (clone 561), anti-human CD38 (clone HB-7), anti-human CD138 (clone MI15), anti-human CD3 (clone OKT3, UCHT1, HIT3a), anti-human CD19 (clone HIB19), anti-human CD14 (clone M5E2), anti-human CD16 (clone 3G8), anti-human CD56 (clone HCD56), anti-human CD33 (clone P67.6) and anti-human CD180 (clone MHR73-11). All antibodies were purchased from Biolegend, USA, if not otherwise indicated.

### Multiplexed Immunofluorescence (mIF)

Multiplex immunofluorescence was performed on formalin-fixed, paraffin-embedded (FFPE) bone-marrow trephine biopsy sections from n = 5 patients with AML. Biopsies were fixed in 4% formalin, decalcified using [EDTA-based decalcification 10% Roth, for 8 hours], and paraffin-embedded. Staining was carried out using the Opal multiplex IHC system (Akoya Biosciences). Sections were dewaxed and rehydrated, followed by heat-induced epitope retrieval. Each staining cycle consisted of endogenous peroxidase blocking, primary antibody incubation, an HRP-conjugated secondary antibody, and tyramide signal amplification with the assigned Opal fluorophore. A heat-mediated stripping step between cycles simultaneously served as epitope retrieval for the subsequent marker. This cycle was repeated sequentially for the full panel (CD180, CD20, CD3, PD-L1, and CD34), and nuclei were counterstained with spectral DAPI. The following antibodies were used: CD180 (EPR14720, abcam), dilution 1:100, Opal 620; CD20 (L26, Leica), dilution 1:150, Opal 520; CD3 (LN10, Leica), dilution 1:200, Opal 570; PD-L1 (ERP20529, abcam), 1:100, Opal 690 and CD34 (QBEnd, Leica), 1:100, Opal 780.

Whole-slide multispectral images were acquired on a Vectra Polaris imaging system (Akoya Biosciences) and spectrally unmixed in inForm software (Akoya Biosciences) using a spectral library and an autofluorescence reference; the unmixed single-channel images were exported as TIFF files.

For spatial analysis, cell segmentation and phenotyping were performed in Cellprofiler (v4.2.8). Nuclei were segmented from the DAPI channel (Minimum Cross-Entropy thresholding) and cell boundaries were evaluated by CD20 and CD3. Each cell was classified as a B cell or a T cell on the basis of mean CD20 and CD3 intensity, respectively, using fixed thresholds. For each T cell, the distance to the nearest B cell was then determined, and inter-cell distances were converted from pixels to micrometres using the image pixel size. Results are reported as the median] nearest-neighbour distance (µm) per sample.

### iPSC cell line and maintenance

The human iPSC line (HMGU1, hPSCreg IFSi001-A) was provided by the iPSC Core Facility, Institute of Stem Cell Research, Helmholtz Center Munich. Human iPSCs were cultured under feeder-free conditions on Geltrex-coated (Gibco) plates in mTeSRplus medium (STEMCELL Technologies), with medium changes every other day, passaging with ReLeSR (STEMCELL Technologies) every 4– 5 days, and routine testing for mycoplasma infections.

### Bone marrow organoid differentiation

Bone marrow organoids (BMO) were differentiated as previously described by Frenz-Wiessner et. Al ^44^. Briefly, on day -3, iPSCs were dissociated into single cells using Accutase (Gibco), collected, and resuspended in aggregation medium (KnockOut DMEM/F12 supplemented with KnockOut Serum Replacement, penicillin-streptomycin L-glutamine, ß-mercaptoethanol, and nonessential amino acids) containing ROCK inhibitor to form embryoid bodies (EBs). After 24 hours, the medium was replaced with fresh aggregation medium without ROCK inhibitor, and mesoderm induction was initiated on day 0 using mTeSRplus with BMP4, CHIR99021, and VEGF. All cytokines were purchased from StemCell Technologies. On day 2, the medium was switched to Essential 6 (Gibco) with VEGF, bFGF, SCF, and SB431542. On day 4, EBs were embedded in a collagen I (ibidi)-Matrigel (Corning) mixture for further development in Stempro-34 medium (Gibco) with VEGF, bFGF, SCF, and SB431542. The embedded EBs were then cultured in Stempro medium containing IL-3, Flt-3L, TPO, VEGF, and SCF, with medium changes on days 6, 8, and 10. On day 10, sprouted EBs were extracted from the 3D matrix and transferred to a low-attachment plate for continued culture in a cytokine-supplemented medium, with half the medium refreshed every 3–4 days until full maturation by day 17. BMOs were subsequently used for co-culture experiments.

### Bone marrow organoid, B cell and AML cell co-culture

For co-culture experiments, BMO were generated as described above. B cells were isolated from healthy donor PBMCs with a B cell isolation kit (Miltenyi Biotec, Germany) and labeled with CellTrace™ Violet Cell proliferation dye according to manufacturer’s instruction. The experiment was performed at an B cell to AML ratio of either 10:1 or 1:1. Either 50,000 B-cells and 5,000 GFP-expressing OCl-AML3 cells (10:1) or 10,000 B cells and 10,000 GFP-expressing OCl-AML3 cells (1:1) were added to each BMO and co-cultured in StemPro medium supplemented with IL-3, Flt-3L, TPO, VEGF, and SCF for 7 days. After 7 days, BMOs were collected and either analyzed with confocal microscopy (see below) or enzymatically dissociated using a mixture of Dispase II, Liberase TL and DNAse at 37°C for 25 minutes. Following incubation, single cell suspensions were washed with protein-containing FACS buffer and then analyzed by flow cytometry as described above.

### Immunofluorescent staining of whole BMOs

BMOs were fixed with 4 % paraformaldehyde for 1 hour at room temperature (RT). After fixation, they were permeabilized in a blocking buffer (3 % FBS, 1 % BSA, 0.5 % Triton X-100, 0.5 % Tween 20, 0.01 % sodium deoxycholate) for 3 hours at RT on a shaker. The BMOs were then incubated overnight at 4°C with a primary antibody (anti-NGFR/CD271 Sigma, catalog no. HPA004765, 1:500) or anti-Nestin (R&D Systems, catalog no. MAB1259, 1:20) in blocking buffer. After three 10-minute washes with PBST (0.05 % Tween in PBS), they were incubated with a secondary antibody (AlexaFluor 568, 1:300) for 2 hours at RT. The BMOs were washed three more times with PBST, mounted in a 1.0 mm iSpacer on a 22×50mm no. 1.5 coverslip, sealed with double-sided tape, and stored in the dark at 4°C until imaging.

### Confocal microscopy

Confocal laser scanning microscopy (CLSM) of whole organoids after engraftment was conducted using an inverted Zeiss LSM800 microscope equipped with a x20/0.8 M27 air objective (Zeiss). Tile scans were acquired to cover the entire organoid in the x-y plane, with z-stacks spanning a depth of 50μm. To capture images, spectral channels were taken in sequence and adjusted to optimally excite the fluorophores, while minimizing bleed-through and background noise. Images were acquired using ZEN blue software (v.2.6). Subsequent image processing was done in Imaris (v.10.0).

### Suppression Assay

Peripheral blood mononuclear cells (PBMCs) were isolated from whole blood using a ficoll density gradient centrifugation. In some assays, bone marrow mononuclear cells (BMMC) were used. T cells and B cells were enriched from the PBMCs or BMMCs using a Pan T cell isolation kit (Miltenyi Biotec, Germany) and a B cell isolation kit (Miltenyi Biotec, Germany). B cell subsets were further differentiated using CD27 Microbeads (Miltenyi Biotec, Germany) and IgD Microbeads (Miltenyi Biotec, Germany).

To assess cellular proliferation, T cells were labelled with the CellTrace™ Far Red Cell Proliferation Kit (Thermo Fisher Scientific, USA) in accordance with the manufacturer’s instructions. Dye dilution was analyzed by flow cytometry on day 5. Moreover, the T cells were stimulated with human T-Activator CD3/CD28 Dynabeads^®^ (Life Technologies, Germany).

Where indicated, the anti-PD-1 antibody nivolumab or the anti-BTLA antibody tifcemalimab (MedChemExpress) was added at 1 µg/ml to assess reversal of B cell-mediated suppression.

To assess suppression of CAR T cell function, healthy-donor B cells were polarized with primary AML blasts (pAML) for 5 to 7 days and then co-cultured with indicated CAR T cells or untransduced (UT) T cells and pAML blasts as targets for 48 h at the indicated effector-to-target ratio. Non-polarized B cells and CAR/UT T cells without B cells served as controls. Successful polarization was confirmed by PD-L1 upregulation, and CAR T cell–mediated lysis of pAML blasts as well as T cell counts were quantified by flow cytometry.

### B cell polarization experiments

CD19 B cells were isolated from healthy-donor PBMCs using CD19 MicroBeads (Miltenyi Biotec) or B Cell Isolation Kit II per the manufacturer’s instructions. For cell-line co-cultures, B cells were cultured with AML lines (OCI-AML3, MOLM-13, PL-21) or, as a control, the B-ALL line Nalm-6 at a 1:1 ratio for 7–14 days. For primary co-cultures, B cells were co-incubated with primary AML blasts (pAML) or bone-marrow plasma from patients with AML or ALL on irradiated (20 Gy) MS-5 stromal cells in primary blast medium (see above), for up to 7–10 days; B cells on MS-5 alone served as a control.

For supernatant stimulation, cell-free supernatants were generated from co-cultures of CD33-28z or CD19-28z (control) CAR T cells with OCI-AML3 cells, alongside single-cell-type (AML alone, CAR T cells alone) and medium controls, and pooled 1:1 from two CAR T cell donors. B cells were plated at 100,000 cells per well in a 96-well plate in B-cell medium (IMDM, 10% human serum, penicillin/streptomycin, L-glutamine) and stimulated 1:1 with the respective supernatant (125 µl final volume). For pathway inhibition, B cells were pre-incubated (30–60 min) with the TLR4 inhibitor CLI-095 (TAK-242; 1 and 0.1 µM), the TLR7/9 antagonist ODN 2088 (1 and 0.1 µM), the TLR9 antagonist ODN INH-18 (5 and 0.5 µM), or the JAK1/2 inhibitor ruxolitinib (300 nM), with DMSO as vehicle control, before supernatant addition.

For cytokine and agonist screening, B cells were plated at 1 × 10 cells per well and stimulated for 72 h with recombinant IFN-α (100 ng/mL), IFN-γ (100 ng/mL), IL-1β (100 ng/mL), IL-18 (100 ng/mL), CpG ODN (0.1 µM), R848 (1 µg/ml ), soluble CD40L (100 ng/ml) or plate-bound CD70/CD27L (500 ng/mL, coated overnight), alone or in the indicated combinations. Tumour cell lysates were generated from OCI-AML3 and NALM-6 cells. Across all assays, recovered B cells were quantified and phenotyped by flow cytometry at the indicated timepoint.

### Generation of CAR constructs

Anti-human CD180 single chain fragment variable (scFv) was obtained from US patent Number: US 9,416,188 B2. CAR constructs were designed *in silico* and provided by commercial vendors (TwistBioscience, USA). The different CAR configurations are depicted in **Figure 6A and Extended Data Figure 10A.** Anti-CD19 and anti-CD33 CAR T cells have been previously described^6^. All CAR constructs included a myc tag to enable CAR detection.

### Retroviral human T cell transduction

Retroviral transduction was carried out using established protocols, comprehensively described _elsewhere_ 89,90,94,95.

### Co-cultures, cytokine measurements and proliferation assays

For all co-culture experiments, all different CAR T cell conditions were adjusted to similar transduction efficiencies with untransduced T cells (UT). Between 30.000 and 50.000 tumor cells, transduced with the above described luciferase construct, were plated into a flat-bottom 96-well plate. Indicated amounts of CAR T cells or control T cells were then added to the plate and incubated for 48 to 72 hours, respectively. After two or three days, plates were centrifuged and the supernatant was collected. Supernatant was analyzed using BD Bioscience IFN-γ ELISA kit, according to manufacturer’s instructions. Cell pellet was then transferred into a new 96-well plate. Killing of tumor cells was measured using BioGlo Luciferase Assay System according to manufacturer’s instructions or with flow cytometry.

For proliferation assays, CAR T cells were stained with a CellTrace™ Far Red Cell Proliferation Kit (Thermo Fisher Scientific, USA) prior to co-culturing with tumor cells. T cell proliferation was quantified via flow cytometry after 7 days of co-culture.

### Hematopoietic stem and progenitor cell (HSPC) assay

HSPC assay were performed as previously described. In short, enriched bone marrow-derived CD34^+^ HSC were purchased from Stem Cell Technologies and expanded for 7 days using complete StemSpan II medium (Stemcell Techologies) supplemented with UM729 small molecule inhibitor to preserve an immature phenotype^96^. After expansion, HSPC were collected, counted and co-cultured with CAR T cells as described above. Killing of HSPC was then quantified by flow cytometry.

### Software and statistical analysis

Flow cytometric analysis was performed with FlowJo V10.3 to V.10.9.0. BLI images were analysed using Living Image 4.4 (PerkinElmer, USA). Statistical analyses were performed with GraphPad Prism software V9.3.1 to V10.0.1 (San Diego, CA, USA).

## Data and code availability

Python and R scripts for replicating the figures from the scRNA-seq data will be uploaded to a GitHub repository upon publication of the manuscript.

The transcriptomic single-cell atlas used within this study uses published data sets and can be accessed through the respective studies 27 to 50 and scripts can be accessed via https://gitfront.io/r/MThomas/hXvhiqhnRjjy/HL-off-tumor-antigen-expression/.

All other scripts can be accessed through the following link, which will be made publicly available upon publication: https://gitfront.io/r/vlkavaka/CBe2oAqd5v3x/atypical-b-CD180/.

## Material availability

All raw data generated in this study will be uploaded to the LMU open data repository (https://data.ub.uni-muenchen.de) and can be freely accessed in accordance with the principles of open science of the European Research Council (ERC). All materials will be made available upon reasonable request to the lead authors Dr. Adrian Gottschlich and Prof. Dr. Sebastian Kobold, pending the execution of an institutional material transfer agreement.

